# Social and genetic diversity among the first farmers of Central Europe

**DOI:** 10.1101/2023.07.07.548126

**Authors:** Pere Gelabert, Penny Bickle, Daniela Hofmann, Maria Teschler-Nicola, Alexandra Anders, Xin Huang, Iñigo Olalde, Romain Fournier, Harald Ringbauer, Ali Akbari, Olivia Cheronet, Iosif Lazaridis, Nasreen Broomandkhoshbacht, Daniel M. Fernandes, Katharina Buttinger, Kim Callan, Francesca Candilio, Guillermo Bravo, Elizabeth Curtis, Matthew Ferry, Denise Keating, Suzanne Freilich, Aisling Kearns, Éadaoin Harney, Ann Marie Lawson, Kirsten Mandl, Megan Michel, Victoria Oberreiter, Jonas Oppenheimer, Susanna Sawyer, Constanze Schattke, Kadir Toykan Ozdogan, Michelle Hämmerle, Lijun Qiu, Noah Workman, Fatma Zalzala, Swapan Mallick, Matthew Mah, Adam Micco, Franz Pieler, Juraj Pavuk, Catalin Lazar, Tibor Paluch, Maja Krznarić Škrivanko, Mario Šlaus, Željka Bedić, Friederike Novotny, László D. Szabó, Orsolya Cserpák-Laczi, Tamara Hága, Zsigmond Hajdú, Pavel Mirea, Emese Gyöngyvér Nagy, Zsuzsanna M. Virág, Attila M. Horváth, László András Horváth, Katalin T. Biró, László Domboróczki, Tamás Szeniczey, János Jakucs, Márta Szelekovszky, Farkas Zoltán, Sándor Sztáncsuj, Krisztián Tóth, Piroska Csengeri, Ildikó Pap, Róbert Patay, Anđelka Putica, Branislav Vasov, Bálint Havasi, Katalin Sebők, Pál Raczky, Gabriella Lovász, Zdeněk Tvrdý, Nadin Rohland, Mario Novak, Matej Ruttkay, Dusan Boric, János Dani, Martin Kuhlwilm, Pier Francesco Palamara, Tamás Hajdu, Ron Pinhasi, David Reich

## Abstract

The Linearbandkeramik (LBK) Neolithic communities were the first to spread farming across large parts of central Europe, settling fertile regions from Ukraine to France during the second half of the 6th millennium BCE. The LBK had a high degree of material culture uniformity, albeit with regional differences in settlement patterns, subsistence, and mortuary practices. To date, ancient DNA data from LBK individuals have been generated for a limited number of locations and often in small sample sizes, making it challenging to study variation within and across sites. We report genome-wide data for 178 LBK individuals, from the Alföld Linearbankeramik Culture (ALPC) eastern LBK site of Polgár-Ferenci-hát in Hungary, the western LBK site of Nitra in Slovakia, and the enclosed western LBK settlement and massacre site of Schletz in Austria, as well as 42 LBK individuals from 18 other sites. We also report genome-wide data for 28 Early Neolithic Körös and Starčevo individuals from 13 sites, viewed as the predecessors of the LBK. We observe a higher percentage of western hunter-gatherer (WHG) admixture among individuals in the eastern LBK than in the far more widely distributed western LBK, showing that these two archaeologically distinct cultures also had different genetic trajectories. Most WHG-farmer mixture occurred just before the dawn of the LBK culture and there is no evidence that the WHG ancestry came systematically more from males or females. However, we do find strong genetic evidence for patrilocality among the LBK, extending previous findings based on isotopic analysis, with more genetic structure across sites on the male than on the female line, and a higher rate of within-site relatives for males. At Schletz we detect almost no first-degree relatives despite reporting data from almost every skeleton present at the site, showing that this massacre involved people from a large population, not a small community.

## Introduction

The archaeological roots of the Linear Pottery culture (Linearbandkeramik, LBK) ca. 5500-5000 BCE are conventionally traced to the Starčevo culture of central Transdanubia ^1–3^, and the Körös culture of the eastern Carpathian Basin ^4^. The LBK is often divided into two subgroups: the ’eastern LBK’ Alföld Linearbankeramik Culture (ALPC) limited to the east of the Carpathian Basin, and the expansive ’western LBK’. The western LBK spread in two waves, first from Transdanubia, at ca. 5500 BCE, to Slovakia, Austria, Moravia, Bohemia, and central and eastern Germany. Several centuries later, a second wave reached the Paris basin and adjacent areas of France as far west as Normandy and as far east as Poland, Ukraine, Moldova, and Romania ^1,2^.

LBK material culture appears strikingly uniform, given its geographic extent, with the typical LBK settlement pattern consisting of clusters of sites along the alluvial plains of rivers. Nevertheless, archaeological studies^5^ have documented subtle differences in subsistence, settlement patterns, health, and lifeways among LBK communities, identifying three sociocultural regions: the great Hungarian plain and Transdanubia, central Europe, and the Paris basin^6^. The LBK culture disappeared around 5000-4900 BCE. Studies of the temporal distributions of radiocarbon dates suggest a demographic collapse in that century ^7^, potentially linked with violence exemplified at the Late LBK massacres sites of Talheim and Schletz ^8,9^.

Studies of variation in strontium isotope ratio have provided insight into mobility patterns in the LBK, notably at Nitra, Schwetzingen, and Vedrovice. These analyses revealed higher variability in strontium isotopic ratios in females than males^5^, suggesting women were more likely than men to have originated from outside the communities where they were buried, pointing to patrilocal practices. Further evidence for patrilocality came from a study showing that males buried with polished stone adzes, likely indicative of high social status, had less strontium variation than males without them, suggesting that the former tended to be born and live in their natal communities ^10^. In the archaeological context of settlement layouts, these results suggest that LBK society may have been organized into patrilocal clan-like groups ^11^, with land inherited on the male line. Most LBK sites are located on loess soils, and subsequently, movements of individuals within loess regions are not easily detectable based on isotope ratios. Paleogenomic methods have the potential to reveal differences between male and female behaviours regardless of local geology. There have been new discussions about this problem ^12^.

Analyses of whole genome data from 157 LBK individuals published before this study showed that they inherited their predominant ancestry from Early European farmers (EEF) who then mixed with local European Mesolithic populations, resulting in admixed groups with typically 5% western hunter-gatherer (WHG) ancestry ^13–18^, with a possible differential contribution of Starčevo to LBK and Körös to the ALPC ^19^. However, some LBK individuals have a much higher percentage of WHG ancestry ^17^, suggesting a more complex admixture process ^15^. The only published cemetery-scale studies of LBK substructure focused on the western LBK sites of Derenburg-Meerenstieg II and Stuttgart-Mühlhausenin both in Germany, both with homogenous ancestry ^15,20^.

We present a fine-grained analysis of intra- and inter-site variation at three locations with different archaeological characteristics: A) The ALPC settlement site of Polgár-Ferenci-hát (5500-5100 BCE) in eastern Hungary in which individuals were buried between houses rather than in a cemetery, B) The cemetery of Nitra, western Slovakia, which corresponds to the LBK expansion phase from 5200–5000 BCE, and C) the enclosed settlement and massacre site of Schletz in Lower Austria dated to the Late LBK around 5000 BCE. We co-analyzed these data with newly reported data from 31 other archaeological sites and previously published data to address the following topics: 1) the extent of genetic differentiation between the LBK and ALPC; 2) kinship patterns of LBK communities and the extent of their correlation to variations in isotopic values and grave goods; 3) correlations between kinship and differences in diet and mobility (which have previously been hypothesized to be related to LBK social structure); and 4) the genetic structure of the individuals massacred at Schletz.

## Results

We generated genome-wide data passing standard metrics for authentic ancient DNA data from 248 individuals of the Starčevo, Körös, and LBK/ALPC cultures from Austria, Slovakia, Croatia, Romania, Serbia, and Hungary, using enrichment for 1.24 million single nucleotide polymorphisms (SNPs) (Figure 1A, Supplementary Tables 1-2). The new data include 19 Starčevo, 9 Körös (pre-LBK), 67 Hungarian ALPC (henceforth “Hungary_ALPC”), 3 Hungarian LBK (“Hungary_LBK”), 87 Austrian LBK (“Austria_LBK”) and 63 Slovakian LBK (“Slovakia_LBK”) from a total of 34 archaeological sites (Figure 1A). Individuals with fewer than 30,000 SNPs covering the autosomal targets were not included in ancestry analyses, but their data are reported. In addition, we did not use data from 1st-degree relatives of higher coverage individuals in the data set for ancestry analyses. We co-analyzed these individuals with published data for 172 Starčevo, Körös, ALPC, and LBK individuals ^14,15,17–22^.

**Figure 1:**
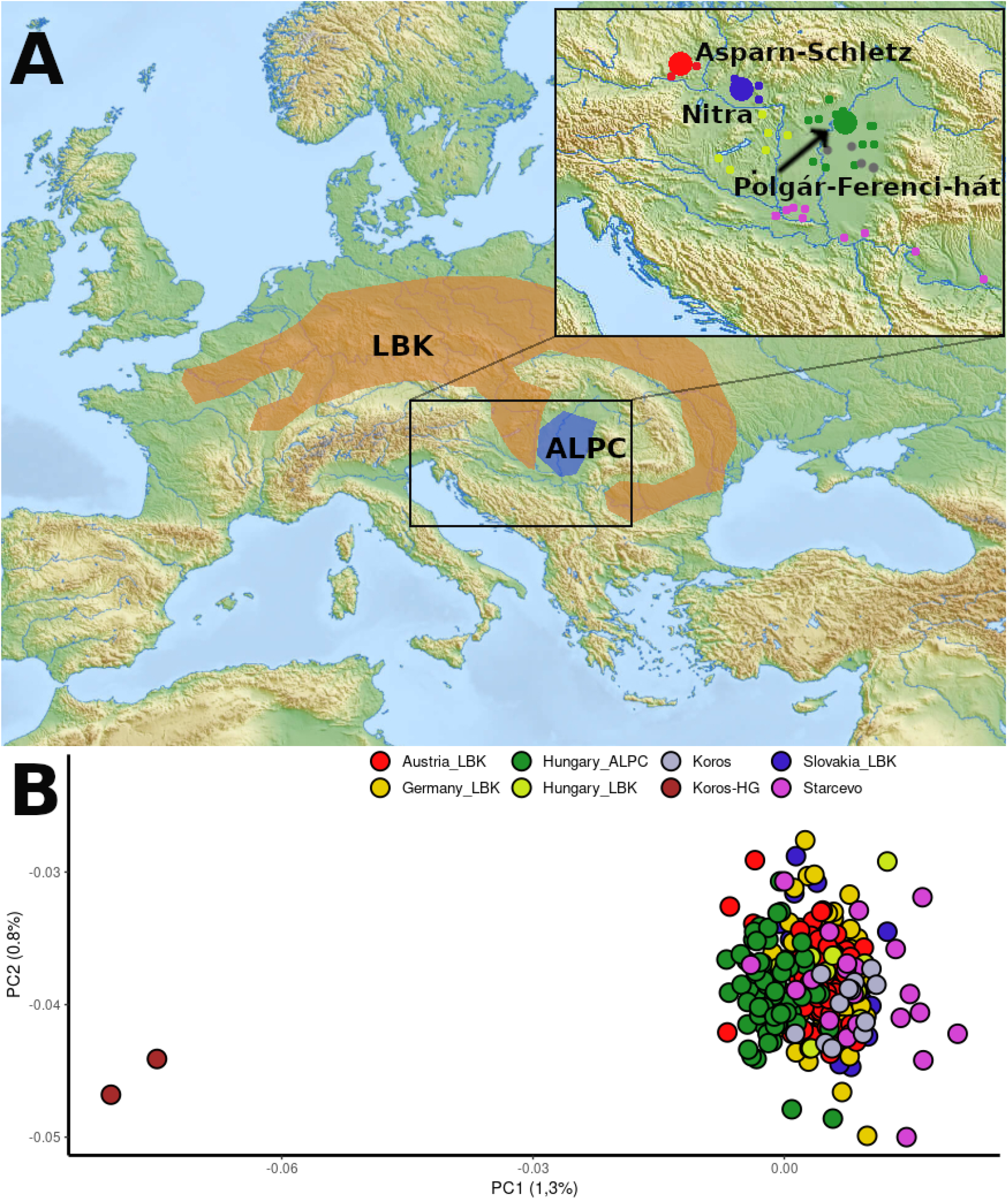
A) Map of studied sites in central Europe. Circle size corresponds to the number of individuals (large circles sites with more than ten). B). PCA shows the clustering of the LBK and the position of individuals along the X axis, indicating differential WHG affinities and showing that WHG (represented by two Körös culture outliers with entirely WHG ancestry) are more closely related to ALPC

### Elevated WHG ancestry in the eastern LBK

We used *smartpca* ^23^ to perform a Principal Components Analysis (PCA) (Figure 1B) on genome-wide data from present-day European populations genotyped on the Affymetrix Human Origins SNP array, and then projected the ancient individuals. The PCA distinguishes the Carpathian Basin ALPC from the rest of the Neolithic populations, with the ALPC individuals systematically closer to the WHG. The Körös and Starčevo individuals cluster with the western LBK, suggesting that the analyzed ALPC individuals, which might be the result of mixture between an early Neolithic population and additional WHG.

We grouped individuals based on archaeological culture and geography (proxied by present-day country): Austria_LBK, Slovakia_LBK, Hungary_LBK, Hungary_ALPC, and Germany_LBK. We estimated ancestry proportions with *qpAdm*, using as proxies for the sources a pool of Balkan early farmers with little or no WHG admixture (Balkan_N) and a pool of western European hunter-gatherers (WHGA) ^25^. As Right reference outgroups, we used pools of Turkey_N, ancient Africans, and Mesolithic European hunter-gatherers more divergent in time or space (WHGB) (Supplementary Methods). We used *qpWave* to identify significant outliers from the main cultural and geographical groups at *p-value*<0.05 adding the tags HGEXC (“hunter-gatherer excess”) and EEFEXC (“Early European Farmer excess”) (Supplementary Table 1, Supplementary Figure 1). Eastern LBK Hungary_ALPC individuals have, on average, 11.5±0.4% WHG ancestry (*p*=0.73 for fit) (Figure 2). In contrast, Slovakia_LBK and Austria_LBK individuals have an average of 4.0±0.4% WHG (*p*=0.06). Ten western LBK individuals from Transdanubia (Hungary_LBK), which do not fit the *qpAdm* model (*p*=0.001) (Supplementary Table 3 and Supplementary Section 5).

**Figure 2:**
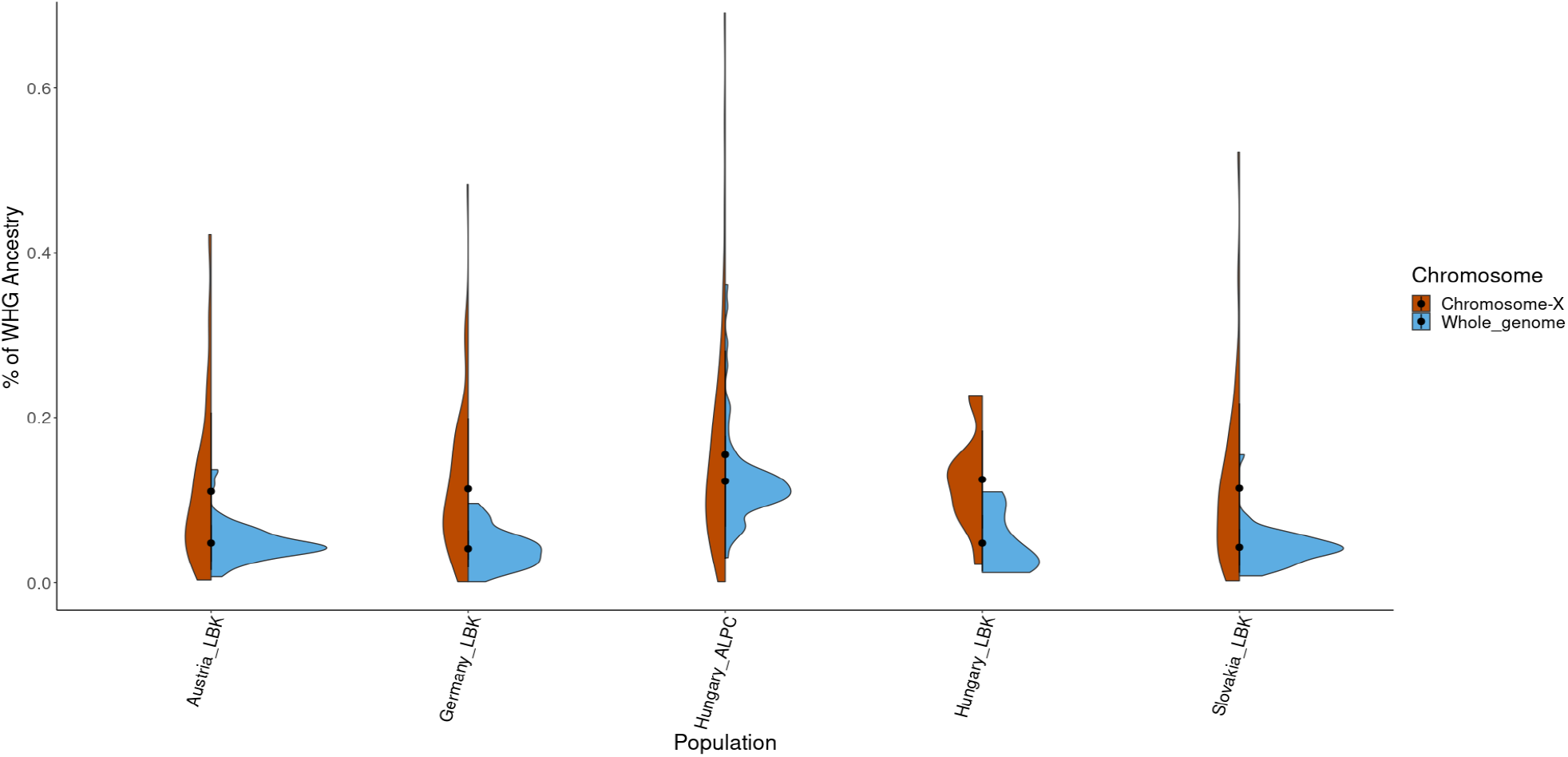
Histograms of point estimates of ancestry proportions of LBK and ALPC individuals for both the autosomes and X-chromosome (generated with ggplot2 ^24^).

### WHG-farmer mixture without significant evidence for sex bias

We used DATES ^26^ to estimate the age of admixture in WHG and Early European Farmers. Consistent with previous findings but now with higher resolution^21^ we infer that the mixture occurred on average ∼400 years before the Austrian_LBK, Slovakia_LBK, and Germany_LBK populations lived (range of 95% CI: 6057-5536 BCE), and 400 years before the ALPC individuals (range of 95% CI: 5968-5842 BCE) . This suggests a scenario in which the dawn of the archaeologically defined LBK culture was marked by the completion of a period of mixture, reflecting a social incorporation of WHG communities which plausibly could have been part of the process by which the LBK distinguished itself from preceding cultures.

Some degree of mixture with WHG continued into the LBK period itself, as documented by individuals at the early LBK site of Brunn (Austria) who show evidence of admixture in the last couple of generations before they lived ^17^ (Supplementary Table 6). We found further evidence for this using the RFMix ^27^ method, where we inferred the locations and size of segments of WHG ancestry in each LBK individual after filling in missing genotypes and phasing the data using the imputation engine GLIMPSE ^28^ (Supplementary Figure 2, Supplementary Table 5, Supplementary Section 8). We correlated the summed length of inferred WHG segments from RFMix greater than 0.2 cM to *qpAdm* estimates of WHG ancestry, and observed a high Pearson correlation coefficient of 0.96, suggesting that these inferred segments are largely reflecting true WHG admixture (Supplementary Figure 3). We identified long putative WHG segments in some ALPC individuals (up to 55 cM, individual I21902 from Polgár-Ferenci-hát, 5371-5216 cal BCE) which can only arise in the context of mixture in the last few generations in their history, similar to the pattern at Brunn. Furthermore, at the ALPC sites of Polgár-Ferenci-hát with its elevated rate of WHG ancestry, we also detect significant within-community variation in WHG ancestry. In one family (“Family B”), all three individuals had significantly elevated WHG—36% for the father and 26% and 29% for the offspring—with the daughter who we estimate to be 27-28 years old at the time of her death buried with many grave goods which were otherwise uncommon at this settlement. In another family (“Family C”), the daughter had significantly elevated WHG ancestry (20%), while the father’s ancestry is typical of the site (13%). In Family B, the mother is unsampled but we can conclude her WHG ancestry was ∼9% lower than the father’s (to explain why the offspring had intermediate proportions). In contrast and by a similar calculation, in Family C we can conclude that the unsampled mother had ∼7% higher WHG ancestry than the father. Thus, we conclude that WHG admixture was mediated by people of both sexes.

We tested directly for sex-bias in WHG admixture patterns by comparing qpAdm estimates of ancestry on the X-chromosome which depends 2/3 on female ancestry, and the autosomes which equally reflect women and men. The estimates are statistically indistinguishable in all tests (Supplementary Table 3) (Figure 2), providing no evidence for a scenario either of primarily male WHG contribution to early farmers ^29^, or hunter-gatherer Mesolithic women preferencing farmers due to perceived higher status ^30^. A caveat is that null results may reflect limited precision in X chromosome *qpAdm* estimates.

### Genetic evidence of LBK patrilocality at both the continental and local scales

The large sample size of LBK individuals analyzed in this study allows us to perform a continental scale comparison of patterns of variation on the Y chromosome reflecting the history of the entire male line, and mitochondrial DNA which reflects the history of the entire female line. In the Y-chromosome analysis, we detect previously unappreciated geographic variation across the LBK (Fig 4) (a χ^2^(209,42)=242 test for heterogeneity is highly significant at *p*<10^-12^), with haplogroup G dominant in the Slovakian, German, and Hungarian LBK; haplogroup C in the Austrian_LBK; and most Hungary_ALPC individuals carrying haplogroup I (61%), associated with Mesolithic populations such as those of the Iron Gates regions of Serbia and Romania ^19,31^. In contrast, we do not detect significant structure in mitochondrial DNA haplogroup frequencies, with no haplotype with a frequency greater than 30%, and no evidence for haplotype frequency differences across the regional groups (Supplementary Figure 4), (χ^2^ (449,54)=61.3, *p*=0.23). Taken together, these results provide evidence of limited gene flow among LBK communities on the male line, which could be explained by a much higher rate of movement of females between communities.

We also find genetic evidence for patrilocality in LBK communities by studying the distributions of close relatives in the two cemeteries where we detect many relationships (Figure 3). At Nitra we detect nine families including a pedigree spanning four generations, and at Polgár-Ferenci-hát we detect 4 families including one with 13 individuals. Combining the two cemeteries, we find that relatives up to the 3^rd^ degree (Supplementary Table 7, Supplementary Sections 2.2 and 2.3) tend to be buried together more often than random pairs of individuals At Nitra, we did not detect significant differences in the number of relatives between males and females χ^2^ (47,1)=0.14, *p*=0.70. In contrast, we detect strong evidence of patrilocality at Polgár-Ferenci-hát, with more relatives for males (21 of 22) than for females (14 of 23): χ^2^(45,1)=7.78, *p=*0.005 (Table 1).

**Figure 3:**
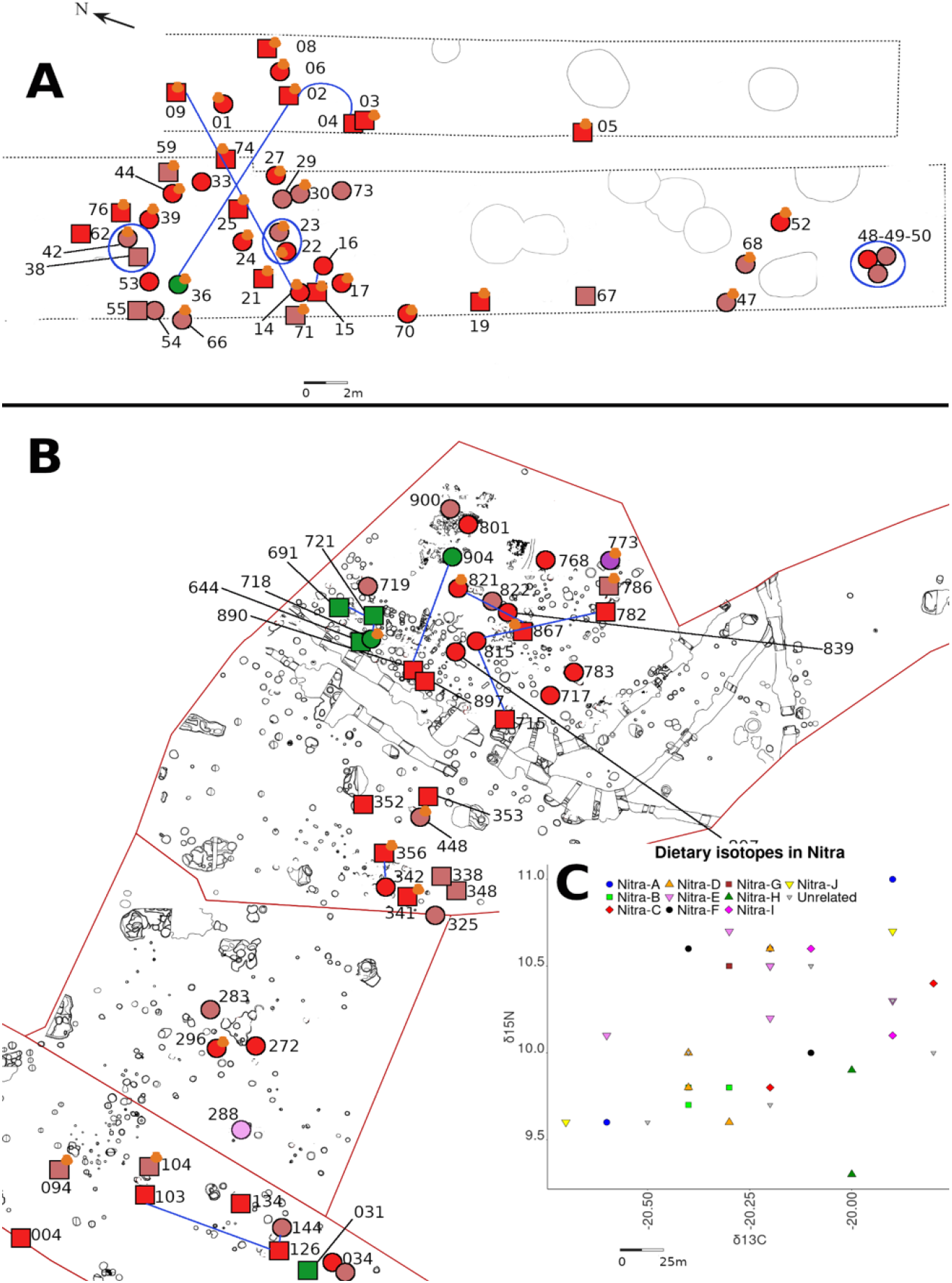
Burial layouts for A) Nitra (top) and B) Polgár-Ferenci-hát (bottom). Each symbol represents one individual: squares males, circles females. Red denotes main genetic cluster, green WHG outliers, violet EEF outliers. Light brown are children. Blue lines or circles are 1^st^-degree relatives, and the yellow pottery symbol grave goods in burials. C) Dietary isotopes at Nitra coded by families. Families at Nitra do not cluster in dietary-specific groups. All plots restrict to individuals with qpAdm estimates.

**Table 1:** Patterns of relationships at three LBK sites with substantial new data (*p<0.05)

|  | Polgár-Ferenci-hát<br>(n=45) | Nitra<br>(n=47) | Schletz<br>(n=93) |
| --- | --- | --- | --- |
| Ratio<br>related/unrelated | 0.83 | 0.65 | 0.15 |
| Ratio of<br>males/females related | 0.94/0.69* | 0.66/0.65 | 0.1/0.05 |
| Average no. of<br>relatives | 1.1 | 0.60 | 0.1 |

### No evidence for kinship-associated differences in mobility and diet

We analyzed the findings of genetic relatedness together with dietary (carbon, δ^13^C, and nitrogen, δ^15^N) and strontium isotope data (^87^Sr/^86^Sr) ^32^ (Supplementary Section 3). We did not perform similar analyses for Schletz as we had dietary isotopic data for too few individuals and too few detected genetic relatives.

We detect significant within-family variation in mobility isotopic measurements both at Nitra (Levene’s test for variances n=12, *p*=0.01) and at Polgár-Ferenci-hát (Levene statistic for the difference in variance = 16.74, *p=*0.001). This suggests that the people at both sites genetically related individuals varied in the places where they resided over their lifetimes.

We next tested for significant differences across families in their dietary patterns but found no strong signals. The only notable correlation we detect is at Nitra, where we found a marginally significant signal of variation across families for δ^13^C carbon isotopes (Kruskal-Wallis=17.20, N=26, *p*=0.04), providing some evidence that families sourced food from different landscape contexts, either through variation in direct consumption or through variation in consumption of animals eating these plants^27^. However, in light of the fact that we carried out multiple hypothesis tests (below), the observation of one marginally significant signal of correlation like this should not be interpreted as strong evidence.

We do not detect significant variation in strontium isotope ratios across families at Nitra (Mann-Whitney U test, n=21, *p*=0.16), nor do we detect a correlation between family structure and the presence of grave goods (Supplementary Section 2.1, Supplementary Table 8) (δ13C, Kruskal-Wallis=4.99, *p*=0.17; δ15N, Kruskal-Wallis = 1.45, *p*=0.69). At Polgár-Ferenci-hát, we also do not detect variation in isotopic ratios across families: δ13C, Kruskal-Wallis = 4.99, *p*=0.17; δ15N, Kruskal-Wallis = 1.45, *p*=0.69 (Supplementary Figure 5).

### Variation across the LBK in community size, migration, and mate choice patterns

We attempted ancient DNA analysis from all excavated skeletons from Schletz, corresponding to 103 individuals from the ditch system associated with a massacre, and 19 individuals from settlement burials. A total of 85 of the 122 individuals had enough genomic data for high-resolution analyses (Supplementary Table 1). Of the 71 individuals with genome-wide data from the base of the ditch system, including 51 genetic males and 20 genetic females, we detected only a single pair of 1st/2nd-degree relatives, and possibly two pairs of individuals between ditch and settlement contexts. Only six of the 71 analyzed individuals from Schletz are related up to the 3^rd^ degree, contrasting with much higher rates at Nitra and Polgár-Ferenci-hát (Table 1). We identified a single relationship between an older male adult (I24892) and a subadult (I24280) from within the massacre context, providing further evidence that this was not an event that affected only a small community that might have been expected to include more families.

We used HapNe ^33^ to infer the effective population size trajectory of unrelated individuals from the Schletz massacre in the hundreds of years before they lived (n=56). We find no evidence for a contraction in the gene pool in the hundreds of years before the individuals lived, which could be explained if the people massacred at Schletz were drawn from many communities and not a single community that would be expected to be drawing from a more local gene pool. In contrast, at Nitra (n=18), we observe strong evidence for such recent geographic isolation and community substructure (Figure 4, Supplementary Section 6).

**Figure 4:**
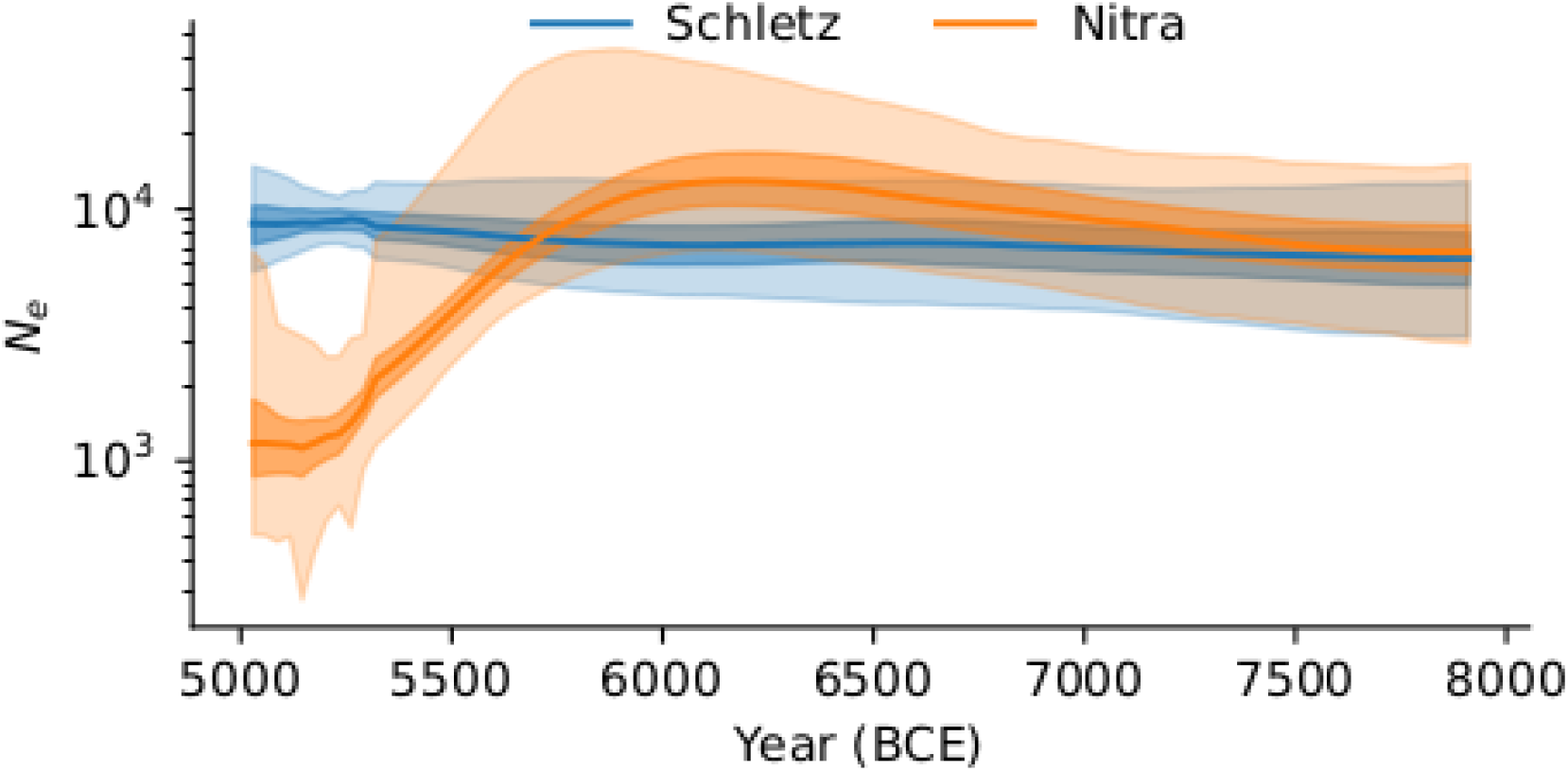
Inferred population size trajectory of Schletz and Nitra. The recent contraction in Nitra likely reflects undetected families in the sample, while the Schletz individuals have no evidence of being more closely related to each other than more widely sampled LBK.

**Figure 5:**
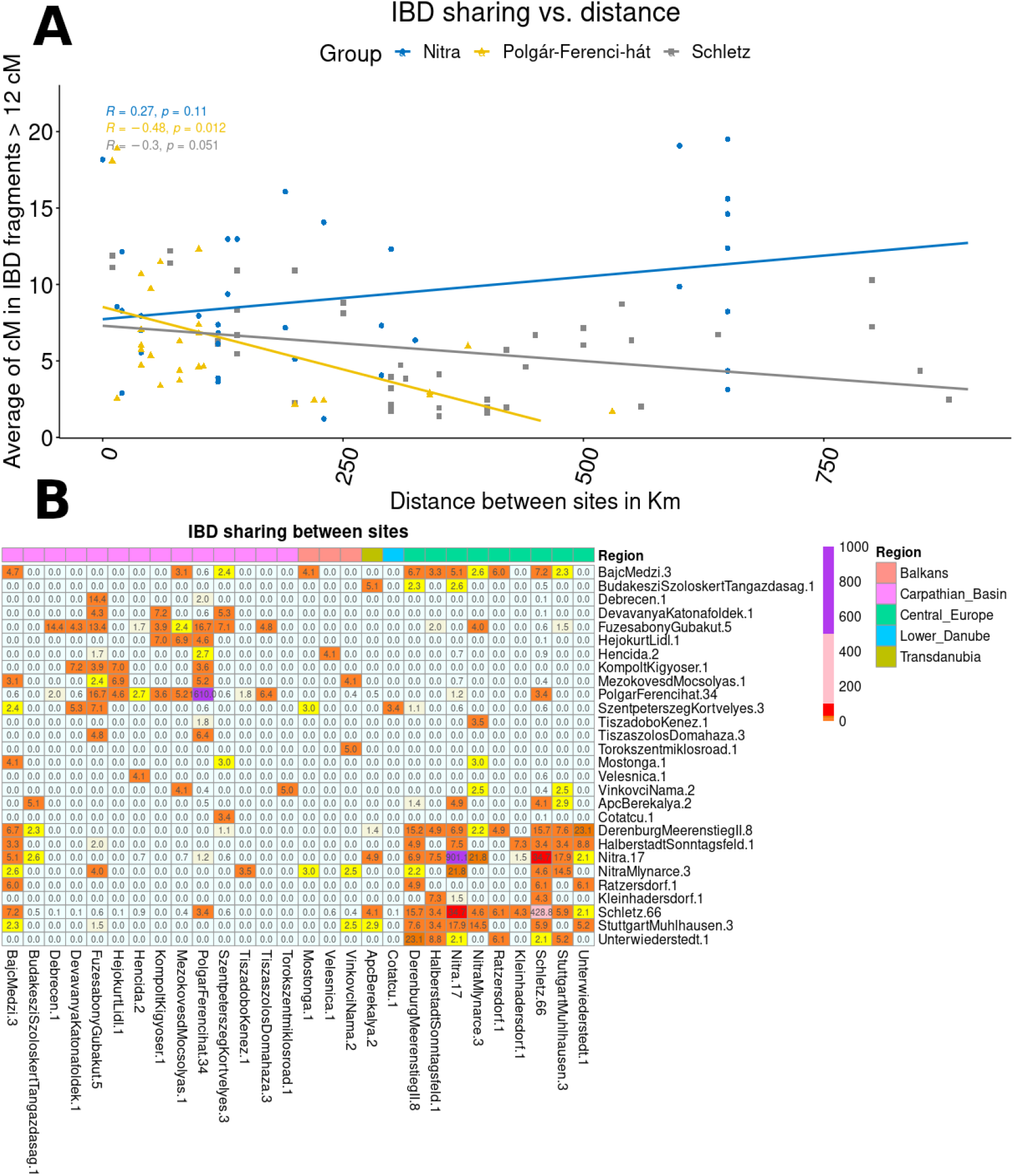
IBD patterns: A) Regression of summed IBD >12cM shared between individuals of each pair of sites (averaged over all pairs), and geographic distance. Polgár-Ferenci-hat has more connections with closer sites supporting a localized ALPC community, while Schletz and Nitra do not show a clear association with distance, as would be expected if the western LBK expansion was so rapid that nearby groups were hardly more closely related than groups far apart. B) A heatmap showing the intensity of IBD, presenting the average total length of IBD segments > 12cM shared between all possible pairs, by area or period. The numbers after the site names show the number of individuals per site included in these analyses.

Further evidence for the Schletz individuals being drawn from a much larger population than those at the other sites comes from IBD sharing patterns between the studied individuals (Supplementary Table 9), inferred based on analysis of the imputed and phased dataset. We observe significantly less average sharing of IBD segments >12 cM among individuals at Schletz (26 cM) than at Nitra (174 cM) or Polgár-Ferenci-hát 158 cM. The reduction is significant (*p*=0.001), even after excluding 1st, 2nd, and 3rd-degree relative pairs (*p*=0.005), showing the signal is driven by distant relatives in sites, not just close relatives.

Eighth individuals from Polgár-Férenci-hát and four from Nitra have elevated rates of Runs of Homozygosity (ROH), a type of data that provides evidence about whether or not individuals reproduced within their own genetic lineages?^34^. In contrast, the rest of the individuals at these sites and all of those from Schletz have no segments with ROH >4 cM ^34^ (Supplementary Figure 6).

The genetic data give evidence of two separate regional networks: one for the Carpathian Basin where the across-site rate of sharing averages 918 cM, and one for Central-Western Europe where the across-site rate of sharing averages 490 cM, but with a lower 31 cM of sharing across regions. This is in accord with archaeological studies that imply that Nitra and Polgár-Ferenci-hát are associated with different LBK expansions and periods ^21,35,36^. We further observe that the rate of IBD sharing decreases significantly with distance from Polgár-Ferenci-hat (*p=*0.011) consistent with the idea of a localized network of people within the ALPC. In contrast, there is weak or no detectable association of IBD sharing with geographic distance in the western LBK, as would be expected if the western LBK expansion was so rapid that nearby groups were hardly more closely related than groups far apart. Finally, since the Hungary_ALPC individuals have on average 16.5 cM in ROH and are the LBK group with the largest fractions of their genome in ROH, they appear to exhibit more restricted reproduction practices than the more widespread western LBK.

### Screens for natural selection based on high-frequency long-range haplotype

We scanned the imputed diploid genotype data for the LBK and ALPC individuals for signals of selection by searching for haplotypes that had evidence of being very recent in origin based on their common frequency and large scale. Because of the poor haplotype phasing expected for ancient genomes, we carried out these scans with both the phased andunphased versions of the iHS and nSL scores, as implemented in Selscan 2.0 ^37^. We also used BetaScan^38^ to detect signals of long-term balancing selection in the population.

We detect evidence of long-term balancing selection in the HLA region on chromosome 6, with elevated B1 scores (Figure 6), consistent with previous evidence of balancing selection at this locus in Neolithic Europeans ^39^. A second notable finding is 29 genes with evidence of balancing selection in the ALPc and LBK (Supplementary Table 12). Most were also reported as significant outliers based on analysis of patterns of variation in modern Europeans ^40^.

**Figure 6:**
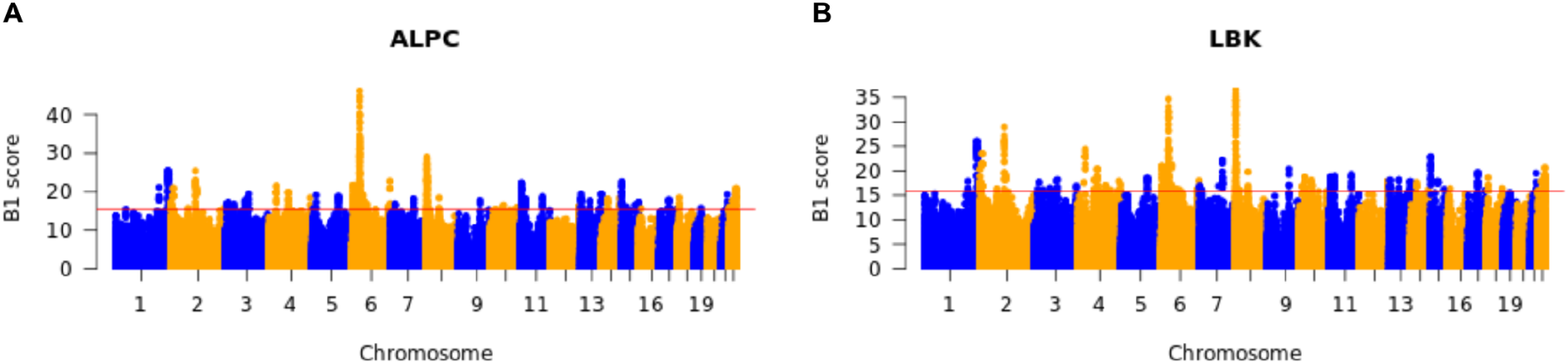
(A) B1 scores in the ALPC. (B) B1 scores in the LBK. B1 shows regions with balancing selection, the highest signal on chromosome 6 at HLA.

We identified 40 genes with evidence of positive selection in the LBK (Supplementary Tables 11-12, Supplementary Section 7, Supplementary Figure 7), including notable examples associated with pigmentation. Variation in the *BNC2* influences human pigmentation and has also been found to be affected by natural selection in modern Europeans^41–43^. The *PRKCH* gene encodes the PKCη protein in melanocytes which is involved in the protein kinase C-dependent pathway regulating melanogenesis ^44^. The *PTPRN2* gene had a higher level of expression in lightly pigmented melanocytes than in darkly pigmented melanocytes, similar to *SLC45A2* which contains one of the strongest known signals of pigmentation selection in Europe ^45^. When we correlate the WHG local ancestry components with our selection values, the non-WHG ancestry seems to contribute more to sites under selection (Supplementary Section 7, Supplementary Figure 8).

## Discussion

Our study reveals differences in kinship structure, admixture, demography, and ancestry across the LBK. We report an average of around 12% WHG ancestry at Polgár-Ferenci-hát, a proportion that reached as high as 35% in some individuals. This contrasts with the much lower average among the studied individuals from Schletz (an average of 4% with a range of up to 14%) and Nitra (an average of 4% with a range of up to 8%). This suggests that the admixture between farmers and hunters in the Carpathian Basin was more extensive than among the more westerly LBK communities. This admixture shows no evidence of a sex-biased trend despite the high fraction of Y-chromosome haplogroups associated with WHG.

Correlation between isotopic and genetic shows no statistical differences in diet and mobility patterns between families in Nitra and Polgár-Ferenci-hát, but we find evidence for high variation in mobility within families at least at Nitra. We observe no evidence of correlation of genetic patterns to archaeological markers of social status.

We find that at both Nitra and Polgár-Ferenci-hát, relatives are buried closer to each other than non-relatives. Polgár-Ferenci-hát males had significantly more relatives than females. This pattern, combined with the evidence of limited regional diversity in the Y-chromosome and long IBD tracks, is consistent with a patrilocal society. We observed much less IBD across western LBK sites and ALPC sites than within either community, suggesting they were part of different mating networks.

The proportion of relatives in Schletz is lower than at any other LBK site analyzed. We only identified relationships between males and children and only one with an adult male. This raises doubts regarding the idea that the individuals recovered at the ditch represent a local community, and instead suggests that people massacred at this key were likely drawn from a widespread population ^46^. When comparing Schletz and Nitra, we find evidence that Schletz but not Nitra represents a large population. One possibility is that Schletz was a central site that drew a population from a larger area in times of stress, such as outbreaks of violence observed at other LBK sites ^9^. Another explanation could be that communities in the broader LBK expansion area were formed with few biologically related individuals, as at Derenburg-Meerenstieg II and Stuttgart-Mühlhausen, Germany. In any case, our results suggest that frequent mobility between sites was a factor in many LBK communities ^47^. A lack of related individuals has also been found in the Eneolithic massacre of Potocani, Croatia ^48^.

Our results illuminate how whole-cemetery ancient DNA, with isotopic and archaeological data, can reveal the structure of past societies as well as evidence for local variations in mobility and diet, shedding light on unappreciated aspects of past human behaviour.

## Acknowledgments

We dedicate this paper to the memory of Tibor Paluch, who passed away during its writing. This study followed principles for ethical DNA research on human remains ^49^. We thank the authorities and sample stewards, including museums, museum curators, and archaeologists; Nicole Adamski, Rebecca Bernardos, and Kristin Stewardson for help with sample handling and sample preparation; and Zhao Zhang for bioinformatics support. Ancient DNA data generation and analysis was supported by the National Institutes of Health (NIGMS GM100233), the John Templeton Foundation (grant 61220), a gift from Jean-Francois Clin, the Howard Hughes Medical Institute, and by the Allen Discovery Center program, a Paul G. Allen Frontiers Group advised program of the Paul G. Allen Family Foundation. The physical anthropology and archaeological work were supported by a grant from the Hungarian Research, Development, and Innovation Office [project number: FK128013] (T.H.), the Bolyai Scholarship of the Hungarian Academy of Sciences (T.H.), and the New National Excellence Program of the Ministry for Innovation and Technology from the source of the National Research, Development and Innovation Fund (TSZ, T.H.). Zdeněk Tvrdý was supported by the Ministry of Culture via institutional funding for the long-term conceptual development of the Moravian Museum research organization (DKRVO, MK000094862). Susanna Sawyer was funded by the Austrian Science Fund (FWF) M3108-G. Mario Novak was supported by the Croatian Science Foundation (HRZZ IP-2016-06-1450). Catalin Lazar was funded by a grant from the Romanian Ministry of Research, Innovation, and Digitisation (41PFE/30.12.2021) within PNCDI III. We thank Lindsey Buster for the critical discussion on the paper editing.

## Author contributions

P. G, R. P, P. B, M. T-N, A. A., and D. R conceived the study. M. T-N, A. A, F. P, J. P, C. L, T. P, MK. S, M. S, Z. B, F. N, L.D. S, O. CL, T. H, Z. H, P. M, E.G. N, Z.M. V, AM. H, LA. H, K.T. B, L D, T. S, J. J, M. S, F. Z, S. S, K. T, P. C, I. P, R. P, B. H, C. S, G. L, Z. T, D. B, M. R, M. N, J. D, T. H, P. R, A. P, B. V. provided samples. O. C, D. M.F., C. S, K. B, F. C, G. B, D. K, S. F, K. TO, K.M, V. O, E. H performed the experiments. P. G, X. H, P. B, I. O, P. F, M. K, S. S, H. R, A. A, P. F. P, O. C, M. H, S. M, M. M, A. M, I. L, R. F, N. B. N. W, F. Z, L. Q, A. K, AM. L, analyzed the data. P. G, R. P, P. B, and D. R wrote the manuscript with inputs from all co-authors.

## Methods

### Ancient DNA data generation

The 319 individuals screened in this study were sampled with authorization from the authorities responsible for each of them. Bones or teeth were powdered, and DNA was extracted in dedicated laboratories. DNA was extracted from powder using an automated protocol with silica-coated magnetic beads and Dabney binding buffer ^50^. DNA extracts were converted to double-stranded libraries using a partial UDG treatment ^51^. Amplified libraries were enriched using two rounds of consecutive hybridization capture enrichment 1240k strategy ^52,53^). Captured libraries were sequenced on an Illumina NextSeq500 instrument with 2 × 76 cycles (2 × 7 cycles for the indices) or an Illumina HiSeq X10 with 2 × 101 cycles (2 × 7 for the index). We trimmed adapters, merged paired-end sequences, and aligned to the human genome (hg19) and mitochondrial genome (RSRS) using BWA 0.6.1 ^54^. The computational pipelines are available on GitHub (https://github.com/DReichLab/ADNA-Tools, https://github.com/DReichLab/adna-workflow).

We evaluated ancient DNA authenticity using several criteria: a rate of cytosine deamination at the terminal nucleotide above 3%; a ratio of Y to combined X + Y chromosome sequences below 0.03 or above 0.35 ^55^(intermediate values are indicative of the presence of DNA from at least two individuals of different sex); for male individuals with sufficient coverage, an X chromosome contamination estimate whose lower bound of the 95% confidence interval is <1.1% (all but one below 0.5%); and an upper-bound rate for the 95% confidence interval for the rate to the consensus mitochondrial sequence that exceeds 95%, as computed using contamMix-1.0.10 ^56^. We added tags to samples that gave evidence of contamination by any of these criteria and discarded samples with at least two signals of contamination.

### Genetic sex, mitochondrial and Y chromosome haplogroup determination

To determine genetic sex, we searched for evidence of a Y chromosome by computing the ratio of Y-chromosomal 1240k positions with available data divided by the number of X-chromosomal and Y-chromosomal 1240k positions with available data. Individuals with a ratio of more than 0.35 were considered genetic males, and individuals with less than 0.03 were considered genetic females. To check for sex chromosome aneuploidies, we computed the mean coverage on X-chromosomal and Y-chromosomal 1240k positions. We normalized these values by autosomal coverage on 1240k positions for each individual. We did not find any evidence of sex chromosome aneuploidies.

To determine mitochondrial haplogroups (Supplementary Table 1), we constructed a consensus sequence with samtools and bcftools ^57^, restricting to sequences with a mapping quality of >30 and a base quality of >30. We called haplogroups with Haplogrep2 ^58^.

We determined Y chromosome haplogroups (Supplementary Table 1) based on the nomenclature of the International Society of Genetic Genealogy (http://www.isogg.org) version 14.76 (25 April 2019), restricting to sequences with a mapping quality of 30 or more and a base quality of 30 or more.

### Biological kinship estimation and family reconstruction

We followed the same approach described by ^59^. We focused on 1st, 2nd, and 3rd-degree relatives for family reconstruction, but also noted individuals detected as relatives up to the 4^th^ degree. The complete list is reported in Supplementary Table 7.

### Principal component analysis and f-statistics analyses

We used the Western Eurasian populations genotyped on the Affymetrix Human Origins SNP array to perform Principal Components Analysis with SmartPCA ^23^. In this PCA, we projected all the samples we report in this paper (Supplementary Table 1) as well as other relevant ancient DNA data (Supplementary Table 1). The same dataset was used to perform f-statistics-based analyses using admixtools ^23^.

We performed *qpAdm* analyses following the same strategy as in Patterson et al. 2022 ^25^. Individuals labeled as Ancient_Africa, WHGB, and Turkey_N were used as right populations and WHGA and Balkan_N as left. qpWave was performed using the same strategy.

### ROH

We called ROH with the methodology described in Ringbauer et al.^34^ optimized for the study of ancient people, restricting to individuals with more than 400,000 SNPs.

### Imputation

Genomes were imputed and phased with GLIMPSE ^60^ following the methodology in ^61^.

### Local ancestry maps

We used diploid imputed genotypes to perform the analyses. We ran RFMix v2.03-r0^27^using Balkan_N and WHGA as reference populations. We plotted the results with Python 3.7.6 and Rstudio.

### Selection

The selection analysis is detailed in the supplementary material.

### Data availability

All sequencing data are freely available at the European Nucleotide Archive (ENA) with the accession number: PRJEB64177.

## Notes

### Competing Interest Statement

The authors have declared no competing interest.

