## Supplementary material for "Social and genetic diversity among the first farmers of Central Europe"

#### Table of Contents

Section 1: Supplementary Figures

Section 2: Kinship

Section 3: Isotopic data

Section 4: Site Descriptions

Section 5: Classification of individuals with genetics methods

Section 6: Population size inferences

Section 7: Selection scans in the diploid data

Section 8: Supplementary Methods

Section 9: Supplementary References

### Section 1: Supplementary Figures

**Supplementary Figure 1:** qpWave plots to test for differentiation among individuals, with each population represented in one plot. Grey color means results were highly significant (little genetically related). The number after the name of each individual relates is the point estimate of WHG ancestry from qpAdm.

1.1 Starčevo-Körös-Criş Individuals: I6699 and I1878 are labeled in our analysis as outliers with high WHG ancestry.

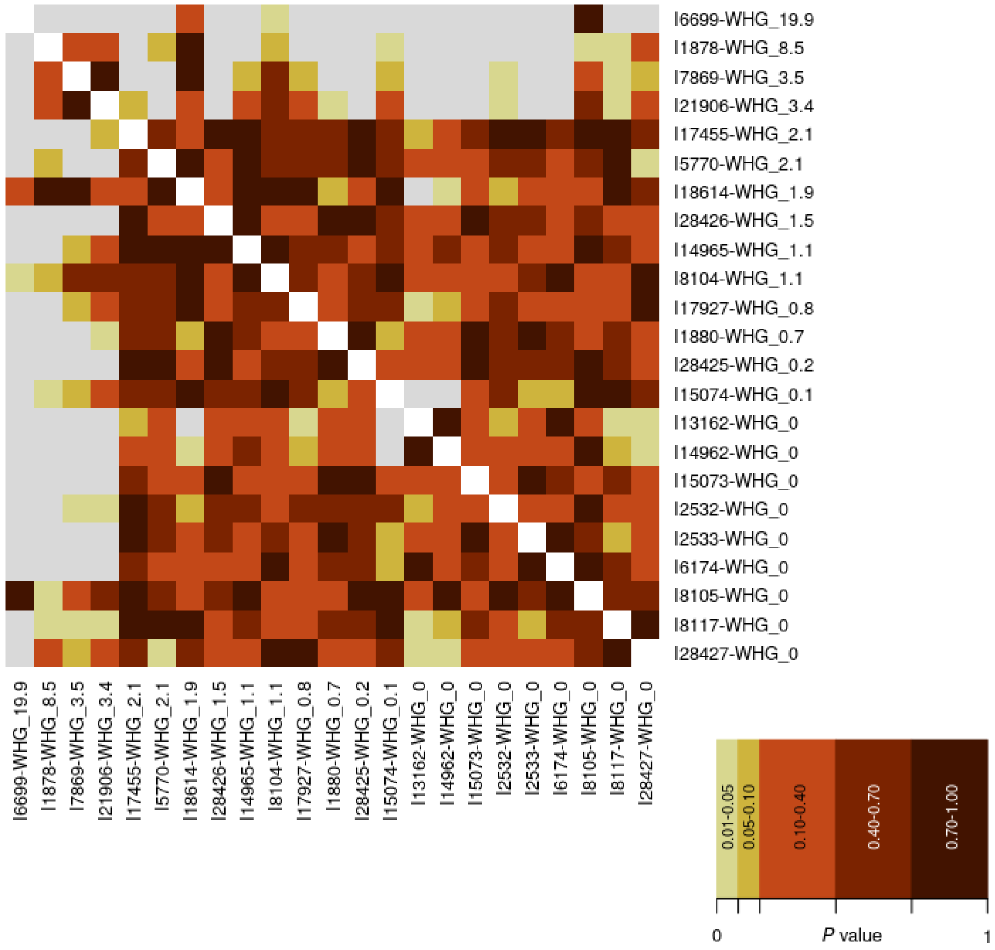

1.2 Hungary LBK Individuals: individuals I1882 and I1883 are labelled in our analysis as outliers with high WHG ancestry.

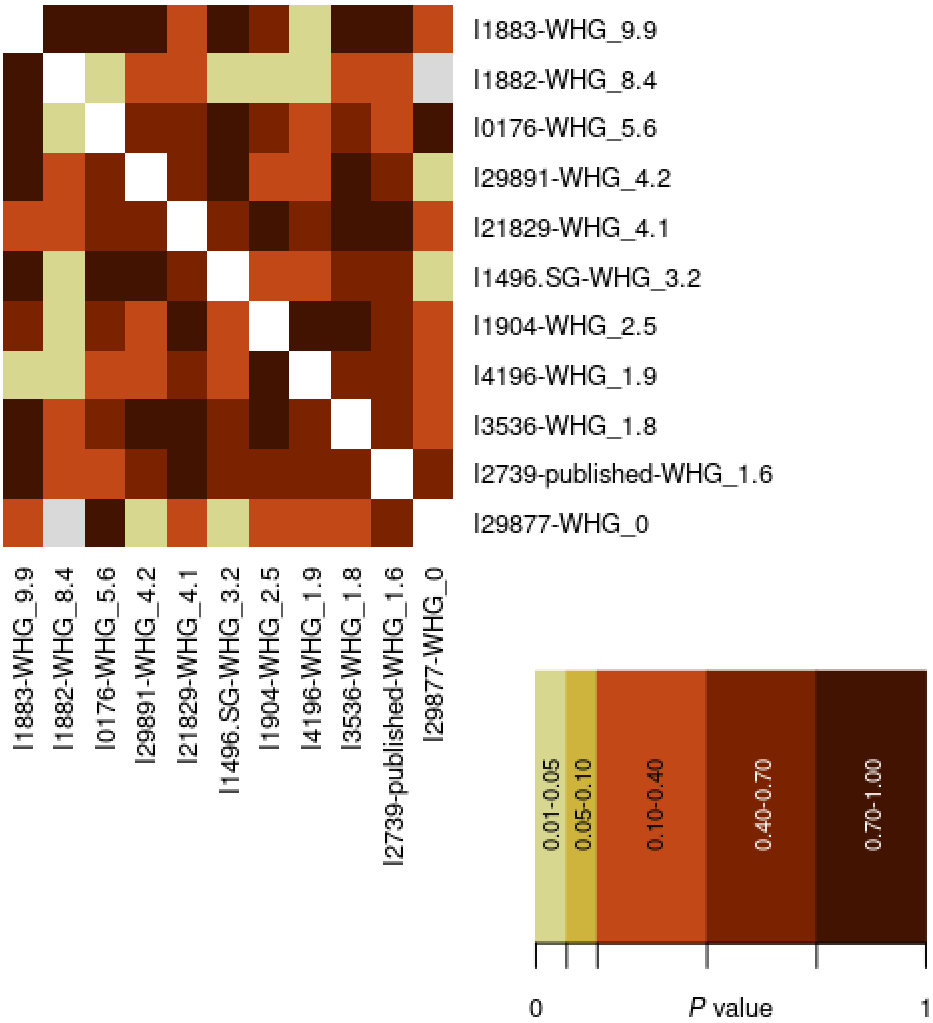

1.3 Hungary ALPC Individuals: Individuals I21898, I10349, I21902, I18660, I10350, I18656, I18695, I4186, I1499, and I21714, are labeled in our analysis as outliers with high WHG ancestry. Individuals: I21828, I21830, I10351, I10352, I10353, I18657, I21767I, 17933, I1500, I2380, I3537, and I4187 are labeled in our analysis as outliers with high EEF ancestry.

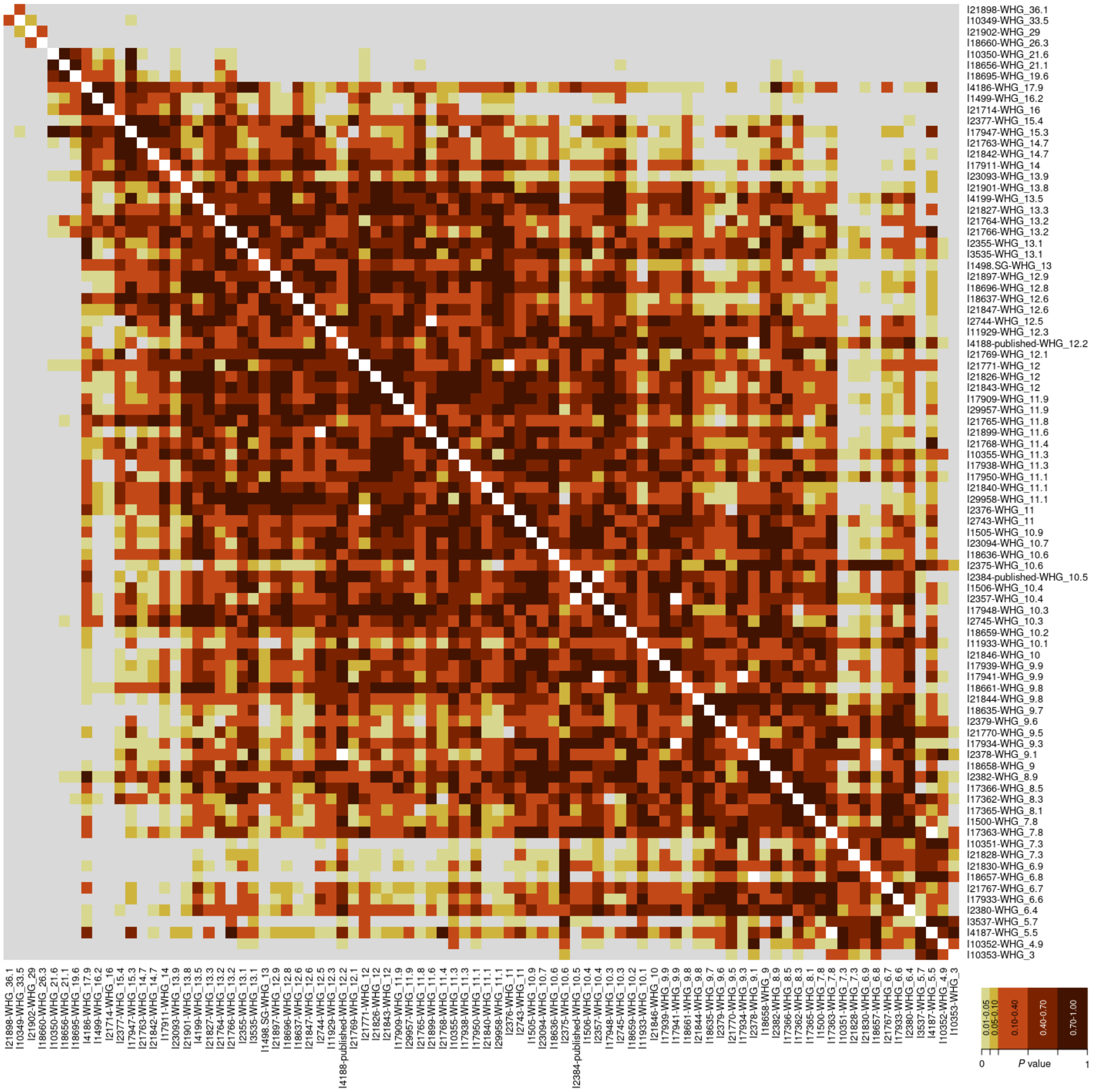

1.4 Austria LBK Individuals: Individuals I27785, I25349, I24028 are labelled in our analysis as outliers with high WHG ancestry.

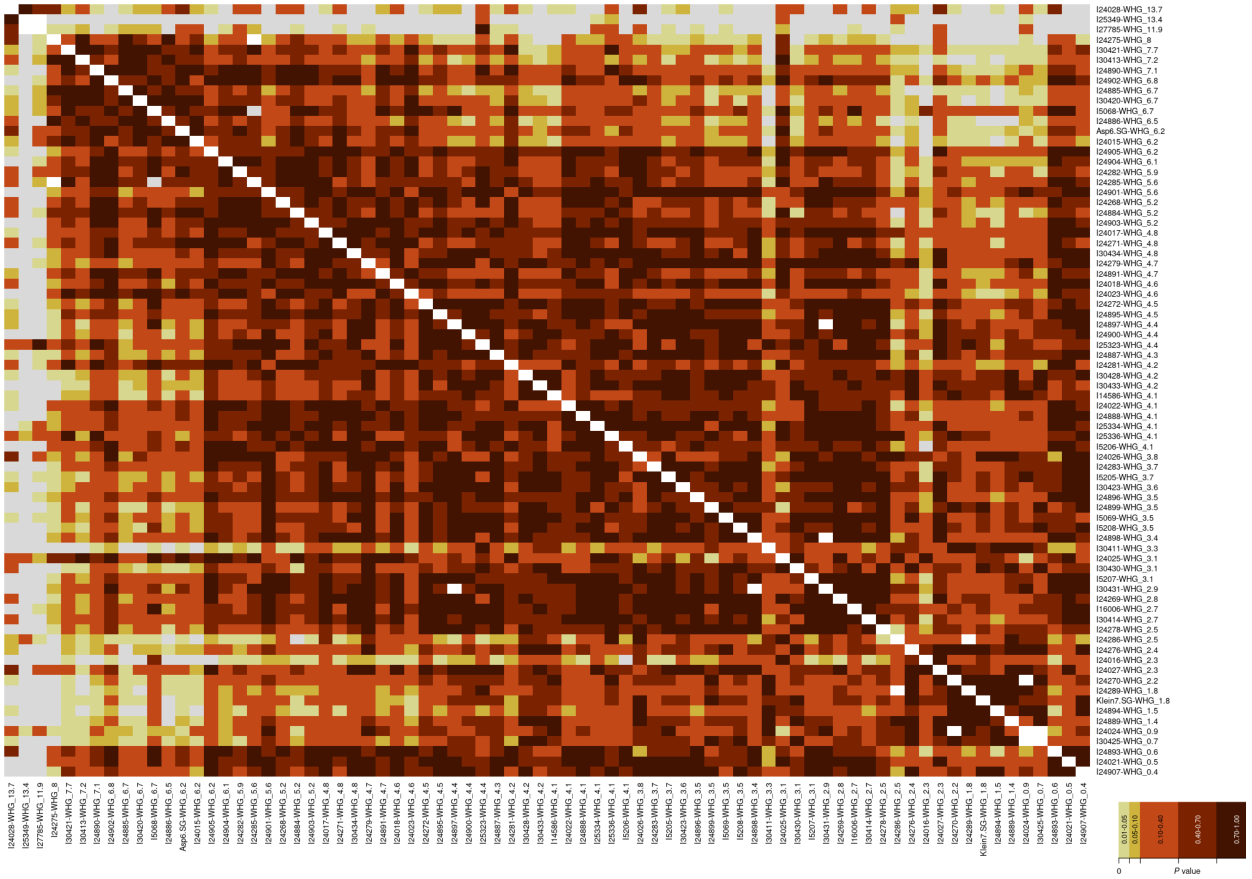

1.5 Slovakia LBK Individuals: Individual I18144 is labelled in our analysis as an outlier with high WHG ancestry.

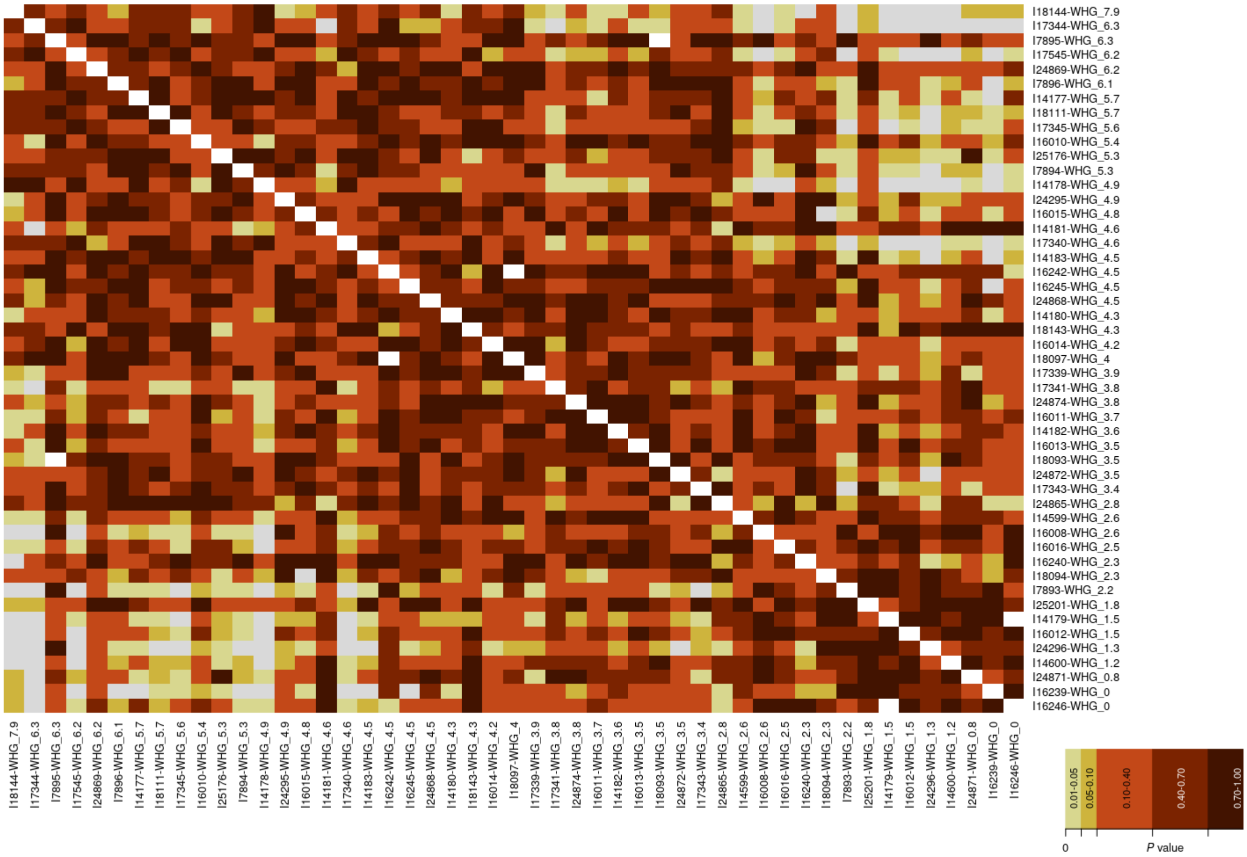

#### 1.6 Germany LBK Individuals

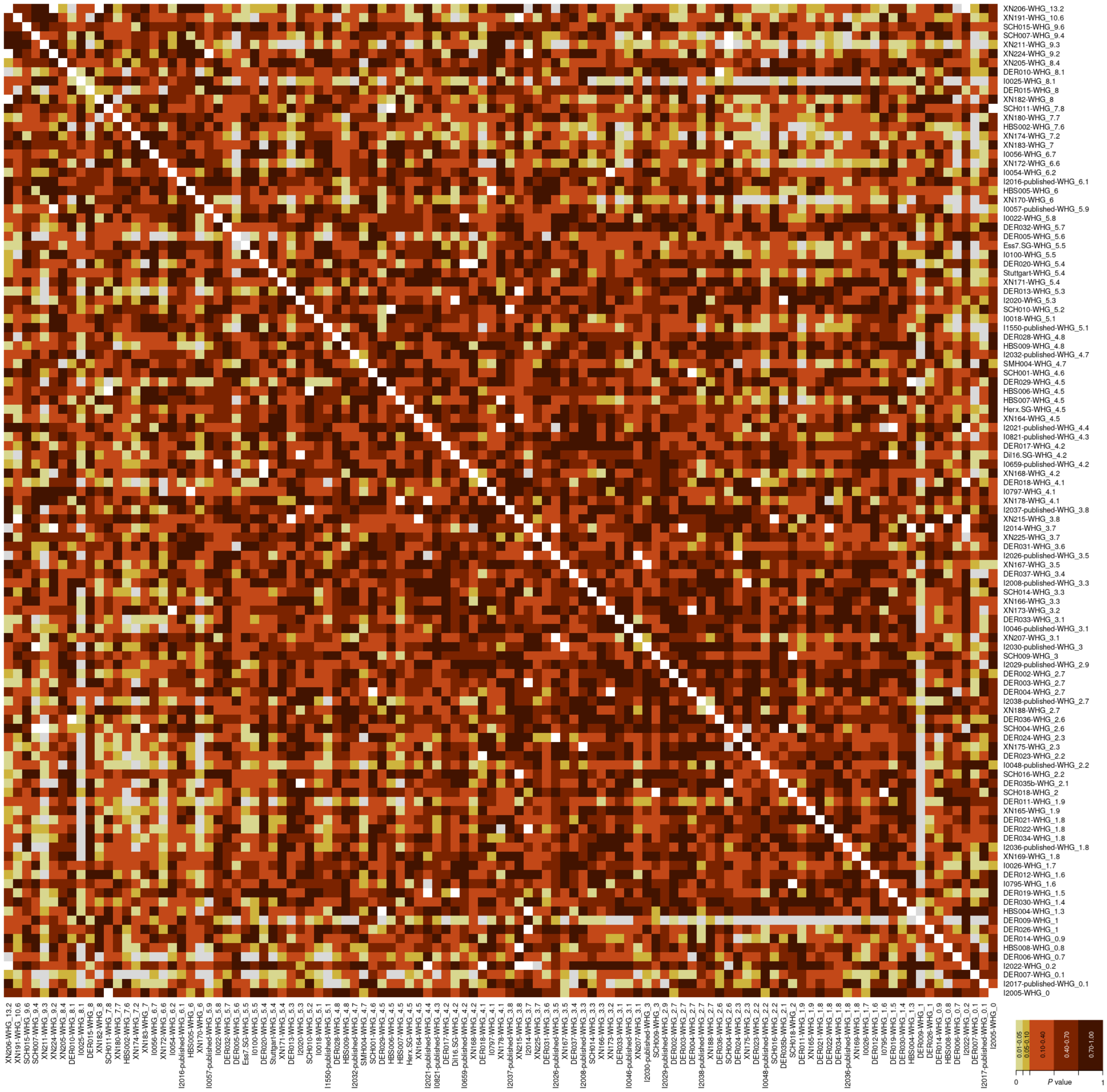

**Supplementary Figure 2:** Local ancestry maps obtained with RFMix on ALPC individuals with more than 400,000 captured SNPs imputed and phased with GLIMPSE and plotted with matplotlib 3.7.1 in Python 3.7.6. Some individuals show much longer inferred WHG segments than others, with I21902 having the longest segment, and the largest summed WHG fractions in I10349 and I21898; these two individuals also are inferred to have the highest WHG ancestry with qpAdm (Table S5).

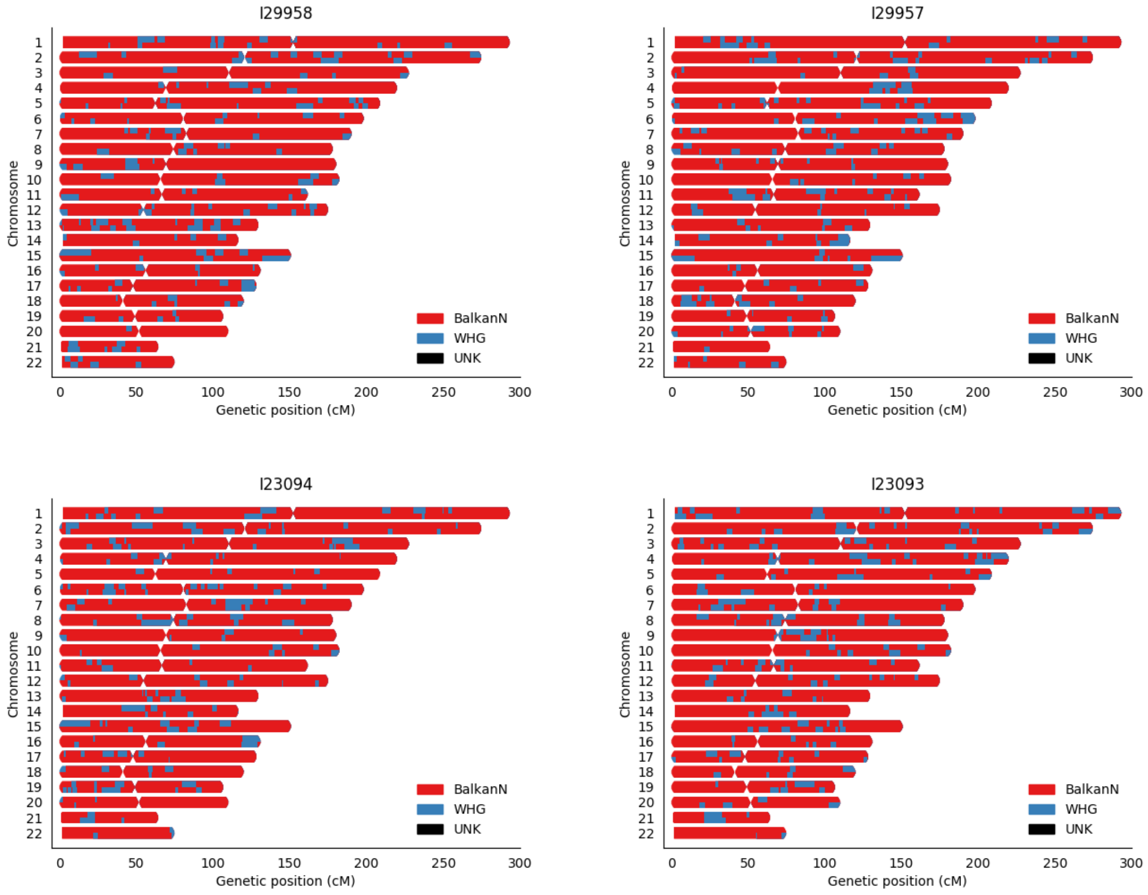

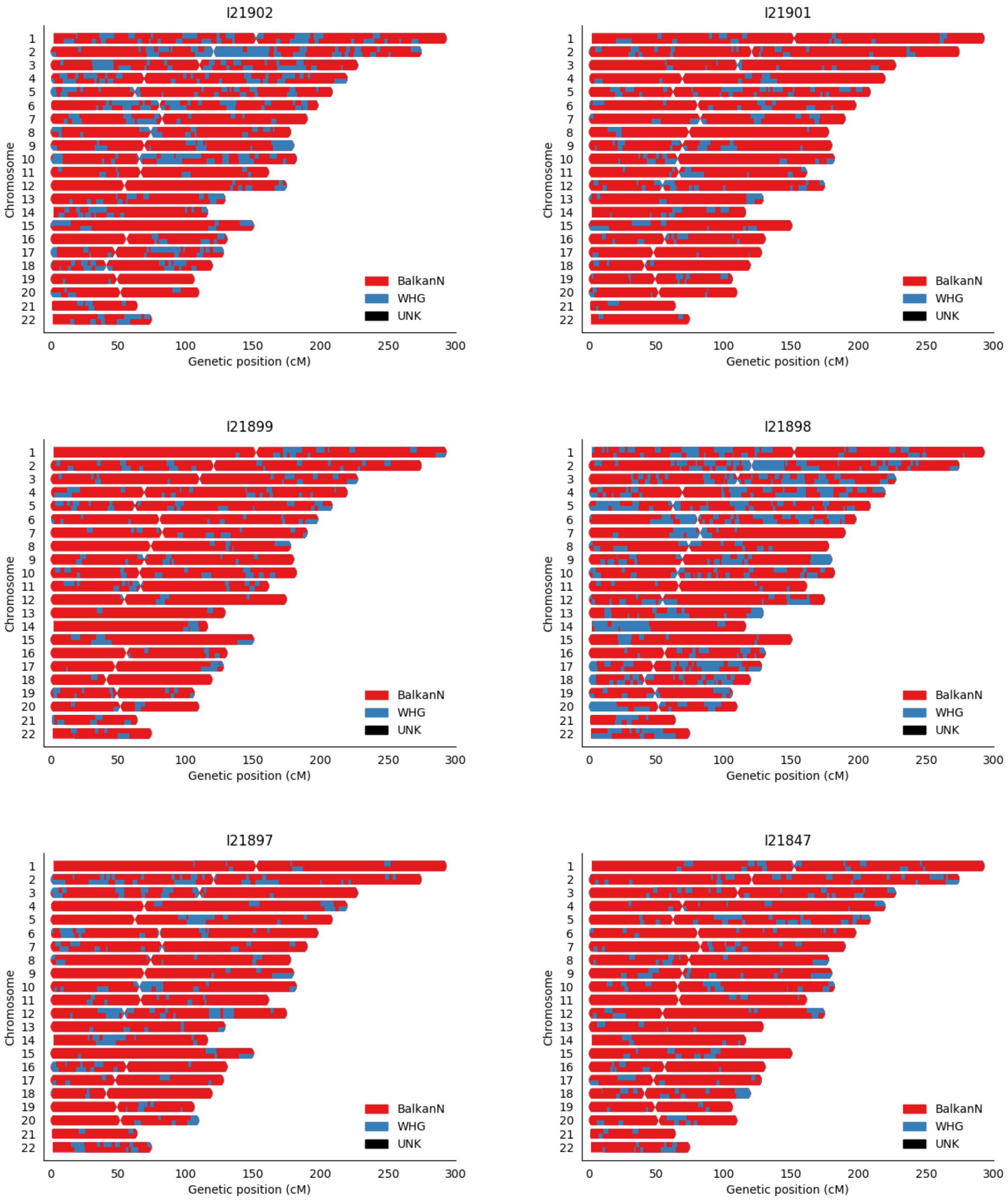
I21902
I21901
I21899
I21898
I21897
I21847
Chromosome
Genetic position (cM)
BalkanN
WHG
UNK

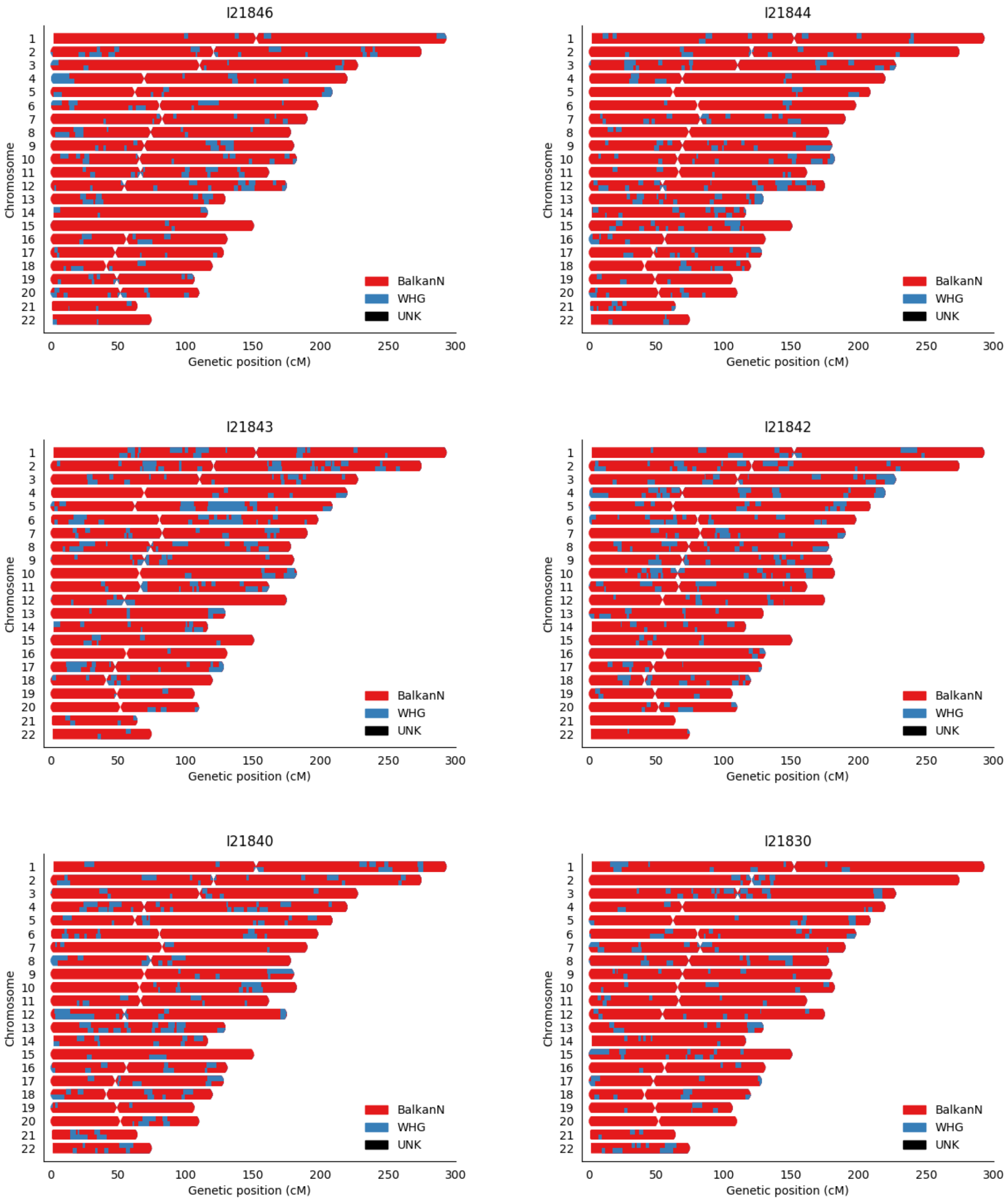
I21846
I21844
I21843
I21842
I21840
I21830
Chromosome
Genetic position (cM)
BalkanN
WHG
UNK

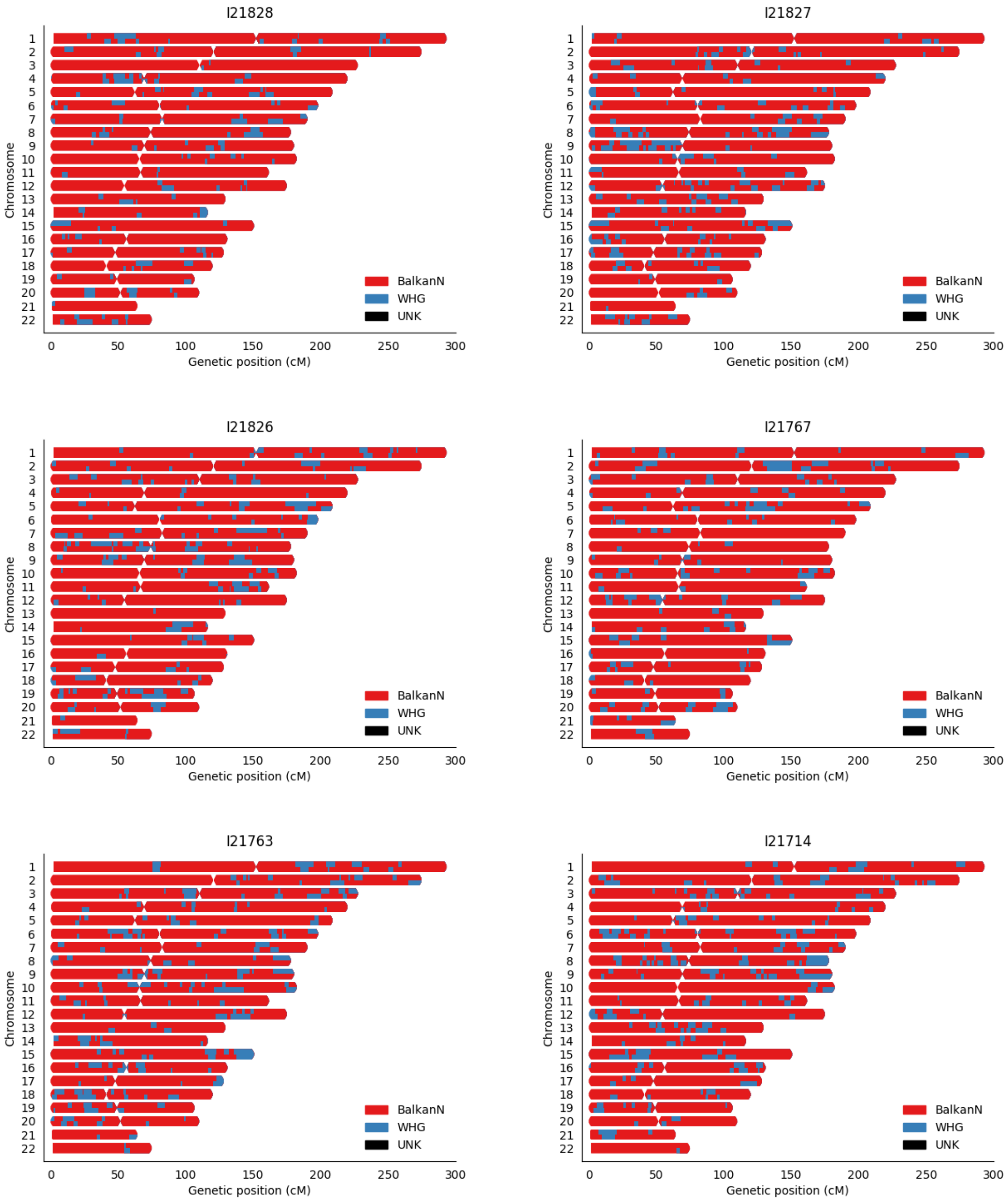
I21828
I21827
I21826
I21767
I21763
I21714
Chromosome
Genetic position (cM)
BalkanN
WHG
UNK

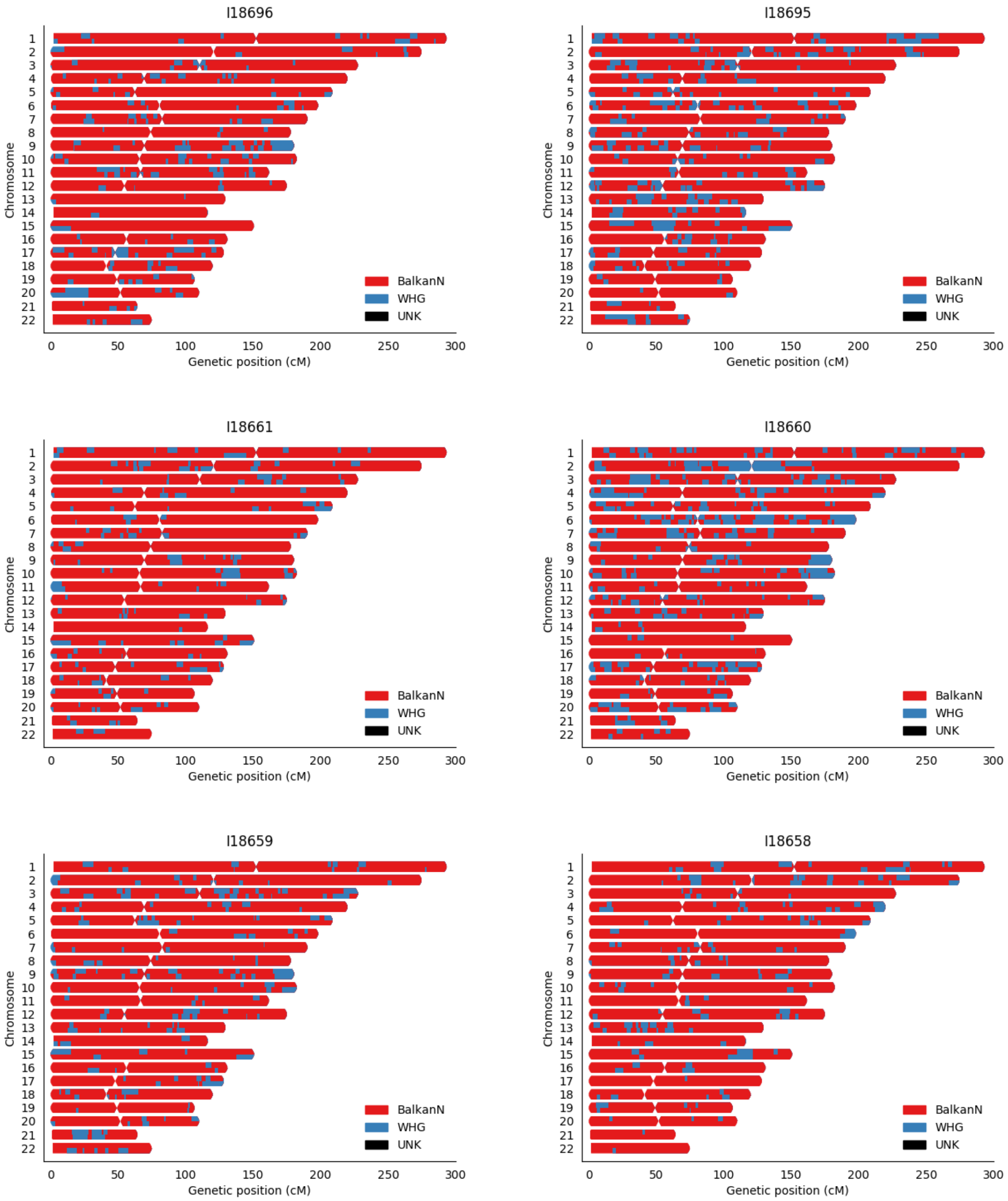
I18696
I18695
I18661
I18660
I18659
I18658
Chromosome
Genetic position (cM)
BalkanN
WHG
UNK

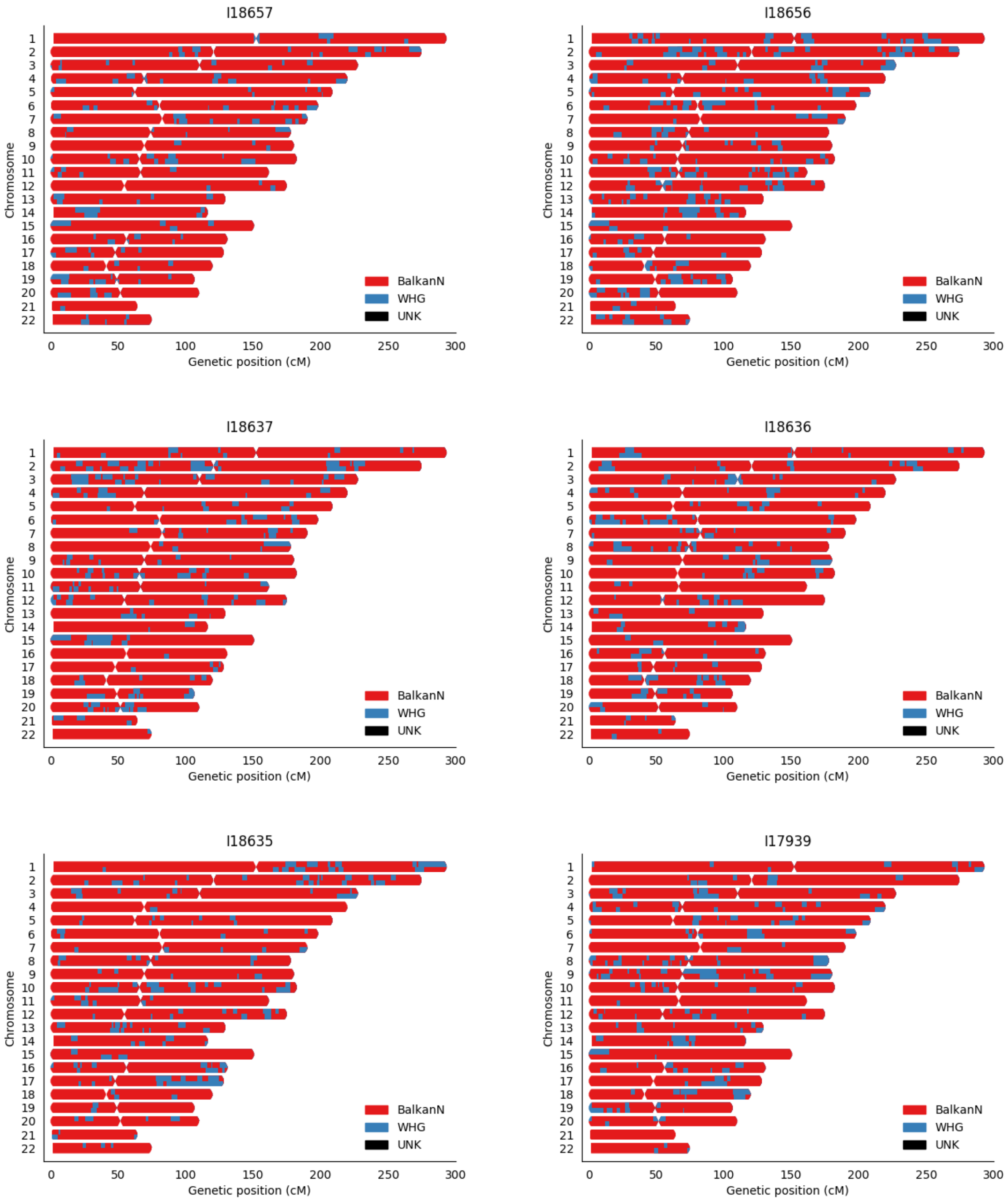
I18657
I18656
I18637
I18636
I18635
I17939
Chromosome
0
50
100
150
200
250
300
Genetic position (cM)
BalkanN
WHG
UNK

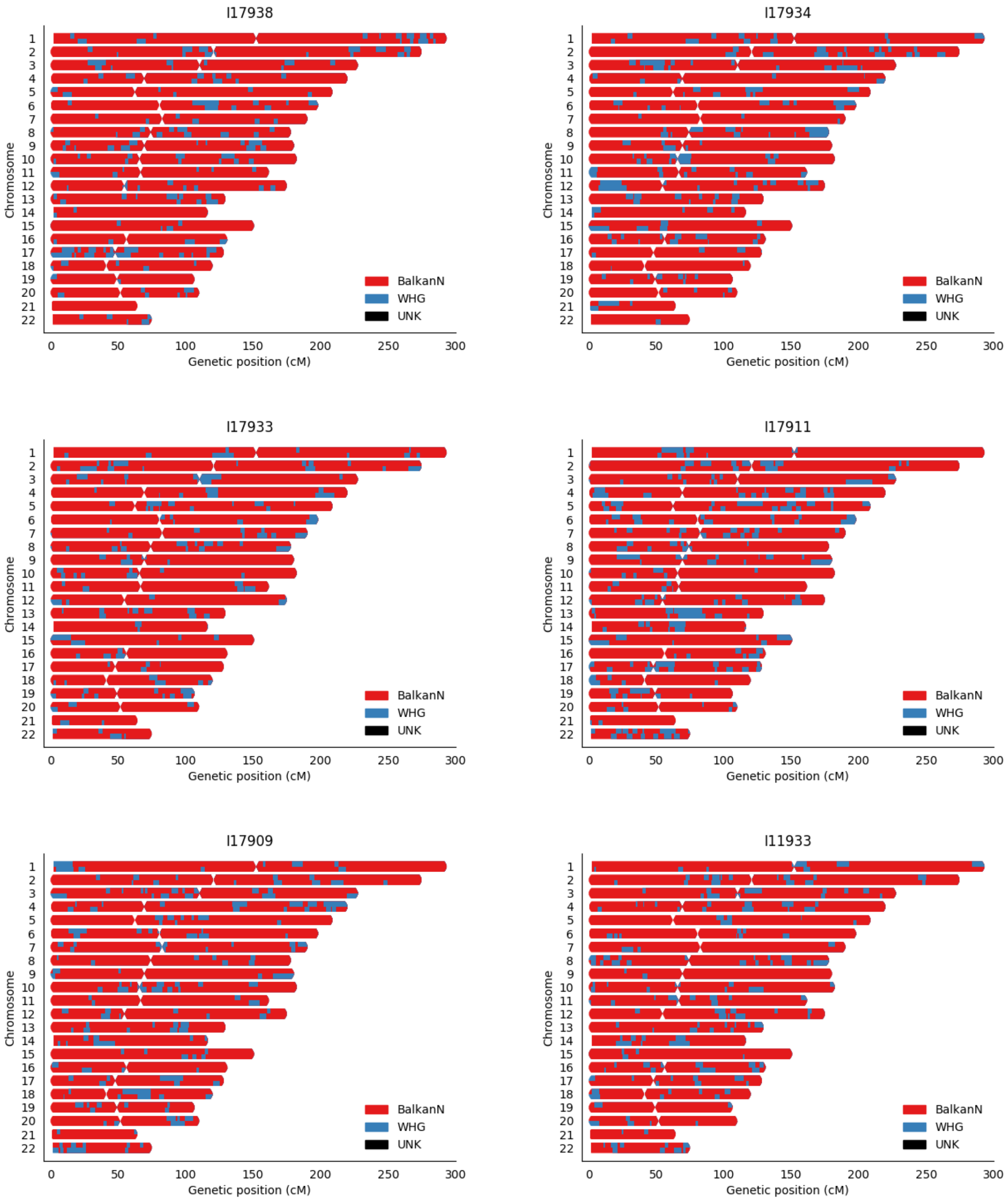
I17938
I17934
I17933
I17911
I17909
I11933
Chromosome
Genetic position (cM)
0
50
100
150
200
250
300
BalkanN
WHG
UNK

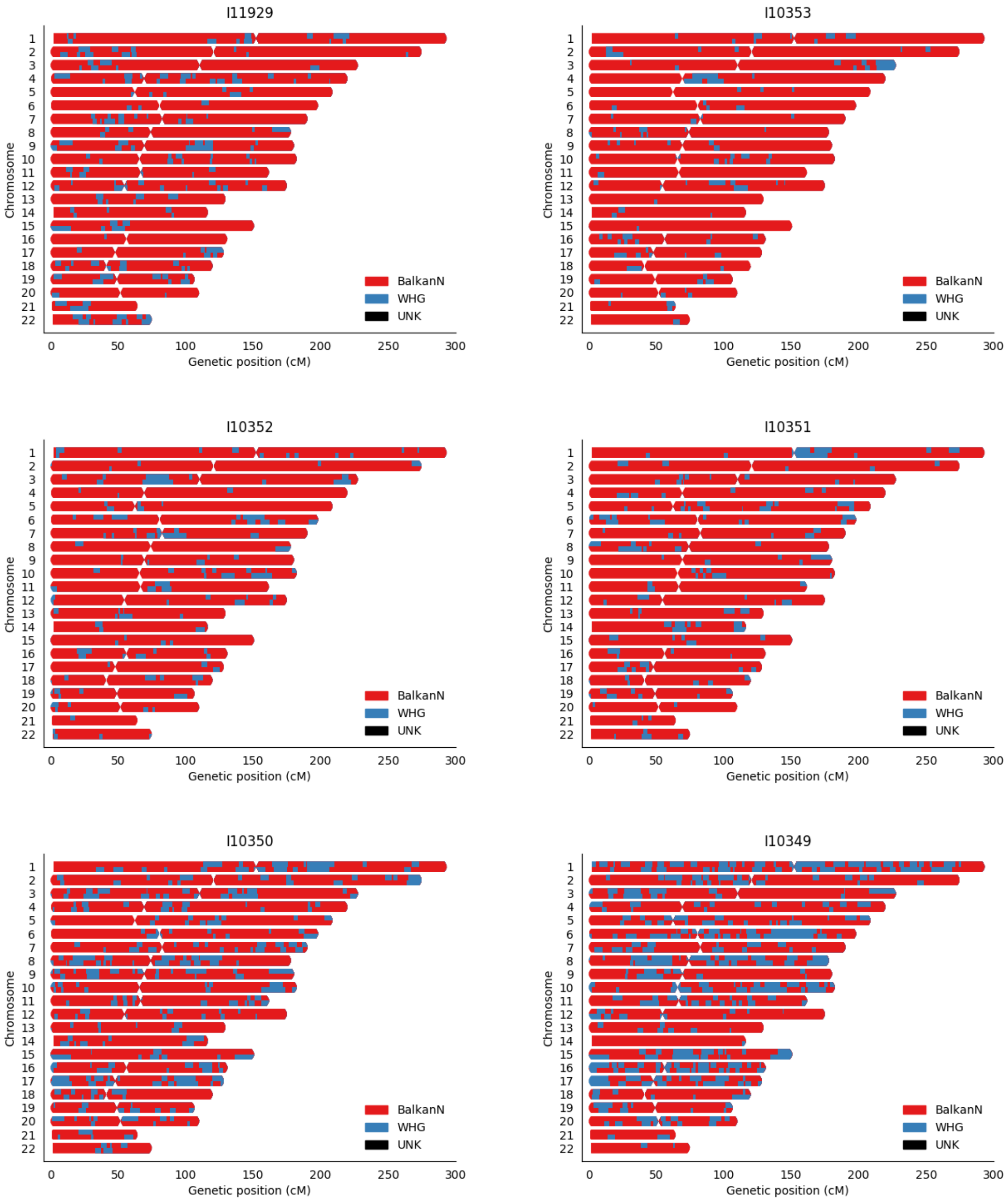
I11929
I10353
I10352
I10351
I10350
I10349
Chromosome
Genetic position (cM)
BalkanN
WHG
UNK
0
50
100
150
200
250
300

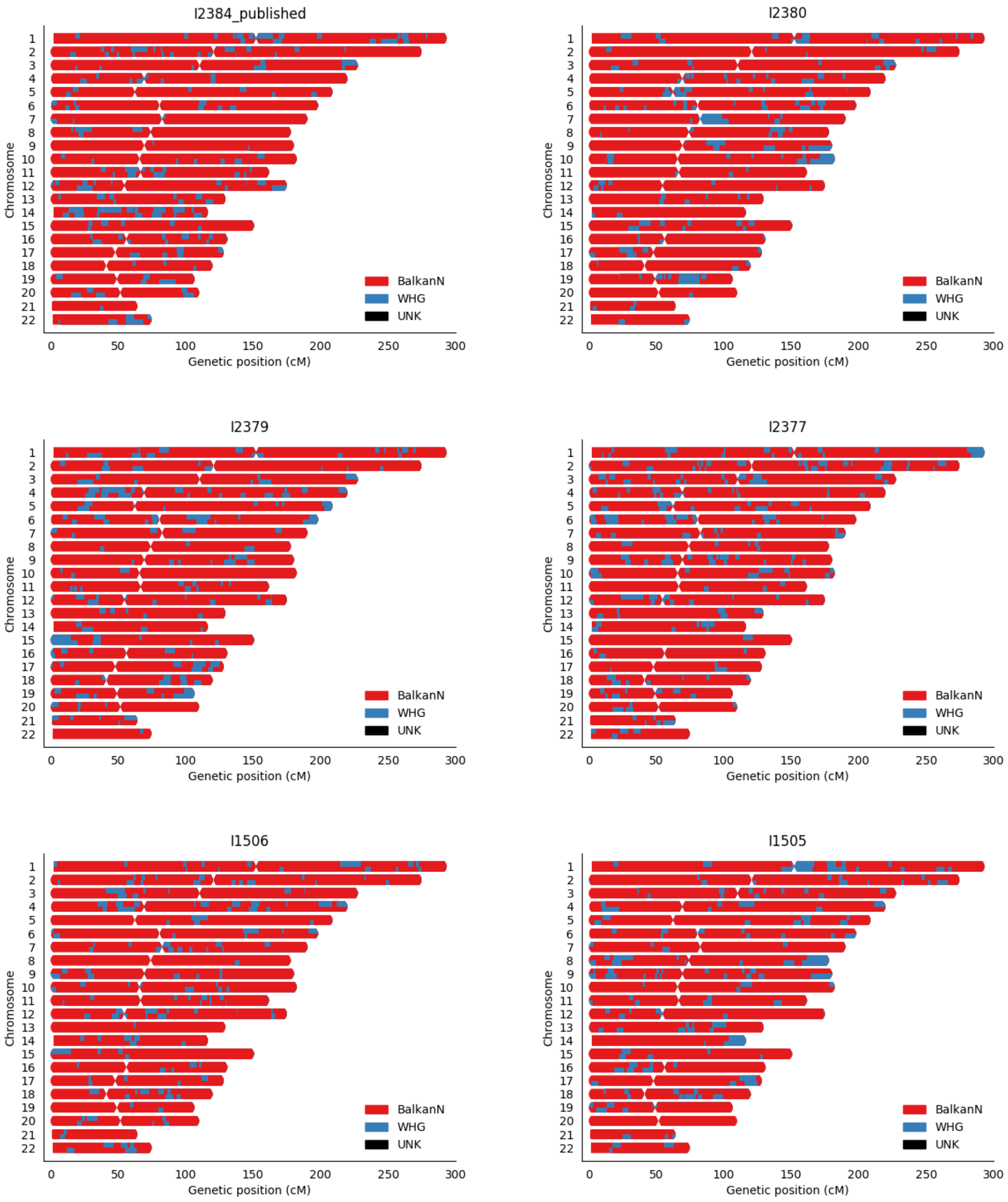
I2384_published
I2380
I2379
I2377
I1506
I1505
Chromosome
Genetic position (cM)
BalkanN
WHG
UNK

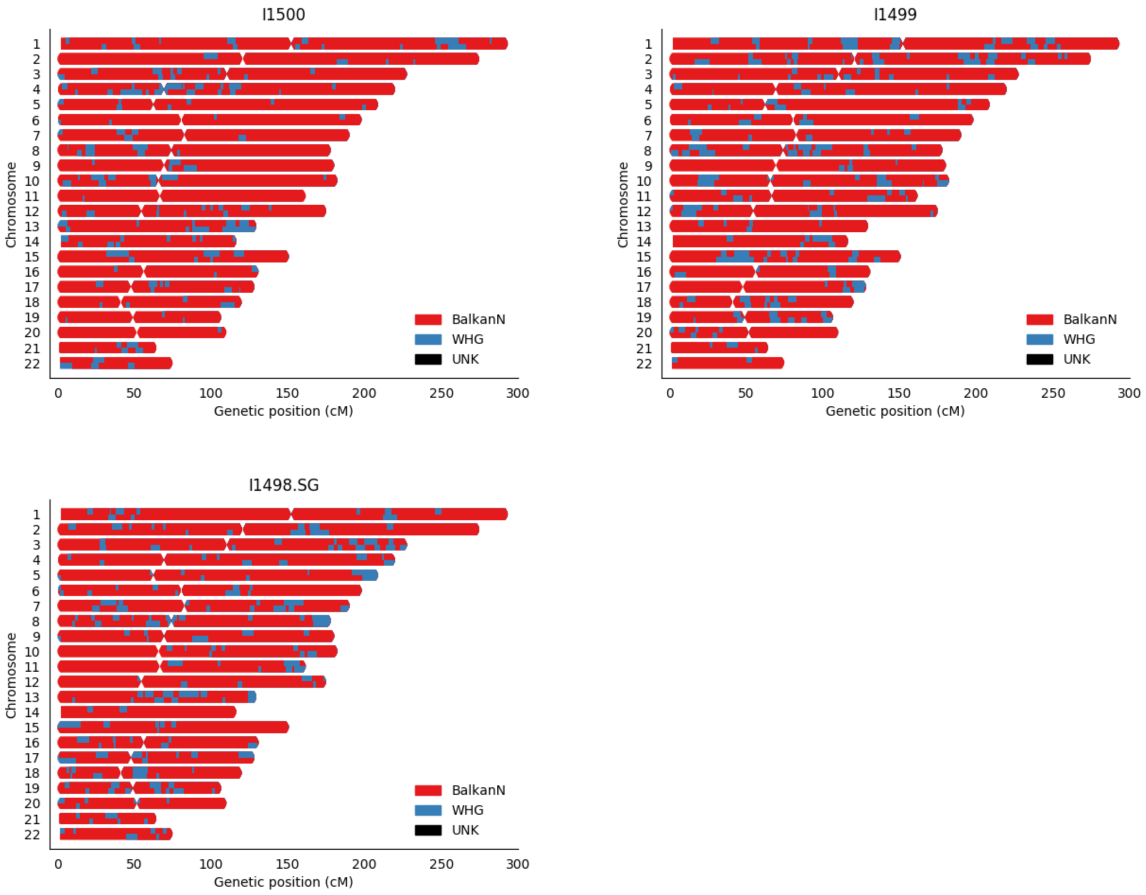
I1500
I1499
I1498.SG
Chromosome
Genetic position (cM)
BalkanN
WHG
UNK

**Supplementary Figure 3:** Regression of RFMix and qpAdm WHG ancestry estimates in ALPC individuals.

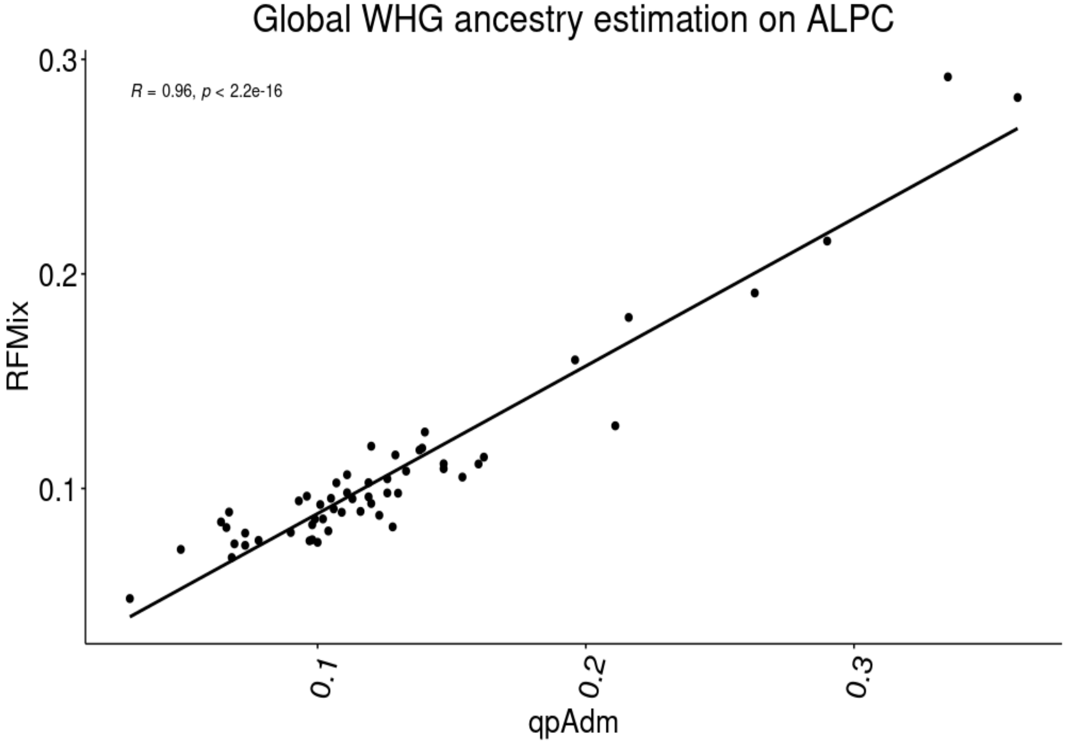

**Supplementary Figure 4:** Distribution of the Y chromosome and mtDNA haplogroups per population. The Y-axis represents the number of individuals.

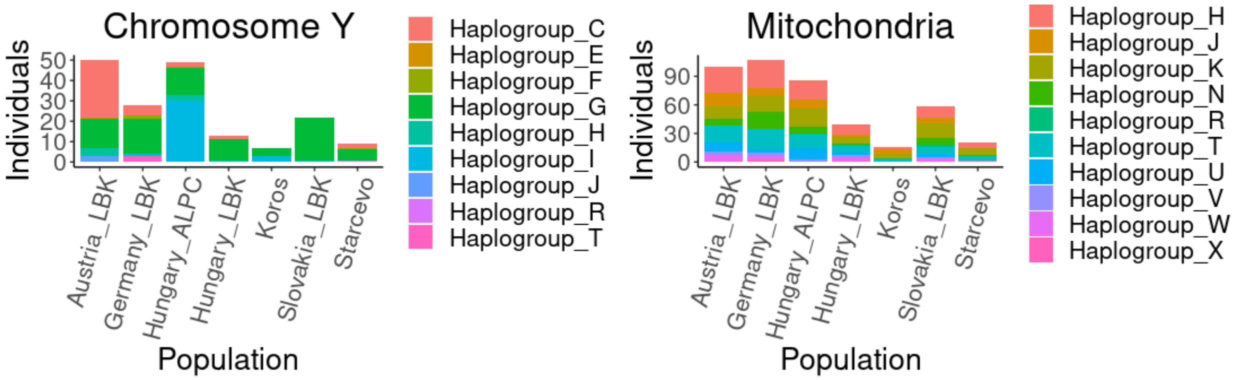

**Supplementary Figure 5:** Isotope data from Pólgar-Ferenci-hát. Here we plot the ratio δ13C/δ15N. Each dot represents one individual and the colour denotes the family.

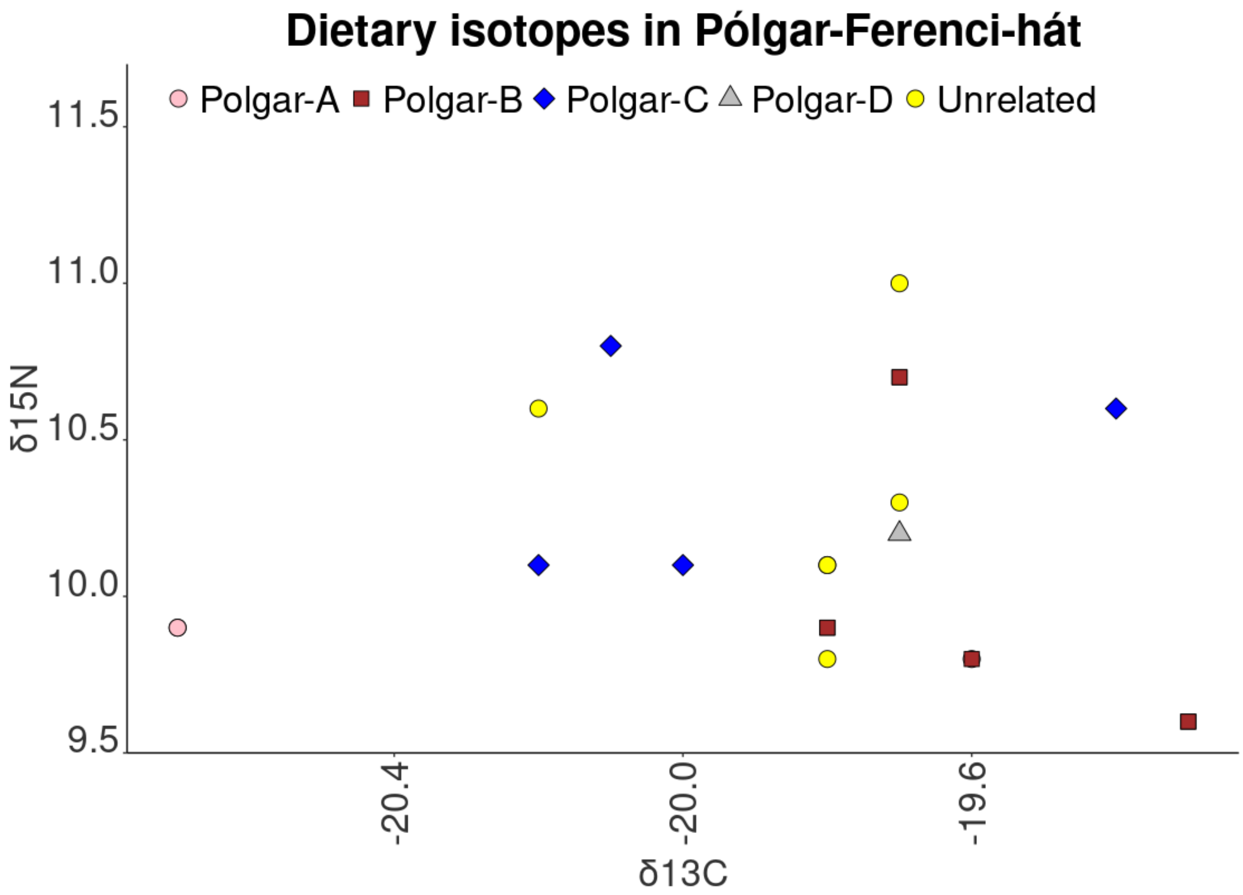

**Supplementary Figure 6:** Individuals with more than 400,000 SNPs and the assessed ROH. Individuals in the ALPC group show a higher rate of close-kin unions (as reflected in the presence of ROH segments >20cM) than the rest of the dataset.

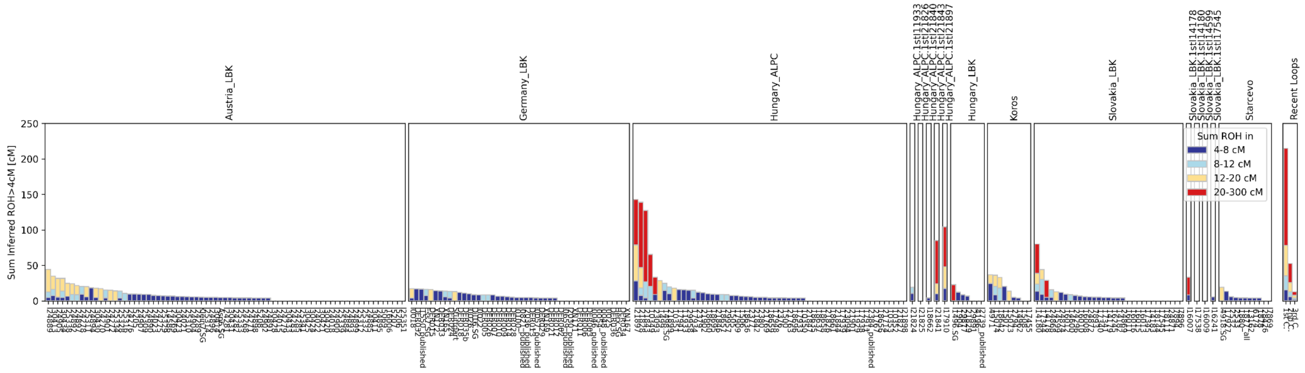

**Supplementary Figure 7:** *Tests for positive selection Cand long-term balancing selection in the ALPC and LBK population, made with the qqman [1]. The red lines indicate the top 0.05% cutoff. (A) Normalized iHS scores in ALPC. (B) Normalized iHS scores in LBK. (C) Normalized unphased iHS scores in ALPC. (D) Normalized unphased iHS scores in LBK. (E) Normalized nSL scores in the ALPC. (F) Normalized nSL scores in LBK. (G) Normalized unphased nSL scores in ALPC. (H) Normalized unphased nSL scores in LBK population.*

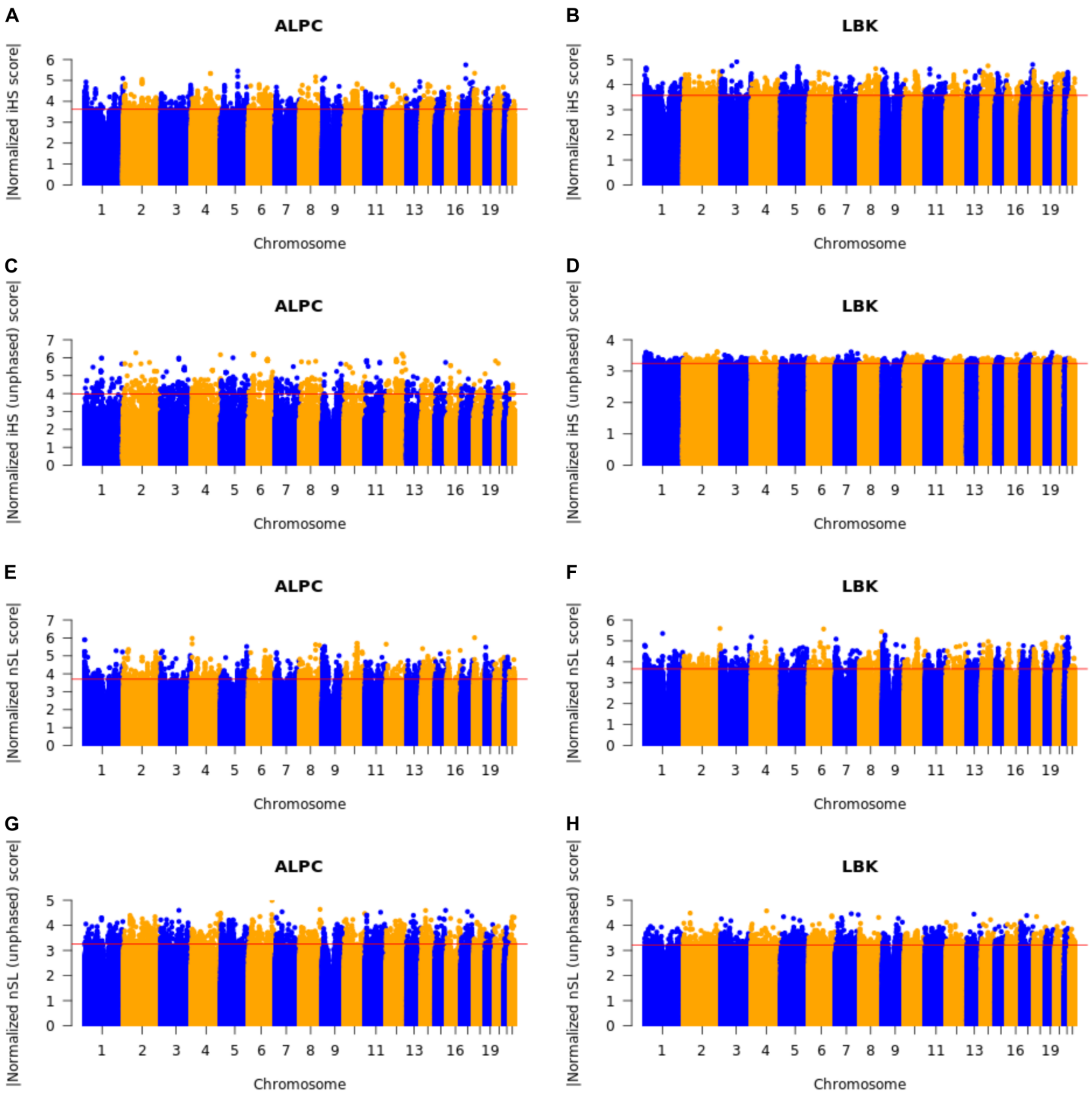

**Supplementary Figure 8:** Correspondence between the ancestry in ALPC segments with the selection scan values. Each dot represents a region of 0.2 cM of the genome, in the Y-axis we display the average WHG ancestry of the region, and in the X axis the average selection coefficient of the region. We show the Spearman correlation coefficients and p-value.

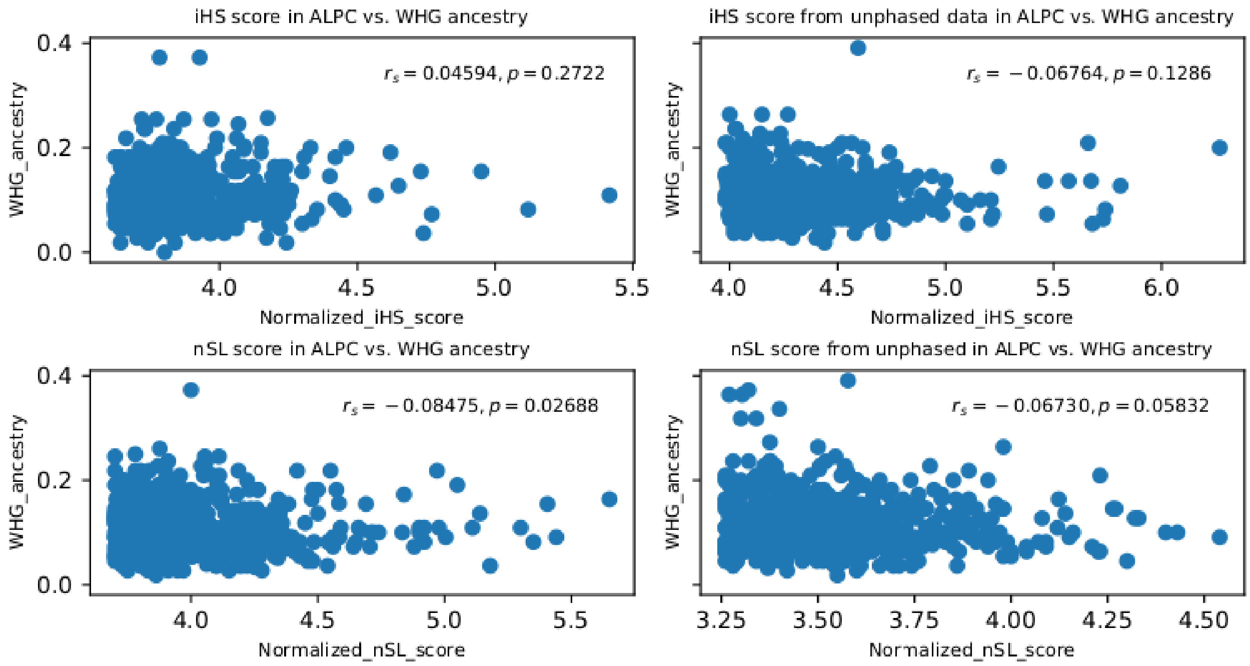

### Section 2: Kinship

**Authors:** Iñigo Olalde, Pere Gelabert

The complete kinship results are displayed in Table S7.

#### Description of the Methodology

To reconstruct the pedigrees and family relationships, we followed the methodology described by [2]. This methodology reconstructs the kinship connections based on two estimators. A) The first is that relatedness coefficient *r* indicating the relationship between individuals based on comparing the observed autosomes mismatches between a pair of individuals and the expected rate for unrelated individuals. B) The second is the results from ngsRelate [3] which uses genotype likelihoods and population allele frequencies to estimate Cotterman coefficients KO, K1, and K2. These coefficients correspond to the probability of sharing 0 1 or 2 alleles by identity by descent (IBD), estimating the degree of relationship. Finally, we used the allelic mismatch rate values across sliding windows of 20 Mb, moving by 1 Mb to differentiate between parent-offspring and sibling relationships. We also took into consideration sex, mtDNA haplogroup, Y chromosome haplogroup, and age to reconstruct the.

We plotted the results in pedigrees. A circle denotes a female, and a square a male. Inside each circle, there are four lines: genetic ID, grave ID (in parenthesis), mtDNA haplogroup, and anthropological age. Inside each square, there are five lines: genetic ID, grave ID (in parenthesis), Y-chromosome haplogroup, mtDNA haplogroup, and anthropological age. A question mark inside a circle means that the individual is not sequenced. A line with dots denotes a relationship that cannot be drawn in the pedigree due to lack of information.

#### 2.1 Nitra

**Nitra family A**

I16007 (female) and I14178 (male) are siblings according to the K1 value. I11873 is 2nd or 3rd-degree relative of them, showing a recombination pattern similar in both cases and very fragmented, which would be consistent with being a paternal aunt. I18143 and I11873 are 2nd or 3d degree relatives, but the relationship's nature is unclear due to low coverage. I11873 is a young woman, and the two other individuals (I16007, I14178) are children who show indications of head trauma and are buried in the same grave. Both children show elevated ROH (33 and 28 cM in regions >4 cM) compared to the rest of the individuals from Nitra, compatible with inbreeding.

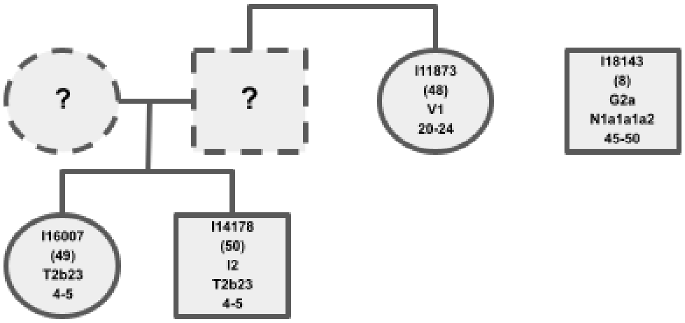

**Supplementary Figure 9:** Pedigree of Nitra family A.

**Nitra family B**

I16246 is the mother of I16008 according to the K1 values.

**Nitra family C**

I25175 is a 3rd-degree relative of I18105. No further information can be added.

**Nitra family D**

I14599 is the mother of I16009 (female), according to K1 values. I17339 is a 3rd degree relative of I16009, probably on the paternal side. I14599 shows low strontium values, and I16009 and I17339 have average strontium values. I16009 is also a 3rd-degree relative of I25179. I14599 is estimated to be a fourth-degree relative of I16010.

**Nitra family E**

I14180 is the father of I17538, consistent with the K1 value, and both are 4$^{th}$-degree relatives of I14183 and I14600. I14600 and I18091 are 1$^{st}$-degree relatives. Possibly I14600 is the son of I18091. I18144 and I18091 are sister and brother, and I16011 seems to be their nephew. Looking at the IBD plots, I14183 is differently related to I14600 and I14183, indicated by different recombination events. I18097 is a 3$^{rd}$-degree relative of I18091 and a 4th-degree relative of I18144; this individual is not displayed in the pedigree as the relationship is unknown.

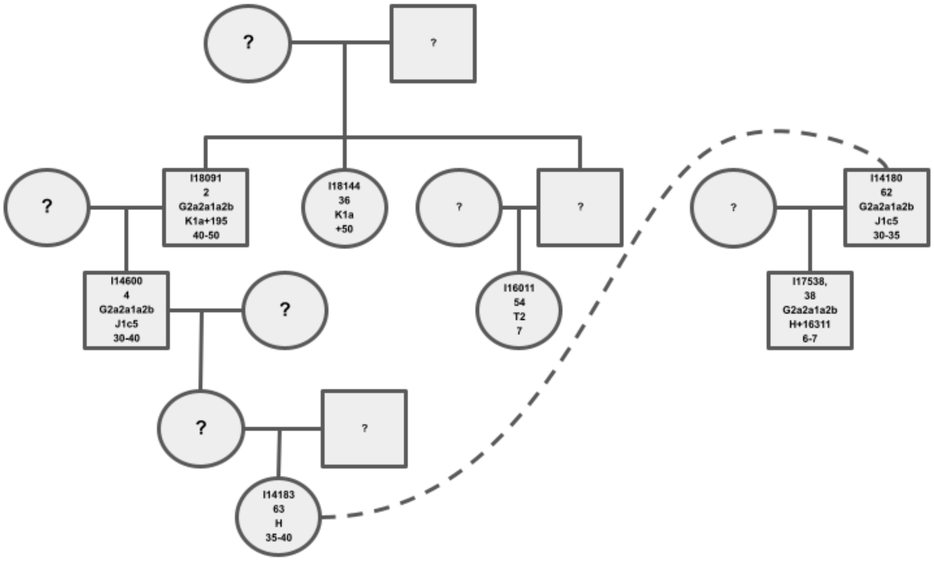

**Supplementary Figure 10:** Pedigree of Nitra family E.

**Nitra family F**

I16013 and I16015, both females and are second-degree relatives. Probably they are half-sisters or aunt-niece.

**Nitra family G**

I16245 and I14177 are 2nd or 3d degree relatives, both males. Both individuals have long ROH segments and different mtDNA haplogroups.

**Nitra family H**

I17345 and I16016 have a 3rd-degree relationship, and both are males. I17344 and I17345 are fourth-degree relatives.

**Nitra Family I**
I14179 and I25176 are 3rd-degree relatives.

**Nitra Family J**
I17545 and I16241 are daughter and mother

#### 2.2 Pólgar-Ferenci-hát

**Polgár-Ferenci-hát Family A**

Family A from Polgar-Férneci-hát consists of 13 individuals. I17910 and I21897 are siblings based on the *r* value and low K1 value. I17910 and I17909 share long IBD and are inferred to be 2nd-degree relatives (I17909 was buried with 2 Spondylus beads). They may have been half-brothers, which is also consistent with the relationship between I17909 and I21897. I21824 and I11933 are 3rd-degree relatives of both I17910 and I21897, but not of I17909, which indicates that the relationship must be on the maternal side which is also consistent with the mtDNA haplogroup. These patterns would be coincident with their being cousins. I11933 and I21824 are brother and sister according to the K1 and the r_X. I21769 is a 3rd-degree relative of I17910 and I21897 and a fourth of I11933. I11929 is a 3rd-degree relative of the father of I17909, I21897, and I17910, but it is unclear which relationship they share. I21843 is the father of I21765 according to the K1 value and the clear similar profile in the X chromosome. This individual is also the brother of I21841 according to the K1 values. Individual I21767 is the nephew of I21841 as he shares a 2nd-degree relationship with I21841 and I21843. Individual I21900 is compatible with being the paternal aunt of I21841 and I21843, but the evidence is not unambiguous. I21901 is inferred to be a 3[rd] or 4th degree relative of I21843, I21841, and I21765. I21841 and I21843 are likely offspring of 1st degree relatives, the same as I21897 and I21910.

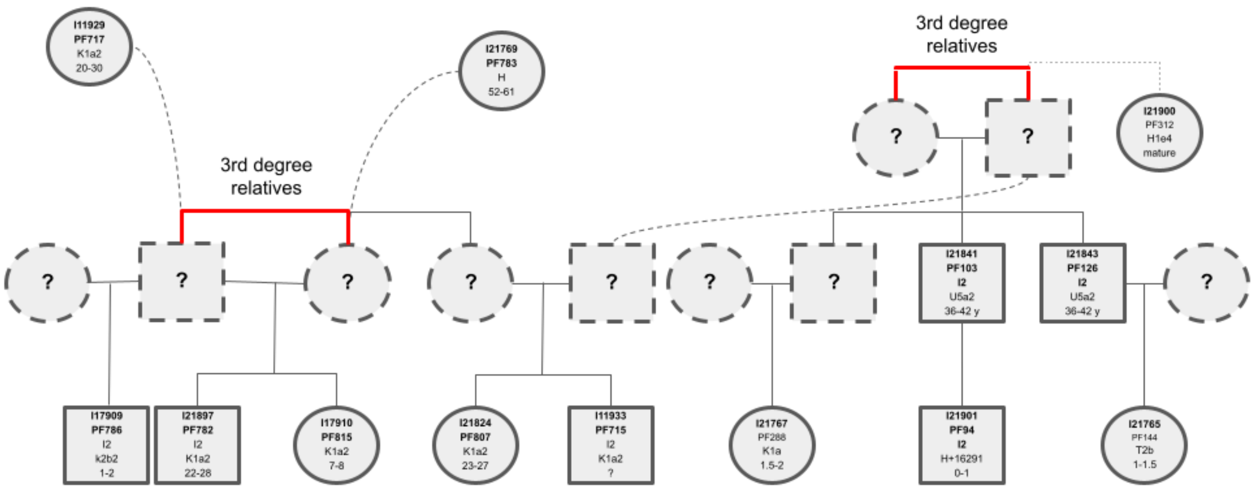

**Supplementary Figure 11:** Reconstructed pedigree of Pólgar-ferenci-hát family A.

The burials of this family are physically in close proximity at the site. Individuals I17909 and I17910 are buried together and in the same area as individual I21879. Individuals I21824 and I11933 are siblings and are also in the same area. The other nucleus of this family, composed of individuals I21841, I21843, I21901, I21765, and I21767, is located in a separate area. The first group (I17909, I17910, I21879, I21824, and I11933) are buried in the ALPC-I area, while I21841, I21843, I21901, I21765, I21767 are located in the ALPC II-IV areas, showing connections between individuals in different phases of Pólgar-Ferenci-hát.

**Pólgar-Ferenci-hát family B**

I21898 is the father of I21902 (male) and I18660 (female). I21898 is a 4th-degree relative of I18695, who is the daughter of I21827. I18695 and I21898 are also 4th-degree relatives of I17911, although the exact nature of that relationship is unclear.

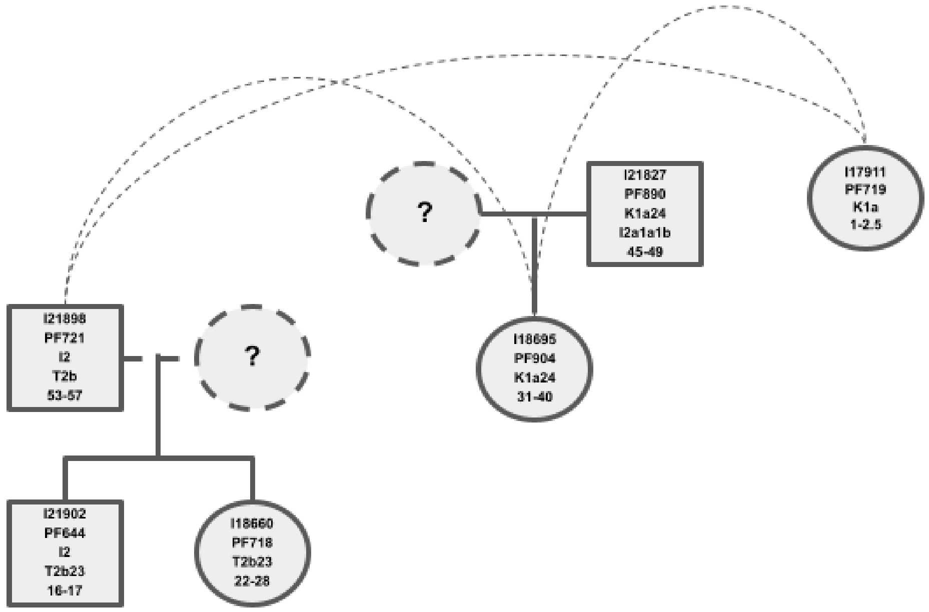

**Supplementary Figure 12** Pedigree of Pólgar-ferenci-hát family B.

Individuals I21898, I21902, and I18660 are buried together, and individuals I18695 and I21827 are buried together and not far from the other ones and also close to I17911. All these individuals are located in the ALPC I area. Individual I18660, which corresponds to PF718 burial, is buried with a Spondylus clay bead and a cattle figurine, which is not common for the period of the burial. Individual I17911 has a lot of small Spondylus beads. In the burials which could imply membership of a privileged group [4]

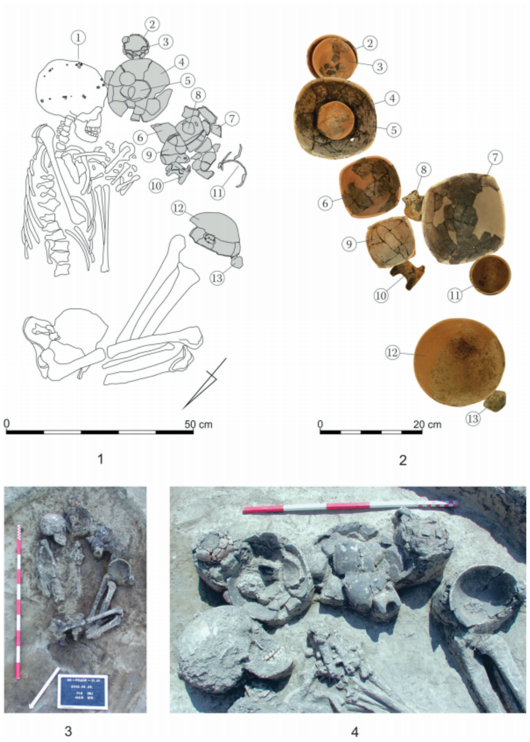

**Supplementary Figure 13:** Burial of individual I18660 with the funerary materials found in the grave, image from [4].

**Polgár-Ferenci-hát family C**

I21840 is the mother of I18662 (male) and the grandmother of I21768 (male) and I21844 (male). I18662 is the grandfather of I18659 and the uncle of I21844. I18659 is compatible with being the grandson of I18662. Individuals I21847, I18862, I21768, and I18659 are all located very close to each other. Individual I21844 is located far from all. Far away from these, individuals I21825, I21826, and I18658 are all close together in the ALPC II-IV area. I18657 is compatible with being a 3rd-degree relative of I21847.

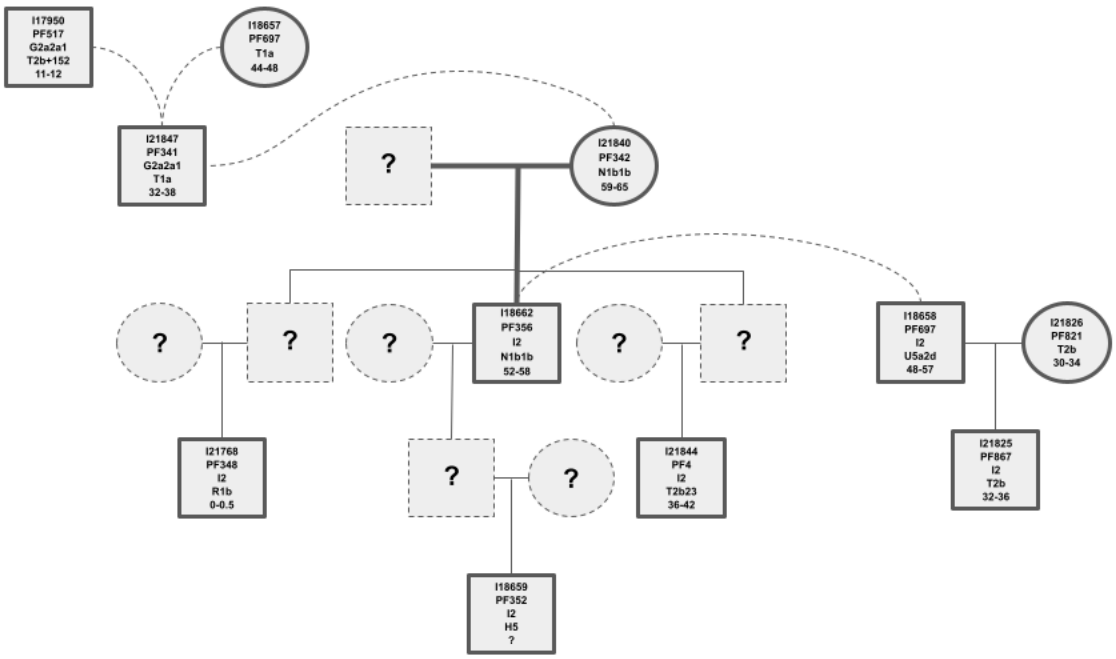

**Supplementary Figure 14:** Pedigree of Pólgar-ferenci-hát family C.

**Polgár-Ferenci-hát family D**

I18656 and I21842 are 2nd-degree relatives and show no similarity at all on the X chromosome, implying that the relationship must be paternal-sided. They also share the same Y chromosome haplogroup and a different mtDNA one. These individuals are compatible with being: half-paternal brothers, uncles and nephews, or grand-father grand-son. I18696 (adult male) is a 3rd degree relative of I18656, but with no evidence of relatedness to I21842, which could be evidence of the first two being half-brothers or grandfather and grandson. I18696 shows a possible 4th-degree relationship with I21826, who is a member of Family C. The graves of individuals I21842 and I18656 are closely located in the site.

#### 2.3 Asparn-Schletz

**Family A**

I30431 (FN 162, probably young adult) and I24892 (1993/4, FN 4464, FN 4518, probably 45-55 years are inferred to be son and father respectively, both from the ditch. They also have a 4th degree relationship to I24017 (FN 9230, 2.5-3 years) from the settlement.

**Family B**

I24278 (1985/ind 57, FN 249, 1-2 years) and I27800 (FN 6950, 2-3 years) are both from the settlement area, and genetically inferred to be brothers.

**Family C**

I30411 (FN 10343, 2-4 years) and I24016 (FN 9366, 4-5 years) are both from the settlement (close to the ovens), and are genetically inferred to be brothers.,

**Family D**

I24906 (2001, FN 10806, 2-3 yrs) and I25332 (2001, FN 10640, 9 months), are from a double burial of males in the settlement, and are siblings.

**Family E**

I24272 (2005, FN 14143, 2-3 years) and I24884 (1996/2, FN 5076, 35-45 years) are 2nd-degree relatives and share part of the X chromosome; 1996/2 could be the maternal grandfather.

**Family F**

I24021 (burial 16, 2000, 2-3 years) and I24904 (1999/3, FN 8328, 1.5-2.5 years) are both from the settlement and probably from the enclosing ditch system. They show the same Y Chromosome (C1a2) and different mtDNA, and could be half-brothers.

**Family G**

Individual I25347 (1991/1992, Fn. 3991 or 3491, age at death unclear) has a 3rd degree relationship with I24269 (2001, S 33, FN11660, 3-4 years), I24889 (1993/20, FN 4529, 2-3 years), and I24894 (1993/6, FN 4462, FN 4456, 3.5-4.5 years). All are located in the ditch 2 in the west, and the probability reflects casualties from a violent attack.

#### Section 3: Integration of genetic and isotopic data

**Author:** Penny Bickle

The isotopic data analyzed here was carried out as part of the LBK lifeways project [5]. The volume includes the full data, including methodology [6], results [5,7], and quality controls ([5], Appendix A). Further comparison with wider isotopic data from the LBK is available in [8]. All three sites were analyzed in the study but to different degrees. There were only five samples from Asparn Schletz that were analysed for both isotopes and aDNA, so no further comparison was carried out.

Our analyses asked two questions, enabled by comparison of the newly reported genetic data to the already collected isotopic data.

(1) Were there statistical differences in the means and variance of the isotope data when relatives (up to a third-degree) and non-relatives were compared? Due to the differences demonstrated between the sexes in previous studies we also analyzed related and unrelated individuals by biological sex.

(2) Did "family" groupings demonstrated different means and variances? As family groups are likely to have non-biological relatives, we interpret any of these differences as roughly indicating that kinship differences shaped diet and mobility.

Three main isotopes were analyzed, carbon, nitrogen, and strontium, with total strontium concentration in the tooth also studied. Carbon δ13C and nitrogen δ13N provide information about dietary protein consumption, with nitrogen isotopes varying with trophic level and hence meat protein consumption. Carbon isotopes are likely to primarily reflect variation in forest canopy cover in an LBK context (rather than consumption of C3 vs. C4 plants as at most later times) [8,9]. Although some freshwater fish consumption may have occurred in the LBK, it is unlikely to have significantly impacted on stable isotope variation [10]. Strontium isotopes, analysed in tooth enamel, are sensitive to lifetime mobility when the Sr ratio from the tooth differs from that of the local geology [11].

We used Levene's test for assessing homogeneity in variance. This is particularly significant for analysing strontium isotope ratios where differences between groups are likely to speak to more or less mobility. As is frequent practice in the statistical analysis of isotopic data in archaeology, we make the assumption that outliers result in non-normal distributions of the data. As non-parametric tests use few assumptions, we also report those tests approaching significance or those that are close to but slightly above $p=0.05$. Thus, we used the Mann-Whitney U test when comparing the results by biological sex and the Kruskal-Wallis between the family groupings. Infants showing a weaning signature by having elevated nitrogen levels were excluded from the analysis (1 for Nitra, and 3 for Polgar).

**Nitra**

A total of 45 individuals from Nitra produced reliable results for aDNA and carbon, nitrogen, and strontium isotope ratios. Not all of these 45 individuals had both strontium and stable isotope values, mainly due to no teeth being preserved, but in some cases because of poor collagen preservation of the ribs which were sampled for the stable isotope analysis. This meant that some comparisons could not be carried out.

The published isotopic results from Nitra suggested a small but significantly elevated average nitrogen isotope ratio for male adults over the rest of the population (Whittle et al. 2013a, 150), potentially suggesting a high protein or meat content in their diet. Adult females had significantly more variable strontium isotope ratios than men, and accounted for all the outliers (Whittle et al. 2013a, 152).

* Significant ($p<0.05$)
** Approaching significance (following Lifeways $0.05<p<0.1$)

**Levene's test for significance difference in variance in Nitra:**

| Groups compared | Isotope | N | Levene statistic | Sig. |
|---|---|---|---|---|
| Related (all) and unrelated (all) | Sr ratio | 34 | 0.01 | 0.975 |
| | Sr conc | 32 | 0.279 | 0.601 |
| | d13C | 35 | 2.328 | 0.137 |
| | d15N | 34 | 0.326 | 0.572 |
| Related males and females | Sr ratio | 20 | 3.121 | 0.093** |
| | Sr conc | 20 | 1.516 | 0.233 |
| | d13C | 25 | 0.1.232 | 0.278 |
| | d15N | 25 | 0.172 | 0.682 |
| Unrelated males and females | Sr ratio | 12 | 1.466 | 0.249 |
| | Sr conc | 10 | 4.707 | 0.055** |
| | d13C | - | | |
| | d15N | - | | |
| Related females and unrelated females | Sr ratio | 20 | 0.164 | 0.689 |
| | Sr conc | 19 | 0.414 | 0.528 |
| | d13C | 23 | 3.188 | 0.087** |
| | d15N | 23 | 2.832 | 0.095** |
| Related males and unrelated males | Sr ratio | 12 | 0.002 | 0.966 |
| | Sr conc | 11 | 13.942 | 0.003* |

| | | | | |
|---|---|---|---|---|
| | d13C | - | | |
| | d15N | - | | |
| Related males and unrelated females | Sr ratio | 18 | 3.702 | 0.07** |
| | Sr conc | 17 | 1.151 | 0.298 |
| | d13C | 18 | 0.179 | 0.677 |
| | d15N | 17 | 0.666 | 0.426 |
| Related females and unrelated males | Sr ratio | 14 | 1.208 | 0.290 |
| | Sr conc | 13 | 0.544 | 0.474 |
| | d13C | - | | |
| | d15N | - | | |
| Family grouping | Sr ratio | 12 | 9.220 | <0.001* |
| | Sr conc | 12 | 6.957 | 0.003* |
| | d13C | 16 | 1.374 | 0.277 |
| | d15N | 16 | 2.017 | 0.106 |

**Tests with significant results ($p<0.05$):**

1. Unrelated males and related males had significant variation in strontium concentration ($p=0.003$). This would be consistent with unrelated males having more variable origins in childhood.
2. Variance in family groupings are not equal across Sr isotope ratio ($p<0.001$) and Sr concentration ($p=0.003$). This suggests that "families" did not share the same mobility patterns.

**Mann-Whitney U tests for difference in means in Nitra:**

| Groups compared | Isotope | N | Mann-Whitney U | Sig. |
|---|---|---|---|---|
| Related (all) and unrelated (all) | Sr ratio | 36 | 148.5 | 0.86 |
| | Sr conc | 34 | 126.5 | 0.845 |
| | d13C | 37 | 134 | 0.987 |
| | d15N | 36 | 106 | 0.59 |
| Related males and female | Sr ratio | 22 | 40.5 | 0.203 |
| | Sr conc | 22 | 66 | 0.722 |
| | d13C | 27 | 118.5 | 0.134 |
| | d15N | 27 | 112 | 0.251 |
| Unrelated males and females | Sr ratio | 14 | 10 | 0.188 |
| | Sr conc | 12 | 11 | 0.727 |
| | d13C | 10 | 0 | 0.2 |
| | d15N | 9 | 8 | 0.222 |
| Related females and unrelated females | Sr ratio | 22 | 59.5 | 0.974 |
| | Sr conc | 21 | 53 | 0.972 |
| | d13C | 25 | 59.5 | 0.487 |
| | d15N | 24 | 71 | 0.697 |

| | | | | |
|---|---|---|---|---|
| Related males and unrelated males | Sr ratio | 14 | 28 | 0.304 |
| | Sr conc | 13 | 19 | 0.593 |
| | d13C | 12 | 10 | 0.333 |
| | d15N | 12 | 1 | 0.333 |
| Related Males and unrelated females | Sr ratio | 20 | 34 | 0.247 |
| | Sr conc | 19 | 44.5 | 0.968 |
| | d13C | 20 | 50.5 | 0.941 |
| | d15N | 19 | 65 | 0.091** |
| Related females and unrelated males | Sr ratio | 22 | 59.5 | 0.974 |
| | Sr conc | 21 | 53 | 0.972 |
| | d13C | 25 | 59.5 | 0.487 |
| | d15N | 24 | 71 | 0.697 |

**Kruskal-Wallis test to compare multiple groups:**

| Groups compared | Isotope | N | Kruskall-Wallis test statistic | Sig. |
|---|---|---|---|---|
| Family groups | Sr ratio | 22 | 11.098 | 0.196 |
| | Sr conc | 22 | 8.808 | 0.359 |

| | | | | |
|---|---|---|---|---|
| | d13C | 26 | 17.208 | 0.046* |
| | d15N | 26 | 10.135 | 0.34 |

**Tests with significant results:**

- Family groupings had different average carbon values to each other although the significance (*p*=0.046) is not compelling after correcting for multiple hypothesis testing. This may suggest that they were sourcing their food from different locations in the landscape.

Overall, the most notable observation is that family groupings are not unified in diet and mobility, and are significantly different in variation from each other.

**Pólgar-ferenci-hát**

A total of 48 individuals from Pólgar-Ferenci-hát produced reliable results for carbon and nitrogen, and 34 for strontium isotope analysis. Only 50% of the burials were analysed for isotopes by the LBK lifeways project (Whittle et al. 2013). In total, only 23 burials had both isotope and aDNA data, 17 related (6 females, 11 males) and 6 unrelated (5 females, 1 male). Due to low numbers of unrelated individuals, comparing the family groupings was the main focus for analysis.

In the published work, the main isotopic results from Pólgar-Ferenci-hát found no evidence of systematic dietary differences were suggested between the sexes, though women's nitrogen values increased as they aged (Whittle et al. 2013b, 84). Adult females had significantly more variable strontium isotope ratios than men, though this was perhaps less pronounced than at other sites (Whittle et al. 2013b, 86). It was hypothesized on the basis of variation in strontium isotopes between the molars that sex-based dietary differences became more pronounced between the ages of 8-12 (Whittle et al. 2013b, 87).

* Significant ($p$<0.05)

** Approaching significance (following Lifeways 0.05<$p$<0.1)

**Levene's test for significance difference in variance in PFH:**

| Groups compared | Isotope | N | Levene statistic | Sig. |
|---|---|---|---|---|
| Related (all) and unrelated (all) | Sr ratio | 15 | 0.33 | 0.574 |
| | Sr conc | 15 | 2.035 | 0.174 |
| | d13C | 21 | 1.244 | 0.277 |
| | d15N | 18 | 0.19 | 0.893 |
| Family grouping | Sr ratio | 12 | 16.747 | 0.001* |
| | Sr conc | 12 | 16.207 | 0.002* |
| | d13C | 16 | 21.767 | <0.001* |
| | d15N | 16 | 3.667 | 0.047* |

**Tests with significant results:**

- Family groupings all have different variances to each other across all isotopes.

**Mann-Whitney U tests for difference in means in PFH:**

| Groups compared | Isotope | N | Mann-Whitney U | Sig. |
|---|---|---|---|---|
| Related | Sr ratio | 17 | 32.5 | 0.792 |

| | | | | |
|---|---|---|---|---|
| (all) and unrelated (all) | Sr conc | 17 | 42 | 0.204 |
| | d13C | 23 | 40.5 | 0.460 |
| | d15N | 20 | 47 | 0.404 |

**Kruskal-Wallis test to compare multiple groups in PFH:**

| Groups compared | Isotope | N | Kruskall-Wallis test statistic | Sig. |
|---|---|---|---|---|
| Family groups | Sr ratio | 12 | 3.007 | 0.391 |
| | Sr conc | 12 | 1.157 | 0.763 |
| | d13C | 17 | 4.994 | 0.172 |
| | d15N | 15 | 1.451 | 0.694 |

**Conclusions for Polgar FH:**

- Family groupings are significantly different in variation from each other for mobility and diet, similar to the pattern at Nitra.

### Section 4: Site descriptions

##### 4.1 Bač, Topole (Serbia)

**Author:** Anđelka Putica

In 1977, systematic probe excavations were carried out at the sites of Šećerana and Topole next to Bač (Northern Serbia, Vojvodina Province) led by Čedomir Trajković (Town Museum of Sombor). Southeast of the town, in the area of the future sugar factory (Šećerana) four pits with material from the Eneolithic period were excavated (Baden/Kostolac ceramic fragments). The excavations were extended to the surrounding areas, between the railway and the Danube–Tisa–Danube channel as well as the area south of the Bač-Bačka Palanka road bounded by the river Mostonga (Topole). Here, in addition to some Eneolithic finds, Early Neolithic settlement objects, such as pits, the floor of a building structure, pottery vessels, clay statuettes, stone tools, as well as three contracted skeleton burials were revealed. On the basis of the findings, the Topole site has been dated to the late phase of Starčevo culture [12–14]. In Porbe I, Burial 1 (female, 20–25 years old) and Burial 2 (male, 40–50 years old) were uncovered beneath the floor of a structure of an irregular rectangular shape. The well-preserved skeletons were in contracted position 50 cm apart, at the same level, lying on their right sides, symmetrically back to back, with their heads pointing in opposite directions. At the skull of Burial 1 there was a fragment of a ceramic vessel. Next to Burial 2, there were fragments of Starčevo ceramics, a shell, and a chipped stone tool [12,13]. These burials were conserved *in situ* in their original position along with the soil around them, and transported to the Town Museum of Sombor. The calibrated AMS date for Burial 1 is 6216-5917 calBCE (7170±50 BP, OxA-8693) (95% confidence interval), while we ignore the direct date for Burial 2 (8085±55 BP, OxA-8504) [12,15]. As the archaeological context suggest simultaneous burials, and the burial customs are characteristic of the Early Neolithic period, the Mesolithic date of the Burial 2 samples which also is genetically consistent with being a first-degree relative of Burial 1 might be explained by a possible contamination of the Burial 2 sample during the conservation process [11]

- I7867: BACT_1, 6216-5917 calBCE (7170±50 BP, OxA-8693)
- I7868: BACT_2, 6000-5300 BCE

##### 4.2 Donja Branjevina, Deronje (Serbia)

**Author:** Branislav Vasov

Donja Branjevina is located in northwestern Serbia, next to the village of Deronje (Municipality of Odžaci) on an old alluvial terrace of the Danube, between the former Mostonga River (today the Danube–Tisa–Danube channel) and the Danube. The locality itself is located on a curve formed by the alluvial terrace, which is 4-6 m higher than the western part of the terrain, and thus a suitable place for settlement. The site was discovered in 1965, during the construction of the 2nd flood protection line, which was needed to protect against the flood caused by heavy rains and the overflowing of the Danube that year. Immediately after the flood, as well as the following year, probe excavations were conducted here under the leadership of local teacher and amateur archaeologist Sergej Karmanski. Afterward, archaeological research continued in the period from 1986 to 1996, which started again in 2020 and continues until today. In addition to the Neolithic, Late Bronze Age and Medieval horizons have also been confirmed at this site. From the Neolithic period, settlement objects, such as pit houses and trash pits were documented. In these objects and the layers above them, the most common finds were fragments of ceramic vessels with typical early Neolithic paint or ornaments. In addition, numerous bone and stone artifacts were found such as spoons, hooks, awls, axes, adzes, hammers, etc. There are also numerous finds of ceramic altars, ritual vessels, and zoomorphic and anthropomorphic figurines. The most notable find is a female figurine with pronounced steatopygia, better known as the "Red-Hair Goddess" [15–18].

- **I7712**: grave 2, DON4, 6000-5300 BCE

In probe I/66, pit 1 of irregular shape (layer 1), an incomplete skeleton of an adult individual was found. The bones were in dislocated position, scattered throughout the pit. There were no grave goods [16,17].

##### 4.3 Siklós-elkerülő út (Csukma-dűlő) (Hungary)

**Author**: János Jakucs

The site 'Csukma-dűlő' is located in the north-western part of the town of Siklós, near the locality of Máriagyűd, in Baranya County (Southern Hungary). The site was discovered during the construction of the road bypassing Siklós from the north, in 1999 (archaeologist: István Ecsedy). Remains of a large-scale settlement of the early Neolithic Starčevo culture wereeen observed during the excavation. A vast number of ceramic fragments, burnt daub, animal bones, and stone tools were uncovered from the early Neolithic layers and pit-complexes. One of the partially uncovered early Neolithic pit-complexes contained the poorly preserved skeleton of a 35-36-year-old female. On the basis of the pottery style, the discovered assemblage is plausibly derived from the latest phase of the Starčevo culture.

- **I29876**: HUNG 447/HUNG 483, 5700-5500 BCE

##### 4.4 Vörs-Máriaasszony sziget (Hungary)

**Authors**: Nándor Kalicz, Katalin T. Biró, Zsuzsanna M. Virág

On the territory of the village Vörs, lying at the eastern margin of the former Kis-Balaton („Little Balaton") marshes, on the western border of Somogy county facing the neighboring Zala county, were sites and finds from various periods from almost all periods of prehistory since the Early Neolithic. Finds include materials from the Early Neolithic (Starčevo culture), from the Early Copper Age Lengyel III. culture, from the Middle Copper Age Balaton-Lasinja culture, from the Late Copper Age Kostolac culture, from the Early Bronze

Age Kisapostag culture, from the Late Celtic and (Early-) Roman period, and from the (Early Medieval-) Árpád-dynasty period.

- **I17927**:HUNG517, Grave 2, 5800-5300 BCE

#### 4.5 Vinkovci – NaMa (Croatia)

**Authors:** Mario Šlaus, Željka Bedić, Maja Krznarić Škrivanko

This Starčevo site was discovered during a rescue excavation in Vinkovci, the peripheral part of tell Tržnica. According to archaeological findings, the site is chronologically defined as the late-classical Spiraloid B phase of the Starčevo culture. Four burials associated with Starčevo culture contained skeletons in flexed positions lying either on their left or right side. Only one burial contained grave goods in a form of a ceramic vessel.[19]

- **I28426**, grave 7, 6000-5300 BCE

  The skeleton belongs to a male aged 17 to 19 years at the time of death. Pathological changes include cribra orbitalia in the left orbit, ectocranial porosity on both parietal and the occipital bone, and benign cortical defect on the latissimus dorsi muscle attachment of the right humerus.

- **I28425,** grave 11, 6000-5300 BCE

  This morphological female aged 30 to 35 years exhibits only dento-alveolar pathologies: five caries lesions, two abscesses, and six teeth lost antemortem.

- **I28427**, grave 15, 6000-5300 BCE

  This morphological male skeleton aged 30 to 35 years has several pathological changes. Ectocranial porosity is present on both parietal and the occipital bone. Linear enamel hypoplasia is recorded on the maxillary and mandibular canines. Degenerative osteoarthritis is present on one thoracic and one lumbar vertebra and

Schmorl's node on one thoracic vertebra. On the lateral condyle of the distal right femur, a possible osteochondritis dissecans measuring 14×10 mm is present while on the same location on the left femur a protrusion measuring 8×5 mm is observed.[20–22]

##### 4.6 Magura-Buduiasca (Romania)

**Authors:** Catalin Lazar, Pavel Mirea

The site of Magura-Buduiasca (TELEOR 003) is located on the lower eastern terrace of Teleorman River, 8 km from Alexandria town and 45 km north of the Danube River. This flat settlement includes several habitation horizons belonging to the Early and Middle Neolithic period (Starčevo-Criş, Dudeşti, and Vădastra cultures), spanning 6100 to 5200 BCE.

The Early Neolithic site of Magura occurs in an area of loess soils that overlie marl (calcareous sediment that includes silt-sized quartz) on the edge of high ground overlooking the floodplain of the Teleorman River. The habitation consists of pits, hearts, and pit huts with different dimensions and depths. All these features are cut into the marl [23]. The features contain rich material culture: potsherds, figurines, flint and stone tools, grinding stones, wood items, bone ornaments and tools, shells, and animal bones [24]. One of the significant features of this settlement is the presence of a considerable number of scattered human bones [25].

Pit C57 (aka Cpl. 57) was identified in the S48 survey. It had an irregular shape , with more fill levels that contained pottery sherds, flints, other small finds, animal bones, and human bones. All human bones are scattered in the pit fill, without anatomical connection or other sign of funerary treatment. Generally, the anatomical elements represented here are fragments of skulls, fibula and femur diaphysis, vertebrae, phalanx, and teeth. According to the C14 data, the C57 pit was used between 6066-5741 cal BCE. From the C57 feature, two samples were analyzed in this study.

The analyzed samples included in this study come from a pit (C57) and the Starčevo-Criş habitation levels (Supplementary Figure 15).

- **I6174** ROM-02-2017, 6074-5927 calBCE (7155±35 BP, PSUAMS-3903)
  A left mandibular central incisor (LI1) was identified in Unit 2870, at -1.40/-1.50 m. It belongs to an adult.

- **I6699** ROM-05-2017, 5292-5000 calBCE (6180±45 BP, PSUAMS-3983)
  A right mandibular central incisor (RI1) was identified in Unit 2871, at -1.50/-1.60 m. It belongs to an adult. This incisor along with the other one identified in Unit 2870 probably belongs to the same person.

- **I6181** ROM-07-B1, 6000-5300 BCE
  An isolated human mandible fragment was identified in Unit 2874, at -1.70/-1.80 m. It belongs to an adult inferred to be morphologically female.

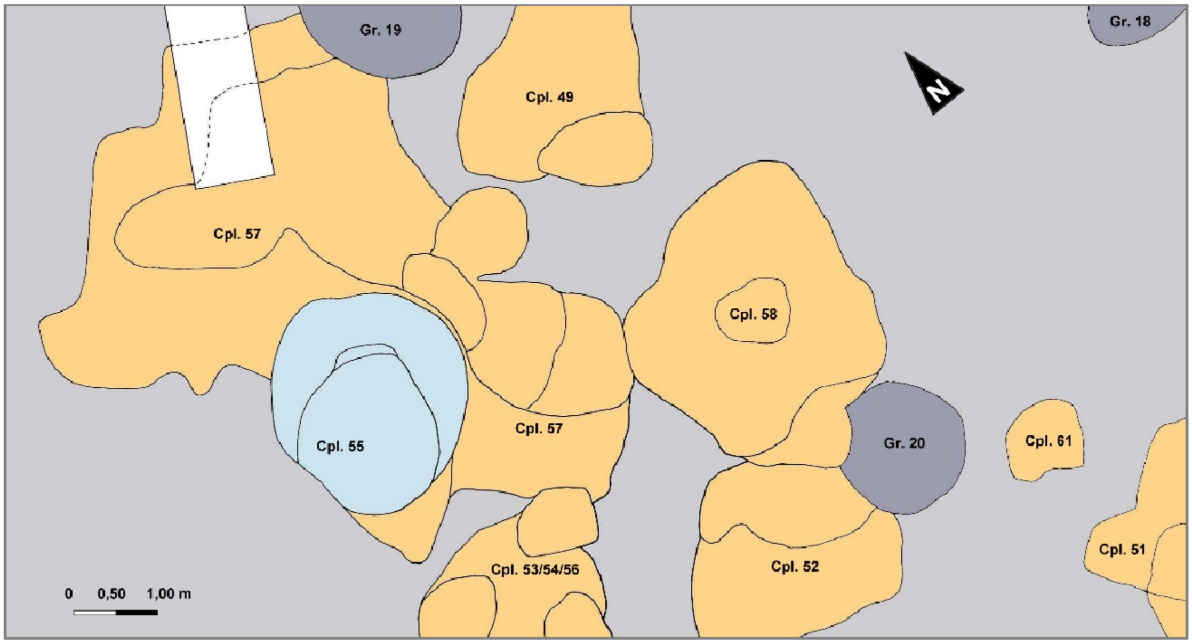

**Supplementary Figure 15:** Map of the S48 survey and location of pit C57 (Cpl.55) and Cpl.57. Image from Catalin Lazar with rights of publication.

##### 4.7 Panečevo, Starčevo-Grad (Serbia)

**Author:** Andrej Starovic, Marija Djuric

- **I8116**: PANC_2 , 6000-5300 BCE
- **I8109:** PANC_11, 6000-5300 BCE
- **I8104**: PANC_4, 6000-5300 BCE
- **I8105**: PANC_5, 6000-5300 BCE
- **I8117**: PANC_7, 6000-5300 BCE

##### 4.8 Donja Strana-Velesnica (Serbia)

**Author:** Dusan Boric

Velesnica is situated on the right bank of the Danube in the Ključ region of the Danube Gorges area, around 50 km from Lepenski Vir as the crow flies or around 100 km downstream along the Danube. The area of 660 $m^2$ was excavated in 1980–1982: [26]. There is a very thin, lowermost, occupation layer that can be assigned to the Mesolithic, with several animal bones and bone tools [27] found beneath the Early Neolithic levels. This layer remains undated at present and was found beneath the layer with Early/Middle Neolithic Starčevo ceramics, burnt house daub, and other artifacts. The Early Neolithic phase at Velesnica is characterized by Starčevo culture ceramics and the thickness of the occupation layer varies from 1.5–1 m along 80 m of the river bank. Irregular stone constructions and a circular hearth with burnt soil, as well as a stone foundation, were found.

The burials at Velesnica can be dated to the Early/ Middle Neolithic period on the basis of their stratigraphic context, association with Neolithic material culture, and crouched body position of the deceased [26,28]. They have also been directly AMS-dated [29]. The burials were concentrated in two adjacent excavation areas marked as Block A and Trench 8 [28]). Burial 1 was a very contracted child skeleton aged between three and seven years old, which lay on its lateral left side and was oriented north-south. The burial was found directly underneath a large stone "altar"/mortar [26]), the pressure from which appears to have crushed the child's skull, which was very fragmentary [30]). A ceramic bowl accompanied the individual in this burial and can probably be considered a grave offering. Burial 2A-G at Velesnica contained the remains of five primary burials, some of which were placed one on top of the other [26]). There were also the remains of two disarticulated individuals. This multiple-burial was located in Trench 7, around 15 m north of Trench 8, and the inhumations were found within a pit with a diameter of around 1.2 m infilled by Early Neolithic Starčevo ceramics, red burnt soil, mollusk shells, and animal bones. The difference in depth between the bottommost burial (at 34.84 masl) and the topmost burial (35.30 masl) was only 0.46 m, and the excavator notes that the burial cut was made from only 0.1 m above the head of the most recent burial, at around 35.4 masl [26]). The most recent inhumation, Burial 2A(+2E), was of an adult woman in a very crouched position, perhaps on her right hip, oriented east-northeast and located along the north edge of the burial pit. A second adult female, Burial 2B(+2F) (Supplementary Figure 16), was placed in the center of the burial pit and oriented east-west. She lay on her lateral left side and her upper limbs were flexed at the elbow. This individual was also characterized by a series of dental pathologies, including caries on two teeth, which is infrequent in the preceding forager population of the Danube Gorges [30]. On the southern edge of the pit lay Burial 2C, of a child of between seven and eleven years of age. The child was laid to rest on its lateral right side with the upper limbs flexed at the elbow, and was oriented west-east. Burials 2B and 2C were found at approximately the same level within the burial pit, faced each other, and were arranged in symmetrical positions. In the central part of the burial pit and directly beneath Burial 2B, Burial 2D was found, of an older adult female. This individual was laid to rest on her lateral left side and was oriented northeast-southwest. Thus, superposed Burials 2B and 2D were both laid to rest on their lateral left sides but with diametrically opposite orientations, so that the skull of Burial 2B lay over the lower limbs of Burial 2C [28 31]. At the bottom of the burial pit was Burial 2G, of a child lying with the torso on its back, with the thigh bones flexed on the torso and with the legs flexed on the thigh bones, splayed outwards and crossed at the ankles. The final burial from Velesnica, Burial 3, was found in block A, and was of a young woman laid to rest in a crouched position on her right side and oriented south-north [32]). [29] reported seven AMS measurements that date seven individuals from Velesnica. All available skeletal remains from the site have now been dated apart from burial 3. The obtained date for Burial 1, after correction for the aquatic reservoir effect, falls in the first 150 years of the sixth millennium BCE (95% confidence). On the other hand, measurements obtained for six primary burials in multiple Burial 2, after correction for the aquatic reservoir effect and modeling within the Bayesian statistical framework, suggest that all these burials were placed here most likely between 6065–5995 cal BCE (95 % probability) [31]. The exception is a Middle Mesolithic date obtained on disarticulated remains of a neonate found with primary Burial 2G at the bottom of this multiple burial place.

- **I5770**: VELE_2B, 6215–6020 cal BCE (7385±39 BP,OxA-19192 )

Velesnica Burial 2B (Lower left PM4, deformed and worn). Context: Tr. 7/7, spit XVIII-XIX. The remains of this individual were directly AMS-dated by OxA-19192: 7385±39 BP (uncorrected for the freshwater reservoir effect); 7235 ± 44 BP (corrected for the freshwater reservoir effect); 6215–6020 cal BCE (calibrated range at 95% confidence); δ13C=–19.3 δ15N =10.7, C/N=3.2

- **I13162**:VELE_2C, 6081–5912 cal BCE (7145 ± 45, OxA-19209)

Velesnica Burial 2C (right petrous). The remains of this individual were directly AMS-dated by OxA-19209: 7245 ± 39 BP (uncorrected for the freshwater reservoir effect); 7145 ± 45 BP (corrected for the freshwater reservoir effect); 6081–5912 cal BCE (calibrated range at 95% confidence); δ13C=–19.2 δ15N =9.9, C/N=3.2

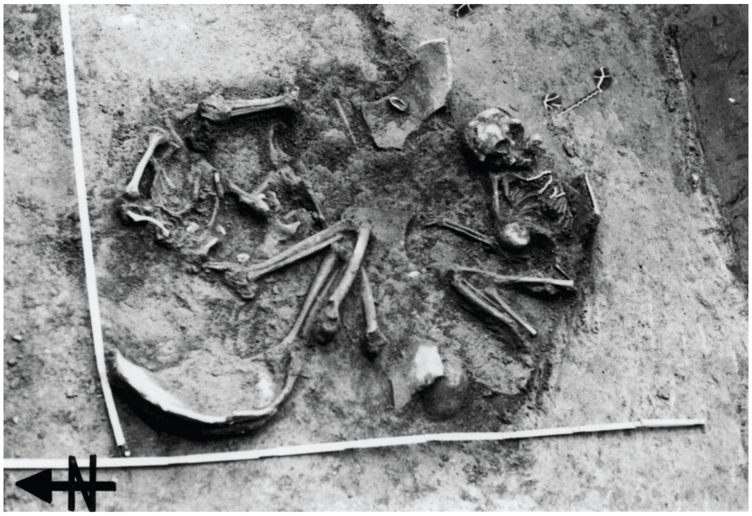

**Supplementary Figure 16:** Individuals 2B(+2F) and 2C, Velesnica, found buried in flexed positions on their lateral sides symmetrically facing one another. Picture from Dusan Boric with rights of publication.

##### 4.9 Mostonga IV Mostanica (Serbia)

**Author:** Tamas Hajdu

A single grave dating back to the 6000-5300 BCE consisting of a 13-15 year old individual. [33].

- **I7869**: MSTG_1, 6000-5300 BCE

##### 4.10 Hencida, Csörsz-árok II./Gyűrű-szeg 2. (Hungary)

**Author:** László D. Szabó

- **I17455**: HUNG515, Feature 9, 5800-5300 BCE

  The individual burial of a juvenile (14-15 years old) (Supplementary Figure 17) was found in the sub-humus layer in the probe nr.111. The deceased boy was lying in the fetal position (strongly contracted legs, drawn up tightly; arms bent at the elbows, pulled in front of the face) on his left side. No grave goods were observed. The body was positioned in the E-W direction. The grave pit was rounded, rectangular, and shallow.

- **I18642**: HUNG516, feature 5 , 5800-5300 BCE

  This is a child burial (9-11 years old) found in the fill of a shallow Early Neolithic (Koros) waste pit (Feature 5/Str 5). The deceased was lying in the fetal position (strongly contracted legs, drawn up tightly; arms bent at the elbows, pulled in front of the face) on the left side. The body (especially the part behind the skull) was surrounded by pottery and bone fragments. A bovine mandible was found 30 cm behind the body. These are interpreted to be finds from the pit and not intentional grave goods.

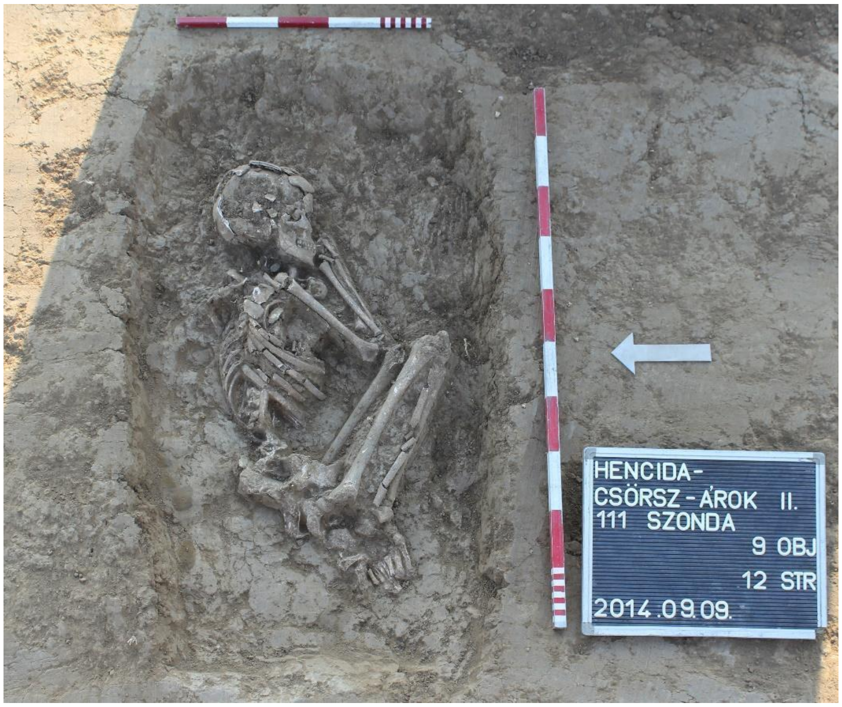

**Supplementary Figure 17:** Dating: the burial can be dated to the Early Neolithic Körös culture. Image from László D. Szabó with permission of publication.

#### 4.11 Dévaványa–Kér-sziget, Katonaföldek (Hungary)

**Author:** István Ecsedy

The site is located along the north-western bank of the river that surrounds the island of Kér, on the outskirts of Dévaványa in SE-Hungary. A topographic survey of the site was carried out in the late 1960's, which documented the remnants of rectangular houses about 20-30 m apart and nearby rubbish pits. The archeological team was also able to locate the Early Neolithic part of the site on the strip of the former riverbank.

- **I17931:** HUNG 667, grave 1: 5800-5300 BCE
  Burial of an adult female lying on her right side, in contracted position. The NE/SW oriented skeleton was complete, without grave goods. The burial feature was rectangular shape, in a grave pit with a rounded corner.

#### 4.12 Berettyóújfalu and Szentpéterszeg–Körtvélyes II (Hungary)

**Authors:** Tamara Hága

The site is located between Berettyóújfalu and Szentpéterszeg, north of the Berettyó River, east of the Herpály tell, on the southern high bank of a watercourse. After a rescue excavation, several features from different periods were found, the vast majority of which dated to the Neolithic. In the south-eastern part of the site there were Early Neolithic (Körös culture) and Middle Neolithic (Alföld Linear Pottery culture, Esztár group) settlement features and burials, while in the north-western part of the site, on a hill, Late Neolithic (Herpály culture) features were excavated.

The burials and human remains related to the Körös culture were not found in separate burial graves, but in settlement features. One human skeleton was usually found in each feature. The exception was the largest pit (Feature 50, SNR: 81) dating to the Early Neolithic, which was originally used for clay mining and which contained sixteen contracted burials and the remains of eight human skulls. The amorphous pit, measuring 21 × 17 cm, was 180 cm deep from the surface without humus. It contained very few finds (pottery, a steatopygous idol, animal bone, wattle and daub, snail, shell, chipped and polished stones, grinding stone) compared to its size and to other Early Neolithic features. All but one of the skeletons and skulls were located at or near the bottom of the pit, on a thin layer of fill. The filling of the graves was identical to that of the pit, with no evidence of digging in. Only the upper part of the pit was intersected by a ditch system from the Árpád Period. An intact horn of an auroch was also found near the burials in the northern half of the pit, on the same level as the burials. The majority of the human remains were female: nine of the sixteen skeletons were female, three were male, and four of them were children. Five of the eight skulls belonged to women, one to a man, one to an adult of indeterminate sex, and one to a child. The orientation of the graves and the degree of their contraction varied [34].

- **I14962**:Obj. 88, Strat. 144, 5665-5525 calBCE (6680±35 BP, PSUAMS-9998)

- **I14965**:Obj. 82, Strat. 132, 5800-5300 BCE

- **I15073**:Obj. 141, Strat. 2595835-5484 calCE [union of two inconsistent dates: 5835-5667 calBCE (6865±35 BP, PSUAMS-10195), 5655-5484 calBCE (6665±35 BP, PSUAMS-9999)]

- **I15074**:Obj. 320, Strat. 623,5800-5300 BCE

- **I18614**:Obj. 309, Strat. 606, 6000-5300 BCE

##### 4.13 Turia (hu: Torja) – Apor-kúria kertje (Romania)

**Authors:** Sándor József Sztáncsuj

The site of Turia–Apor-kúria kertje is located in South-East Transylvania, in the piedmont area on the western edge of the Black River (Feketeügy/Râul Negru) basin. The site was discovered and researched by Zoltán Székely in 1984–1986. Archaeological excavations have revealed traces of habitation from several periods, from the Neolithic to the Middle Ages. The earliest settlement remains from the site belong to the Neolithic Starčevo-Criș Culture, consisting of a 40-50 cm thick habitation layer. Excavations brought to light the remains of three semi-subterranean houses, each of them with an area of approximately 3×3 m, with walls built on a wooden structure covered with clay. The rich archaeological material discovered in the habitation layer and inside the houses, consisting of pottery, stone and bone tools, and various clay artifacts, date the settlement to the late phase of the Starčevo-Criș Culture. Four inhumation graves have also been unearthed from the area of the early Neolithic settlement. The individuals were lying in a contracted position, oriented north to south. The particularly poor funerary inventory consisted of a few fragments of vessels deposited next to the deceased. One of the graves, with no funerary inventory, was disturbed by a pit from a later period.

- **I21906**: Grave 2 , 6000-5300 BCE

Burial of a (possible) adult, discovered in 1984, in trench number II, at a depth of 37 cm. The body was lying on the right side, in a contracted position, oriented north to south, with the head on the north side. The skeleton was incomplete, with only the upper part being preserved, the rest being destroyed by a pit from the Dacian period. No burial goods.

##### 4.14 Arnót –Arnóti-oldal Dél (Hungary)

**Author:** Krisztián Tóth

The Arnóti-oldal Dél site is situated in Borsod-Abaúj-Zemplén county, Northeastern Hungary, 2 kilometers from the eastern part of Miskolc city and 900 meters to the west of the village of Arnót. The Sajó river widens and branches here . According to maps of the military surveys of the 18th and 19th century, the area between the river branches was marshland. Geographically, this is the border between the Great Hungarian Lowland and the North Hungarian Mountains. In this swampy territory, small hills were suitable for human activity. This includes the hill where the site is located, close to the eastern river branch of the river, which is called Kis-Sajó (Small Sajó river).

A total of 5 Neolithic burials were also found at the site, including individual S3 buried 23 meters to the North from the well. The grave was separated by a grayish-brown, slightly oval-shaped patch from the light brown loam, with an obsidian splint on its surface. After digging it, there was a bench 34 cm deep in a circle near the 42 cm deep straight bottom of the tomb. The grave pit was 163 cm long and 130 cm wide. The 35-45 years old (Supplementary Figure 18) male was in a sleeping position orientated to the southeast facing to the south. North of the skeleton and just behind it was a conical, straight-walled bowl (2) On it lay a fragment of a smaller vessel (1). At the pulled-up knees south of the body there was an 8 cm long obsidian blade (3) [35]**.**

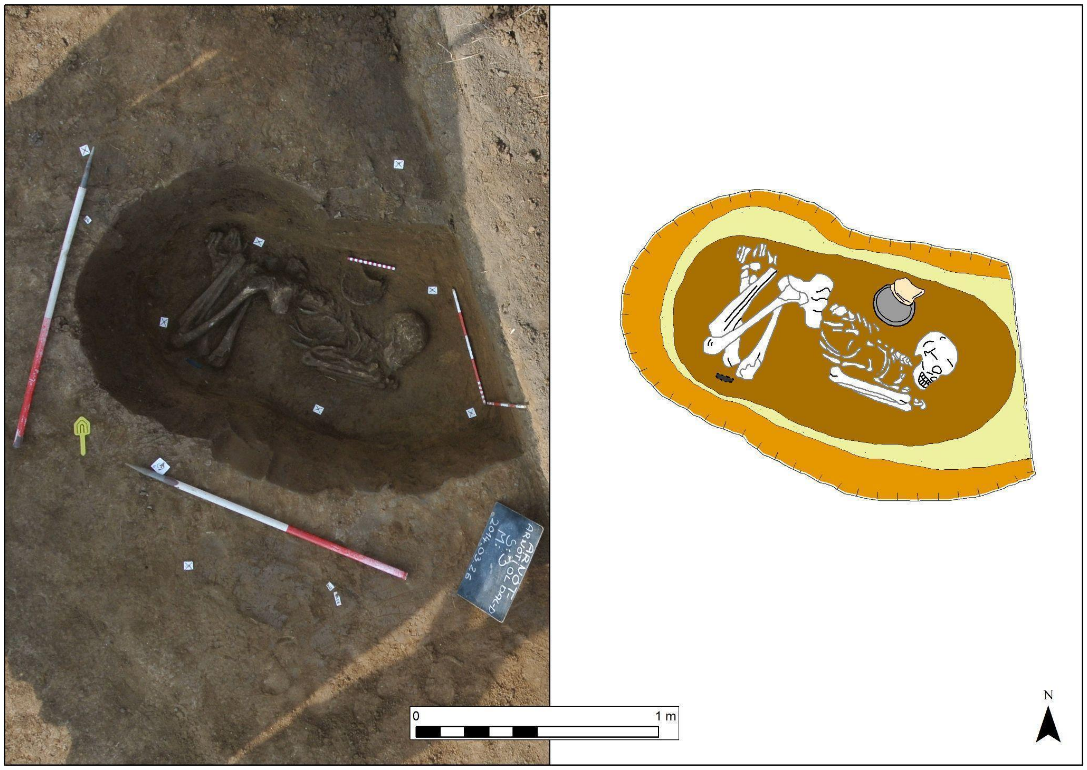

**Supplementary Figure 18:** Individual from Arnót – Arnóti-oldal Dél. Figure from Krisztián Tóth with rights of publication.

- **I29958**: HUNG 954, grave s3, 5500-4500 BCE

- **I29957**: HUNG 953, grave s18, 5500-4500 BCE

##### 4.15 Tiszaszőlős-Domaháza (Hungary)

**Author:** László Domboróczki

The site is located 40 km to the north of Szolnok, along the Tisza river. The site was first occupied by Körös people (one pit and house identified) and later settled during the ALPC period[36].

Remains of 7 individuals have been reported: two complete skeletons, two partial skeletons identifiable by their dispersed body parts, and three other individuals represented only by single bones. The most complete skeletons are the ones found in graves 1 and 6.

- **I21828**: grave 1, HUNG344, 5080-4780 calBCE (6040±60, deb10901)

  40-46 year-old male in contracted position on his left side. The radiocarbon dating shows that it belongs to the late ALPC period.

- **I21830**: grave 6, HUNG346, 5220-4780 calBCE (6060±80, deb-11084)

  34-40-year-old male in contracted position on his left side and dated to the late ALPC period.

##### 3.16 Füzesabony-Gubakút

**Author:** László Domboróczki

The Füzesabony-Gubakút site is situated at the northernmost edge of the Great Hungarian Plain, approximately 3 km south-west of the modern village of Füzesabony and about 10 km to the south of the Mátra-Bükk foothills. The area of the ALPC settlement is separated into two parts by a broad but shallow valley that was cut by a one-time riverbed into late Pleistocene sediment. The site was rescue-excavated between 1995 and 1996 before motorway construction.

The most significant result of this excavation was the discovery of an idiosyncratic settlement layout that has been documented since that time at other ALPC sites [37]. The evidence of triple-partitioned houses, 12–16 m long by 5–6 m wide, and the regular settlement structure here contradicted earlier views of small, irregular ALPC settlements made up principally of pit dwellings. In Füzesabony-Gubakút, the houses were arranged in rows on either side of a stream. Between the houses there were pits, and at the corners of the houses, there were human burials. These graves were dispersed throughout the settlement, all left-crouched and orientated with the head to the southeast or east. Seven graves were furnished with grave goods that were, except for one pot, entirely Spondylus ornamentation. Four graves were unfurnished.

The topography of Füzesabony-Gubakút and the many radiocarbon dates obtained have presented an excellent opportunity to examine the chronology of the site and its internal development. Informal inspection of the radiocarbon results (based on the animal bone from the pits) has suggested 12 settlement phases, each lasting 30–50 years, between 5560–5000 cal BCE [38]. This was supported by seriation based on 74,580 sherds derived from 18 of the 28 Neolithic pits found at the site. House-pit-grave ensembles could be identified as household areas and assigned absolute time limits. Using surface collection data gained in 2007, the whole settlement area could be modeled. The working hypothesis was that the settlement began with the appearance of a pioneer family at the site around 5560 cal BCE and through a continuous history as well as dynamic growth reached its peak with 12–14 contemporaneous houses/families between 5325–5220 cal BCE [38].

Around Füzesabony, an interesting network of ALPC settlements has been recorded, which consisted of regularly spaced larger settlements similar to Gubakút, surrounded by smaller settlements in their close vicinity, echoing the model of larger, central/mother settlements, with smaller, satellite/daughter sites around them proposed elsewhere for the ALPC distribution [38]. The large settlements were aligned along ancient riverbeds and located at regular intervals of some 2 km from one another. By calculating house numbers and possible population levels, and estimating the size of herds and the scale of meat consumption, it was concluded that circles of 1 km diameter probably met the requirements for the pasturage necessary for grazing 15 cattle and 30 sheep per household in a settlement of 14 families. This large land requirement suggests that the extent of the pasturage would have been the greatest determining factor in the sustainability of a settlement during this phase of the Neolithic. Building on this model it was suggested that inter-settlement relationships may have also influenced the demographic development of the region and its settlement pattern, and the individual central settlements — such as Füzesabony-Gubakút — represented different lineages or clans. Modeling the process of settlement development at different scales (by analyzing different patterns and using terms such as local development, in and outward migrations, demographic pressure, etc.) it was concluded that Neolithization may have proceeded along similar principles for hundreds of years in the region [37,38].

- **I10351**: grave 5, 5370–5200 calBCE (6295±40, VERA-4242)
- **I10352**: grave 3, 5220-4780 calBCE (6060±80, deb-11084)
- **I10353**: grave 2, 5466–4991 calBCE (6250±90, deb-11092)
- **I10355**: grave 1, 5350–5210 calBCE (6295±35, VERA-4237)
- **I10349**: grave 10, 5292-5046 calBCE (6200±30 BP, PSUAMS-10194)

- **I10350**: grave 4, 5380–5210 cal BCE (6325±35, VERA-4236)

#### 4.17 Polgár-Ferenci-hát (Hungary)

**Author:** Alexandra Anders

The ALPC site of Polgar-Férenci-hát is located east of the Tisza river, to the north of the great Hungarian Plain, and was first studied in 2004 in a rescue excavation. The site shows two different stages of occupation. The first phase is ALPC I (Szatmár), with grave 718 being a remarkable specimen with a particular grave differentiated from the res t[4]. The second occupation corresponds to phases II-IV of the ALPC. A total of 113 graves have been excavated, with radiocarbon dates spanning from 5300 to 5070 cal BCE. The graves are distributed in small clusters. Previous dietary Carbon/Nitrogen isotope analyses [5] revealed no differences between males and females, and no correlation to grave goods.

The Neolithic settlement of Polgár-Ferenci-hát lines in the Polgár Island micro-region within the Upper Tisza region. This site offers a unique possibility for tracing the cultural changes in the later sixth millennium BCE, as well as the local dynamics of the internal transformation of the Alföld Linear Pottery (ALPC) of the Middle Neolithic.

As previously mentioned, there are two occupational phases in Polgár-Ferenci-hát: The first can be dated to the earliest phase of the ALPC (The ALPC-I: 5467 and 5344 cal BCE). During this period, the settlement had a rather dispersed layout, with six burials and some smaller and larger pits lying quite far from each other. The second period (ALPC II-IV: 5293–5068 cal BCE), this phase corresponds with a period of more intensive usage of the site.

The accumulation of several layers in this central part of the settlement is indicative of the type of stratigraphic events that led to the emergence of tell sites in the southern section of the Great Hungarian Plain, near Szakálhát and Esztár type settlements. Outside the enclosure, traces of a dispersed horizontal settlement were found. With an overall horizontal extent of 9-12 ha and a vertical accumulation of strata in its center, the settlement of Polgár-Ferenci-hát represents a dualistic site-formation process.

Characteristic sherd types found within the closed artifactual assemblages related to the circular ditch system recovered at Polgár-Ferenci-hát are indicative of a synthesis between the Tiszadob-Bükk and Esztár-Szakálhát-Vinča ceramic styles. Previously, late stylistic groups of Alföld LBK pottery were often classified into different chronological phases. Nevertheless, distinctly synchronous occurrences recorded at this site raise the need for seriously reconsidering the relative chronology of these styles. The ALPC II–IV occupation at Ferenci-hát can be dated between 5293–5068 cal BC.

In addition to settlement features, 116 graves were excavated. These graves represent the largest number of ALPC burials in the region of the Tisza River. The most outstanding burial of Phase I was Feature 718. A woman of 22–28 years of age was laid to rest here on her left side, in a crouched position. She wore a string of Spondylus shell beads around her neck. She was buried with 6 regular and 5 miniature vessels and an anthropo-zoomorphic figurine.

The graves of ALPC II–IV Phase were clustered in groups of 3–10 smaller and greater concentrations of burials within the excavated area. The grave pits were oval or trapezoid. Sometimes the deceased was not placed in an ordinary grave pit but in a storage pit. Thanks to the water-logged deeper strata, traces of a coffin or a wooden bier could also be observed for the first time in the history of the Middle Neolithic in Hungary.

The most common orientation was SE to NW, with only minor deviations. With only three exceptions, the deceased was laid on their left sides in a more-or-less crouched position. The majority of the deceased were placed in the grave alone. However, two double burials were observed as well. In four cases, the deceased were buried without their heads. Two of these people were buried in proper graves, while the other two were placed in storage pits. All four of them were men of ature adult ages. According to studies by Zsuzsanna Zoffmann (physical anthropologist), no cut marks were present on the remaining bones, that is, the deceased was not decapitated but their skulls were removed sometime after burial. Two scattered-ash burials are of special significance. Based on previous research, evidence of cremation is known only from the Late Neolithic in Hungary.

In almost 10% of the graves, traces of ochre were found under the skull and the bones of the leg. Eighty-two of the 113 excavated graves contained no grave goods. A total of 16 burials contained vessels. They usually occurred as single finds, although some of the deceased were buried with 2 or even 3 vessels. A special group of burials with vessels is represented by three graves in which vessels full of ochre were placed into the grave pit. Another group of important grave goods is represented by stone tools, although only six graves contained obsidian blades or ground stone chisels which occurred in the burials of women, men, and children alike. Two additional burials deserve special attention. Both of them contained a large obsidian core (exceeding 10 cm in length) placed near the head of the deceased. There were also signs of secondary, post-burial placement of grave goods (Nachgaben). In these cases, pottery sherds were discovered in the end of the grave pit, but well above the skeleton. A total of 24 neonate and child burials were excavated at the site. Pots and jewelry had been placed in the graves of 12 children. The differential distribution of Spondylus objects is perhaps the most pronounced difference between adult and child burials. A total of 38 such ornaments came to light from seven burials and comprised large beads, necklaces and bracelets strung of smaller beads, and arm rings. Of these, only one bead was recovered from the burial of an adult male; the other six graves contained the burials of newborn infants and children aged less than 6-7 years.

Overall, the burial rite observed at the site of Polgár–Ferenci-hát was similar to the general mortuary behavior of the ALPC groups, and largely similar care was taken of the deceased within the broader distribution area of the LBK culture.

- **I21826**: HUNG471, grave 821, 5500-4500 BCE
- **I21827**: HUNG469, Grave 890, 5500-4500 BCE

- **I21840**: HUNG450, grave 342, 5500-4500 BCE
- **I21842**: HUNG457, Grave 134, 5500-4500 BCE
- **I21843**: HUNG448, grave 126, 5500-4500 BCE
- **I21844**: HUNG451, grave 4, 5500-4500 BCE
- **I21847**: HUNG520, grave 341, 5500-4500 BCE, (6165±35, VERA-4333)
- **I17909**: HUNG1109, grave 786,  5500-5100 BCE
- **I17911**: HUNG1026, grave 719, 5500-5100 BCE
- **I18656**: HUNG623, grave 691, 5500-4500 BCE
- **I18658**: HUNG622, Grave 353, 5500-4500 BCE
- **I18659**: 3688, Grave 352, 5500-4500 BCE
- **I18661**: HUNG624, Grave 768, 5500-4500 BCE (6185±40, VERA-4338)
- **I18657**: HUNG631, Grave 773, 5500-4500 BCE, VERA 3058, 6250,35
- **I17950:** 517, grave 338, 5500-5100 BCE
- **I21898:** 1028, grave 721, 5467-5219 calBCE (6355±30 BP, PSUAMS-10203)
- **I18660:** HUNG633, grave 718, 5500-4500 BCE, VERA-3055: 6260,40
- **I18695:** HUNG642, grave 904, 5500-4500 BCE

- **I21902:** PF644, grave 644, 5371-5216 calBCE (6330±30 BP, PSUAMS-10204)
- **I21899:** PF448, grave 448, 5500-5100 BCE
- **I21901:** PF94, grave 34, 5500-5100 BCE
- **I21763**: 83, grave 34, 5500-5100 BCE
- **I21765**: 252, grave 144,  5500-5100 BCE
- **I21766:**  PF283, grave 283, 5500-5100 BCE
- **I21767:** PF283,  grave 288,5500-5100 BCE
- **I21768:** PF348,  grave 348,5500-5100 BCE
- **I21769:** PF78, grave 783, 35500-5100 BCE
- **I21771:** PF822, grave 822, 5500-5100 BCE
- **I21846:** PF900, grave 900, 5500-5100 BCE
- **I21714:**  PF31, grave 31,5500-5100 BCE
- **I23093:** 468, grave 296, 5500-5100 BCE
- **I23094:** 91,  grave 34, 5500-5100 BCE
- **I21897**: 1105,  grave 782, 5500-5100 BCE

- **I21770:** PF801, 5045-4842 calBCE (6050±35 BP, PSUAMS-10207)
- **I21841**: HUNG458, 5500-4500 calBCE, VERA-4331: 6260,40
- **I17910**: 1128, grave 815, 5500-5100 BCE
- **I18662**: 353, grave 356, 5500-4500 BCE
- **I21824**: HUNG467,  grave 807, 5500-4500 BCE
- **I21825**: HUNG470,  grave 867, 5500-4500 calBCE (6235±35 BP, , VERA-3056)
- **I21772**: 123, grave 871, 5500-4500 BCE
- **I21773**: 1236, grave 881 5500-4500 BCE

##### 4.18 Rákóczifalva–Bagi-földek sites 5 and 8 (Hungary)

**Author**: Katalin Sebők.

This chain of sites is situated on a somewhat bluff-like, flood-free bank of a one-time bed of the river Tisza southwest of Rákóczifalva in Central Hungary. Today, the river runs to the west of the sites along a new bed laid out in the 19$^{th}$ century. As part of creating a reservoir, extensive preventive excavations were carried out along the line of a new dam, cutting through the top of the one-time flood-free bank and, thus, the highest points in the area, between 2005 and 2007. Not surprisingly, the plateaus and high slopes of the bank's sandhill row were used in multiple historical periods, including the Middle and Late Neolithic, the Early Copper Age, the Late Bronze Age, as well as the Early Imperial (Sarmatians), Migration (Gepids and Avars), Hungarian Conquest and Late Mediaeval Periods and the Modern Age.

The sampled sites line up next to each other. Site 8 lies at the westernmost end of the chain of hills next to the current riverbed, and probably in the immediate vicinity of the river at the time of the Neolithic and Copper Age occupations. To the east of Site 7, separated by only a slight depression, lies site 8A, covering the highest point of the area. The eastern slope of site 8A concludes in a rather extended but less densely occupied site, no. 5, stretched on a less prominent but still high plateau consisting of several lesser elevations.

The sameness of the occupations on the three sites is manifest in the fact that the cultural features associated to each use period do not necessarily cease at the geographic border but continue on into the "next" sites. Thus, Neolithic and Szakálhát features scatter on all three sites, while the traces of settlements in the Early Copper Age were found in sites 8 and 8a.

The cultural features determined as Neolithic and Szakálhát probably belong to the same settlement. This record has not been evaluated and processed yet; therefore, the following numbers cannot be regarded as final. On the three sites (5/8A/8), altogether, 225 features (73/104/48), comprising about 30 (3/22/5) inhumation burials, belonged to this era. There are no available radiocarbon data for these features.

All burials seemed to follow the same funerary rite: the dead were laid in a narrow oval grave pit or inside a larger pit complex in a crouched position on either side. Persisting grave goods were scarce and included mainly knapped flint (and, occasionally, obsidian) tools and, rarely – probably for women– necklaces with a large cylindrical limestone or spondylus bead as a centerpiece. Several burials did not contain any grave goods.

- **I17941**: grave 272, 5500-4500 BCE

- **I17933**:grave 340, 5500-4500 BCE

- **I17938**: grave 404, 5500-4500 BCE

- **I17939**: grave 351, 5500-4500 BCE

- **I17934**: grave 41, 5500-4500 BCE

- **I18635**: grave 310, 5500-4500 BCE

- **I18636**: grave 249, 5500-4500 BCE

- **I18637**: grave 136 , 5500-4500 BCE

##### 4.19 Furta–Töviskés (Hungary)

**Author:** László Szolnoki

The site located on the outskirts of Furta, in Hajdú-Bihar county, was excavated by László Szolnoki (Déri Museum) in the autumn of 2010 in the course of a rescue excavation. On the site, 25 features were excavated from the Alföld Linear Pottery culture, one of which was a burial.

- **I17362**: grave 102, SNR146, 6000-5000 BCE

  Inhumation burial of a female in a shallow burial pit, in the humus. The deceased was lying on her left side in a strongly contracted position. The arms and legs were strongly bent. Bones are in good preservation. The skeleton was oriented from E to W. On both wrists of the deceased, there was a bracelet made of beads.

##### 4.20 Berettyóújfalu-Nagy-Bócs-dűlő 2

Author: Tamás Hajdu

- **I17947:** HUNG279, 5000-4000 BCE

##### 4.21 Debrecen–Tócó-part, Erdőalja (Hungary)

**Authors:** Emese Gyöngyvér Nagy, Zsigmond Hajdú

The site is situated in eastern part of Hungary, in the Great Hungarian Plain (Hajdú-Bihar County), near Debrecen. In 2008–2009, based on a contract with the Cultural Heritage Service, the Déri Museum carried out preventive excavations on a section of Route 4 bypassing Debrecen. The NE–SW oriented site also yielded remains from 11 other periods from the Middle Neolithic (ALPC and Esztár group). During the excavation in 2008, four ALPC graves were unearthed. Three had no grave goods at all, while the fourth contained, in addition to vessels and limestone beads, traces of ochre as well. In 2009, in the area north of the railway, we excavated six ALBK inhumation burials, six postholes that probably belong to this period, a well, and 25 pits. Four of the pits could only be partly excavated due to their large size. In the southern part of the site, beyond the railway, excavation was hindered by the fact that the Middle Neolithic (Esztár group), Late Iron Age (Celtic), Sarmatian, Árpád Period and Late Medieval settlements spatially overlapped: there were many superpositions even within a single period. The Esztár group was represented by a well, two inhumation burials and 23 huge, clay extraction pits.

According to the physical anthropological study, the Neolithic skeletal material is rather heterogeneous and is in a bad state of preservation, so it is unsuitable for generalizations and group identification.

- **I17365:** HUNG305, Feature 353,5000-4000 BCE
  Burial of a male lying of his left side, in a strongly contracted position, excavated in 2008. The skeleton was incomplete and oriented southeast to northwest 146˚ with

hyper-flexed legs. The grave was disturbed by a Sarmatian circular ditch. A pottery grave good was situated beside the legs. Other grave goods include traces of ochre under the skull and four large limestone beads.

- **I17948:** HUNG305, Feature 930, 5000-4000 BCE

  Burial of an adult male lying of his left side, in a strongly contracted position, excavated in 2009. The skeleton was incomplete and oriented southeast to northwest 108˚ with hyper-flexed legs without grave goods.

##### 4.22 Ebes-Zsong-völgy (Hungary)

**Author:** János Dani

- **I17363**: HUNG274, Feature 1412: 5000-4000 BCE

Based on the anthropological examination the deceased was a 43-60 year old (mature) woman. The burial represents the local, late phase of the ALPC (Esztár group).

##### 4.23 Debrecen-Tesco (Mikepércsi street Sports field) (Hungary)

**Author**: Márta Szelekovszky

- **I17366**: HUNG289, feature 286, 5000-4000 BCE

  This burial is of a female estimated to be 40-60 years old. The bone material is in quite fragmented and incomplete condition. Her arms were probably bent at the elbows, hands in front of the face. In front of the cranium was a fragmentary red-coloured, hemispherical vessel with knobs under the rim. The body was positioned in the ESE-WNW 102° direction (Supplementary Figure 19).

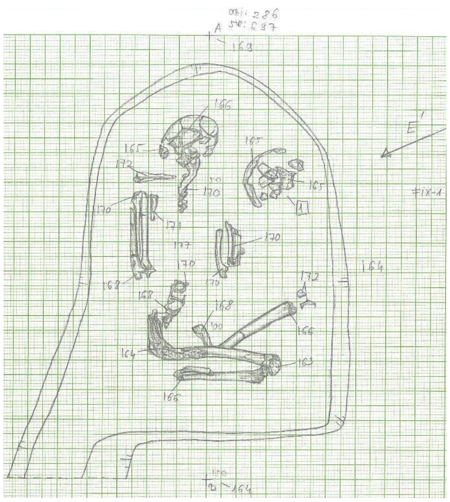

**Supplementary Figure 19:** the burial represents the local, late phase of the ALPC (Esztár group). Image from  Márta Szelekovszky with permission of publication.

#### 4.24 Vonyarcvashegy-Mandulás lakópart III. ütem site

**Author:** Tamás Hajdu

During rescue excavations, 48 settlement features (post holes, houses, pits) and one Neolithic grave were found. The settlement objects came from the Neolithic and Roman periods. In the trench of the Neolithic settlement, a child lying on his left side, without grave goods, oriented east-west (feature 71) was excavated. The width of the grave was 100 cm and its length was 155 cm.

- **I29891**: HUNG 680, 33/A, 5500-4500 BCE

#### 4.12 Apc-Berekalja (Hungary)

**Contact person:** László Domboróczki

The site is situated north of Hatvan, along road no. 21. It is primarily a late ALPC site, but the earliest phase is also present here. The uniqueness of the site lies in the fact that it sits a at the easternmost periphery and is still one of the largest settlements of the ALPC. Several houses and pits were excavated here and some graves too. [39]

- **I29877**: HUNG359, grave 12, 5367-5299 calBCE (6341±33, MAMS-14823)

  53-59 year-old male in contracted position on the left side. The degree of contraction was not as extreme as with others from this time, as the legs were not lifted up above the hip.

**4.25 Kleinhadersdorf (Austria)**

**Author:** Maria Teschler-Nicola

Kleinhadersdorf is the largest cemetery of the Early Neolithic in Austria [40], with hundreds of burials. However most of the remains excavated from it have been lost. The dead were buried either in right- or left-sided crouched position, hands in front of the face. Grave goods, single vessels, adzes, bone awls, spondylus beads, dental rolls, and in some graves red ochre scattering characterized the cemetery. The cemetery dates to the Middle Bandkeramik (Phase II), but also some graves from the older (Phase I) and younger Bandkeramik (Phase 2, cremations); i.e., the burial site had probably been used for several centuries [41,42]

- **I16006:** Kleinhadersdorf Grab 65, 5500-4500 BCE

  Excavated by Christine Neugebauer-Maresch in 199. This individual was buried in a crouched position, left side, head to the south. Preservation of the skeleton: incomplete preserved cranium, 15 mandibular teeth, postcranial remains (only diaphysis of the long bones), and bone surfaces eroded. The age at death was estimated to be 35-45 years, and the morphological sex was inferred to be female. Pathological changes included strong abrasion of the frontal teeth (probably used as a tool), strong muscle insertions at the tuberositas deltoidea (both sides) and tuberositas glutaea (these structures are often observed in Neolithic populations and associated with heavy physical load).

##### 4.26 Ratzerdorf (Austria)

**Author:** Maria Teschler-Nicola

Ratzersdorf at the Traisen Valley is a small village located near the Lower Austrian capital St. Pölten (ca. 60 km west of Vienna). From 1998 onward, agricultural areas between the northeastern outskirts of Ratzersdorf and the Kremser Schnellstrasse S 33 were converted into land for building for the development of commercial enterprises. As archaeological sites were already known and prospected by aerial photography since 1987/88 [43], preventive rescue excavations were carried out in 1998 and 1999 by the Department for Archaeological Monuments of the Federal Monuments Office. Several early Neolithic burials and other objects were identified and recovered (Neugebauer et al. 1999), and the adjacent properties were declared protected. The archaeological survey at the area (Parzelle) no. 1171 started in February 2000. Until the end of the year, about 16,000 $m^2$ were investigated and more than 700 prehistoric objects (house ground plans, storage pits, inhumation graves) of different levels were recovered.

- **I14586**: Verf. 1401, 5500-4500 BCE

The late phase of the LBK (Notenkopfkeramik) is represented by long-house floor plans, storage pits, and objects belonging to the classical and late Notenkopfkeramik. During this excavation, one inhumation burial in a crouched position was recovered within the settlement of this period [44] and included in the present study. [45–47].

##### 4.27 Egerág-Gyilkos-tó (HT 156. site), (Hungary)

**Author:** János Jakucs

This site is located northwest of the village of Egerág, in Baranya County. The site was discovered during an archaeological excavation before the construction of the Croatian-Hungarian gas interconnection pipeline, in 2010 (Archaeologist: Jácint Ligner).

The excavation took place on the plateau between the Szemely creek and the Villány-Pogány-stream. Four Neolithic pits were partially excavated, which belonged to the second half of the 6th millennium cal BCE.

Grave 2/4: This skeleton was discovered in Pit 2. The partially preserved skeleton of the 50-59-year-old man, oriented SE-NW, was lying on its right side, in a semi-crouched position: the head was strongly bent back, the torso was slightly twisted, and the legs were slightly pulled up. The burial did not contain any grave goods or findings relating to clothing. The position of the body suggests that it may not have been a traditional burial, but rather a corpse that was thrown into the pit.

- **I29877:** HUNG 484, grave 2, 5500-4500 BCE

##### 4.28 Asparn-Schletz (Austria)

**Author:** Maria Teschler-Nicola

The Early Neolithic settlement of Schletz (Supplementary Figure 20), located in northeastern Lower Austria, represents the most important Linear Pottery site in Austria. Already known for many years through surface finds, it was not until the early 1980s that greater attention was paid to it after oval-shaped soil discolourations became visible during aerial surveys by the Austrian army; these could finally be interpreted as a backfilled ditch system of a settlement [48,49]. The site was systematically investigated between 1983 and 2005 in annual archaeological excavation campaigns of the (then) Lower Austrian Provincial Museum under the direction of Helmut Windl. In the process, about 20% of the more than 254,000 $m^2$ settlement was exposed, including a complex ditch system with three ditches (to optimize the excavation area, magnetic prospecting was carried out between 1992-1995 by the Central Institute for Meteorology and Geodynamics, see aerial photograph redrawing and entry of archaeomagnetics in [50]). These ditches differ in their shape. In addition to an approximately oval construction with two sole ditches (Sohlgräben) running parallel in sections (ditch I = inner ditch; ditch II = outer ditch), a third sole ditch (ditch III) could be traced, enclosing an approximately trapezoidal area north of the oval [50–53]. They reached a maximum width of up to 4 m and a depth up to 2 m.

Currently, based on the analysis of the pottery, we assume that the settlement and all ditches date to the younger to youngest Linear Pottery, i.e. according to regional chronology to phases IIb - III [54]. According to this, the settlement is Late Linear Pottery (LBK), which was also confirmed by radiocarbon dating, which yielded an age between ca. 5210 and 4950 BCE [55,56]. Within the settlement, numerous settlement objects (postholes of several post-type buildings, cupola ovens, several storage pits, and one object that was interpreted as a well due to its internal construction and depth) and graves either near buildings or in already backfilled ditch sections could be detected. The archaeological finds spectrum includes coarse pottery (bottles, bowls, and bombs), stone and bone implements (including adzes, mace-head fragments, and awls), and abundant animal bone material primarily from the settlement pits and ditches (cattle, sheep, goats, pigs, and dogs; game was less important [50]).

The site has become the focus of much attention, primarily because of the human skeletal remains recovered from the bottom of ditch II. These remains were found in atypical poses with consistently incomplete representation (hand and foot bones and distal portions of the lower arm and leg are missing). An initial bioanthropological/forensic analysis of the skeletal individuals recovered through 1991 (N=67) revealed an unexpected age-at-death distribution that implied a deficit of young females. In addition , the bony remains were characterized by the presence of a variety of perimortem injuries, interpreted as the effect of close combat weapons (e.g., stone axes and clubs) or distance weapons (such as throwing weapons, slingshot weapons or bow weapons). It was thus concluded that the individuals died during a violent attack on the settlement dated to circa 5000 BCE, i.e., the final phase of the LBK. Those killed appear not to have undergone any mortuary ritual but were instead left unburied and were thus exposed to circumstances that resulted in a variety of taphonomic alterations, e.g., carnivore activity in the form of biting [57]. Based on these findings, and the fact that artifacts diagnostic of subsequent cultural developments seem to be absent, it was assumed that the site was abandoned.

As archaeological excavations continued through 2005, additional human skeletal remains of unburied bodies were recovered from the outer trench (ditch II) of the settlement, so this sample currently includes 130 individuals (including the 67 already known). Among them, 58 children and adolescents, corresponding to a proportion of about 43% (Infans I (individuals within 0-6 years): 29 children; Infans II (individuals within 6-12 years): 17 children; adolescents: 12 individuals). A total of 74 individuals were inferred to have reached adulthood, corresponding to a proportion of 57% (32 under 40 years of age, 42 over 40 years of age), with a clear preponderance of males (44 males and 24 females). The proportion of young (up to 40 years old) women is lower than that of older (over 40 years old) women and seems to confirm the finding of a "deficit of young females" derived from the smaller sample [57]. During the continuation of the bioanthropological and paleopathological investigations of the human skeletal remains, the finds recovered at the settlement area of Schletz, which have been excluded from analysis and remained unpublished, are due to be formally published in a study by Pieler and Teschler-Nicola (2023, in press), of which the present section is a summary. In addition to the victims of the violent conflict, two further clusters of human remains were discovered. Among them are 17 regular, “typical” inhumations in grave pits located around the houses or in a sector of trench II (which had already been filled at the time of the massacre or in abandoned settlement/dwelling pits). When comparing the age and sex profile of the 130 massacre victims and the individuals exhumed at "regular" burials, a significant difference is evident. Male individuals dominate the group of massacre victims located in ditch II, while females and children predominate in the "typical" settlement burials. This predominance of children and women in settlement inhumations has been observed at other Linear Pottery sites.

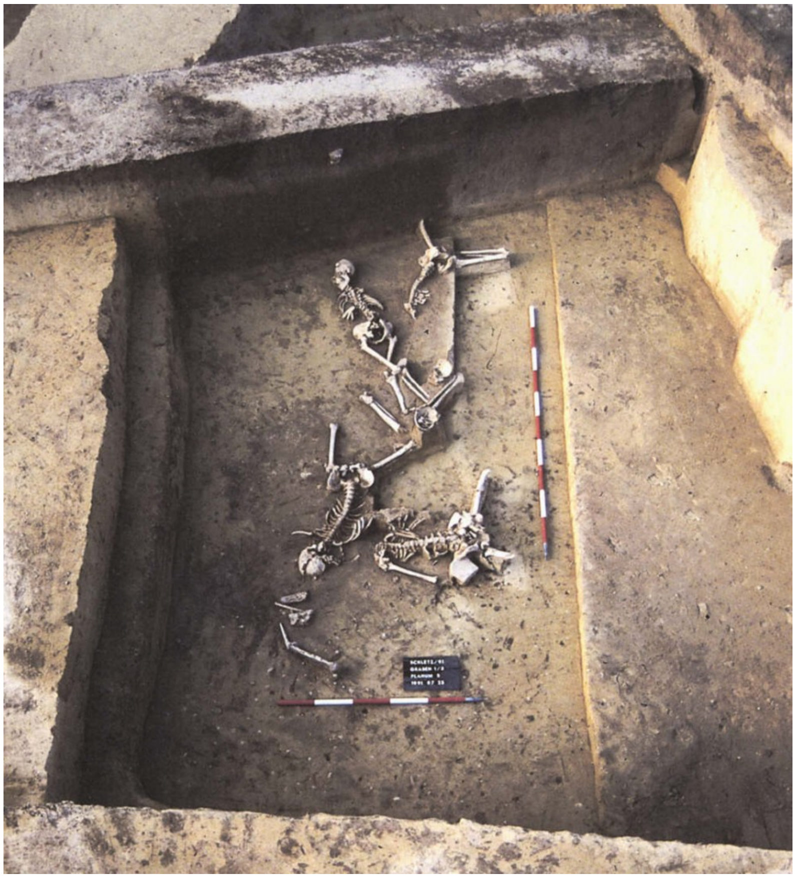

**Supplementary Figure 20:** Skeletal remains of five males in the vicinity of a bridgework *in situ* at the base of the trench of the Asparn Schletz settlement. massacre. These individuals are related to the violent event. Image from MAMUZ with rights for publication.

Another cluster of human skeletal finds in the settlement area is formed by isolated human skeletal elements or small bone fragments, often mixed with those of animals; they were found throughout the area, most frequently in settlement pits, but also in ditches I and II and in the LBK-era well.

To uncover possible diachronic trends in the treatment of the deceased, a few samples were also taken from the inhumations for radiocarbon dating. Preliminary results indicate that some of the "typical" burials are chronologically older (ca. 5,300 BC) than the victims of the massacre recovered from ditches II (ca. 5,000 BC). However, a few dates overlap, and it is not possible yet to establish a clear dating sequence, an undertaking that would need a much larger number of radiocarbon-dated individuals. One internment yielded a slightly younger 14C date, which may indicate at least occasional use of the site after the violent assault.

Different appearances of human remains from Schletz, with a juxtaposition of apparently carelessly deposited, unburied dead and buried in the rite of the time, as well as of scattered individual bones, testify to different practices that existed within the LBK community. The complex situation of the features, especially the ditches, makes it possible here, as at hardly any other site, to investigate the temporal and spatial sequence and relations of the individual phenomena and to shed light on the possible supraregional connections of conflicts because of assumed socio-economic (climatic changes, scarcity of resources, population increase) or socio-cultural changes towards the end of the Linear Pottery period.

We concluded with a note on terminology for the burials analyzed from the Asparn Schletz sites. As mentioned above, the site was systematically investigated over a more than thirty-year period from 1983 and 2005 in annual archaeological excavation campaigns that were consistently carried out in July/August with the help of annually changing student groups and different excavation directors under the responsibility of the (then) Lower Austrian Provincial Museum and the direction of Helmut Windl. Based on these factors and the varying levels of training of the participants, the protocol records and photographic documentation also vary in their form; in addition, we have also to mention that the complexity of this site with many finds of unburied individuals at the base of a ditch (thus, deviating from "normal" burials that were normally characterized and identifiable by “grave numbers”) and the multitude of isolated fragmentary bones which were often not immediately identified as "human remains" (thus, became first included in the animal bone assemblages) were only recognized as such in the course of the current anthropological investigations. This complexity is expressed in the different naming of the human skeletal finds, which were handed over in different portions and at different times for anthropological studies: ind + no. (e.g., ind3) = individual number, allocated during the first anthropological investigations in the 1990s; year + no. (e.g., 2000/1 = year of excavation and individual number used for further findings); FN = finding number either for an inhumation burial or for isolated skeletal elements; names (e.g., "Berbel", “Herbert” ) were allocated by the members of the excavation team; we abstained here from this practice and use the FN.

- **I30418**: ind. 4 1991, FN 3340,5050 BCE (6055+-35)
- **I24884**: 93/1,1993 FN 4520,5205-4847 calBCE (6075±35 BP, VERA-2012)
- **I24885**: 93/18-2,1993 FN 4202,5215-5008 calBCE (6175±35 BP, VERA-2007)
- **I24891**: 93/7-1, FN 4223/4224/4381/433, 5203-4784 calBCE (6025±55 BP, ETH-14373)
- **I24898**: 97/2, FN 5838, 5000 BCE
- **I24899**: 97/4, FN 5839, 5215-5016 calBCE (6175±30 BP, VERA-2737)
- **I27771**: 93/10, FN 4269, 5000 BCE
- **I27772**: 93/11, FN 4264/25310, 5308-5058 calBCE (6235±40 BP, VERA-2020)
- **I27773**: 93/15, FN 4472, 5296-5047 calBCE (6205±30 BP, VERA-2738)
- **I27774**: 93/16, FN 4476, 5000 BCE
- **I27776**: 93/17, FN 4520, 5207-4945 calBCE (6113±23) [R_combine: (6130±35 BP, VERA-2010); (6100±30 BP, VERA-2011)]
- **I27778**: 93/19-2, FN 4503, 5312-5209 calBCE (6268±23 BP) [R_combine: (6254±31 ,MAMS-42229); (6258±31 ,MAMS-42232)]

- **I27780**: 96/4,  FN 5185, 5000 BCE
- **I25323**,96/6, FN 5081, 5000 BCE
- **I27783**: 98/2, FN 6567, 5000 BCE
- **I27784**: 98/4, FN 6449, 5000 BCE
- **I27787_d**: FN, 10640, 5000 BCE
- **I27788**: FN 11057, 5000 BCE
- **I25334**: FN 12566, 5000 BCE
- **I25336**:  FN 12626, 5000 BCE
- **I27793**: FN 12671, 5000 BCE
- **I27794**: FN 209,5000 BCE
- **I27796**: ind. 52, FN 763, 5000 BCE
- **I27805**: FN 14560, 5000 BCE
- **I24268**: FN 11051, 5000 BCE
- **I24270**: FN 12549, 5000 BCE
- **I24271**: FN 12670, 5000 BCE
- **I24275**: FN 265, 5000 BCE

- **I24281**: ind. 41, FN 670, 5209-4951 calBCE (6125±35 BP, VERA-2014)
- **I24282**: ind. 44, FN 646, 5000 BCE
- **I24283**: ind. 50, FN 601, 5000 BCE
- **I24285**: FN 1449 ind. 25, 5000 BCE
- **I30413**: ind 6, 5000 BCE
- **I24289**: ind. 51, FN 2865, 5000 BCE
- **I24886**: 93/13, FN 4471, 5000 BCE
- **I24887**: 93/14, FN 4473, 5000 BCE
- **I24888**: 93/19-1, FN 4503, 5311 BCE (6254±31/MAMS42229)
- **I24889**: 93/20, FN 4529, 5313 (6258±31/MAMS42232)
- **I24890**: 93/25, FN 4444, 5000 BCE
- **I24892**: 93/4, 5197-4844 calBCE (6055±35 BP, VERA-2009)
- **I24276**: Fn. 278, 5000 BCE
- **I30433**: I24277 , ind. 56, 5000 BCE
- **I24279**: ind. 63 FN 286, 5000 BCE

- **I30431**: I24280 ,FN 162, 5000 BCE
- **I24893**: 93/5, FN 4333, 5211-4995 calBCE (6145±35 BP, VERA-2008)
- **I24894**: 93/6, FN 4456, 5000 BCE
- **I24895**: 93/7-2, FN 4223, 5000 BCE
- **I24896**: 93/9, FN 4451, 5000 BCE
- **I24897**: 95/1, FN4694, 5000 BCE
- **I24900**: 97/3, FN 5613, 5000 BCE
- **I24901**: 97/7, FN 5959, 5000 BCE
- **I24902**: 98/1, FN 6316, 5000 BCE
- **I24903**: 99/1, FN 7899, 5000 BCE
- **I24905**: 99/4, FN 8264, 5000 BCE
- **I30421**: 96/3, FN 5184, 5070 BCE(6075±35)
- **I30423**: ind. 5, 5000 BCE
- **I30425**: ind. 7a, 5000 BCE
- **I30428**: 96/2, FN 5076, 5000 BCE
- **I30430**,ind. 1, FN 3342, 5000 BCE

- **I27800**:  ind. 7, 5000 BCE
- **I27785**: 99/2, FN 8053, 5000 BCE
- **I30434**: ind. 47, 5000 BCE
- **I30414**: FN 53, 5000 BCE
- **I24907**:  FN 10821, 5100 BCE
- **I25349**: FN 010351, 5000 BCE
- **I24286_d**: ind. 28 , FN 1465, 5000 BCE
- **I24906**: P8467, FN 10806,5000 BCE
- **I24269**: FN 11660, 5000 BCE
- **I24278**: ind. 57 , FN 249, 5000 BCE
- **I25347**: Graben 3, FN 3491, 5000 BCE
- **I24272**: FN 14143, 5000 BCE
- **I24904**: FN 8328, 5000 BCE
- **I30411**: FN 10343_Siedlung, Berbel ,5000 BCE
- **I24015**: FN 9872_burial , Heli 1, 5214 BCE (6174±25/MAMS38866)

- **I24016**:  FN 9366_burial 15, Silvia , 5033-4847 calBCE (6050±25 BP, MAMS-38865)
- **I24017**: FN 9230, Traude 1 ,5100-4900 BCE
- **I24018**: FN 9230, Traude 2, 5186 BCE (6219±31/MAMS38863)
- **I24021**: FN 9872 ,Heli 3, 5214-5040 calBCE (6174±25 BP, MAMS-38866)
- **I24022**: FN 9872 ,Heli 2, 5214-5040 calBCE (6174±25 BP, MAMS-38866)
- **I24023**: FN 11449_burial 17, Edda, 5100-4900 BCE
- **I24024**: FN 11676, Damia, 5300-5046 calBCE (6210±35 BP, VERA-2198)
- **I24025**: FN 347 , 5302-5041 calBCE (6210±40 BP, VERA-2016)
- **I24026**: FN 14300, Grögar, 5186-5120 (6207±25/MAMS38867)
- **I24027**: FN 13627, Herbert ,5100-4900 BCE
- **I24028**: FN 11803, Daniela, 5212- 5240 (6165±35/VERA2441)

##### 4.29 Nitra (Slovakia)

**Authors:** Daniela Hofmann, Penny Bickle.

Located in the Nitra river valley in western Slovakia, the cemetery lies just where the foothills of the Carpathian mountains begin to stretch eastwards. Like many Linearbandkeramik (LBK) sites, it is located on loess soils, which are found intermittently as the higher ground becomes the Danubian plain to the south and west [58], 5). The site came to light in the course of rescue excavations in advance of the construction of a potato storage building in the south of the modern town of Nitra, in an area named Horné Krškany, about 250–300m from the river bank [59], 231). In 1964 and 1965, Pavúk [58] carried out excavations at the site, identifying 76 graves, two of which were empty, with a few graves destroyed by the initial building works [60], 137). The excavations opened up two parallel trenches, covering an area of 50m by 15m which seems to have included most of the graves that had been preserved, as further test pitting to the north-east and southwest did not uncover more graves [58]. Today 74 individuals are known, with a further 8 cremations, all of which appear to date to the LBK based on both ceramic styles [58], 32) and the radiocarbon dating of 12 graves [61],[58], 84). Assessment of the ceramics accompanying the burials suggested that the cemetery was used over two to three centuries. In terms of Pavúk's chronology, this is from the LBK phase II to the Želiezovce phase, or from the second expansion of the LBK, usually dated to about 5300 cal BC, to its end around 5000 cal BC. A Bayesian model of the 12 radiocarbon dates from the cemetery falls roughly in line with the suggestion of Pavúk and estimates that the cemetery started in *5370–5220 cal BCE (95.4% probable)* or *5320–5230 cal BCE (68.2% probable)*. The end of activity at the Nitra cemetery is estimated to have occurred in *5210–4980 cal BCE (95.4% probable)* or *5210–5090 cal BCE (68.2% probable)*. The duration of burial at the site is therefore estimated to have lasted between *20–360 years (95.4% probable)* or *30–220 years (68.2% probable)* [61]. Thus, it seems possible that the cemetery received burials for anything from a generation to several centuries.

The burials themselves cluster in the northeastern part of the trenches, but beyond this, and in contrast to other cemeteries (e.g. Vedrovice, Cz, and Aiterhofen, Germany), other groupings or sub-divisions are not obvious (see Figure 3). Pavúk, [58] suggested that the graves may be arranged in lines rather than groupings. Some 22 burials appear associated through nine sets of intercutting grave pits (in pairs or clusters of three burials), which is a feature far rarer at other cemeteries (but also suggested for Elsloo [62]. Burials are largely found in single inhumations, except for a triple burial (individuals 48, 49, and 50) of an adult female with two children, the latter of whom had received blows to the head which most likely caused their deaths [5], 148). Most graves were oriented along a southeast-to-northwest axis, with the head to the southeast, though burials could fall between east–west, and south–north [58]. As another counter-point to other LBK cemeteries, no burials are found in the antipodal orientations. Where body position can be determined, the deceased was placed in a crouched position mostly on their left-hand sides [58]), though a few were found with the upper part of their body on the front or back. Only two burials (43 and 71) were found on their right-hand sides. Overall, there is more similarity in grave orientation and body position than at other LBK cemeteries.

The grave goods accompanying the burials are typical for LBK cemeteries and comprise pottery, polished stone tools, chipped stone implements, imported *Spondylus* shell either as beads or as "belt buckles'', worked bone, ochre coloring and, in one instance, pieces of graphite (burial 5) [58]. About a third of burials were accompanied by no grave goods at all, which is comparable with other LBK cemeteries ([5], 142). Unusually, one burial is accompanied by seven perforated human and dog or fox teeth (burial 19) ([58], 11). Pottery decorated in a style that mixed local patterns with those more closely associated with the Alföld Linear Pottery group (located in north-eastern Hungry) was found in grave 17, suggesting wider connections ([58], 84). This particular connection is also suggested by a pot in grave 2 ([58], 84). Overall, older men, and to a lesser extent older women, appeared to be accompanied by the most numerous and diverse grave good assemblages, which led Pavúk [58], 72) to suggest that Nitra was a gerontocratic society, with status increasing for some as they aged. This trend for older individuals to have the most grave goods also holds true when using up-to-date methodologies for estimating age and sex from skeletal data[2].

The osteological collection from Nitra has been subject to several different studies. Linda Fibiger, for the LBK lifeways project, carried out an extensive assessment. The 75 individuals studied by Fibiger consisted of 27 adult females, 18 adult males, 4 unsexed adults, 6 adolescents, 16 juveniles, and 4 infants [5], 143). This is probably not representative of a living population and under-represents the likely rate of infant mortality. Dočkalová and Čižmář [63,64] have demonstrated that in this region, higher rates of non-adults were buried in settlements, suggesting some deliberate selection for burial in cemeteries based on age. Among the adult burials, females have slightly higher representation in the young adult category (18-25 years at death), likely representing death in pregnancy or childbirth;

otherwise, males and females are found in roughly equal proportions as they aged, with the highest numbers of individuals falling into the mid-adult category ([5], 144). Evidence for metabolic and infectious conditions, as well as generalised stress markers like enamel hypoplasias, were equal between the sexes, suggesting that periods of stress likely affected the whole population ([5], 146-149). Overall, at least a fifth of the population had signs of periosteal changes and infection. Alongside the two children noted above, a young adult male (72) and a young adult female (1) also showed traces of skull trauma ([5], 148).

Sex-based differences were suggested based on a number of lines of evidence. Higher rates of dental caries in women, coupled with higher δ15N values ([5], 146, 150) in males, indicate a degree of dietary differences, but these were not detectable by dental microwear [65]. In particular, males buried with polished stone axes had higher nitrogen values than the site mean, while all burials accompanied by imported *Spondylus* shell (independently of the sex of the deceased) had higher δ13C values on average than the rest of the population ([5], 151). Stronger associations between the sex of the deceased and strontium isotope ratios were found. Women were found to have a much wider range of strontium isotope values, and all 6 of the individuals falling above the upper limit of the loess strontium range were women ([5], 152). A sexed division of labor has since also been identified from use-wear analysis of the stone tools accompanying male and female burials. Males were associated with tools that had been used in woodworking, animal butchery and/or interpersonal violence, and harvesting, and women with tools for hide working. Occlusal grooves on teeth also indicated that women engaged in sinew or plant fiber processing more often than males ([66]; [5], 146).

- **I17346**: 2946, grave 7, 5500-4500 BCE
- **I17539**: 2939, grave 40, 5500-4500 BCE
- **I18106**: 50, grave 8, 5500-4500 BCE
- **I11866**: 209, grave 56, 5500-4500 BCE

- **I11872**: 216, grave 26, 5500-4500 BCE
- **I14177**:  1530, grave 675300-5000 BCE
- **I14178**: 1514, grave 50, 5300-5000 BCE
- **I14179**: 1490, grave 30, 5300-5000 BCE
- **I14180**: 1525, grave 62, 5300-5000 BCE
- **I14181**: 1537, grave 74, 5300-5000 BCE
- **I14182**: 1518, grave 55, 5300-5000 BCE
- **I14183**: 73, grave 73, 5300-5000 BCE
- **I14599**:1483 grave 22, 5300-5000 BCE
- **I14600**:1466 grave 45, 300-5000 BCE
- **I16008**:1474 grave 14, 5300-5000 BCE
- **I16010**:1534, grave 71, 5300-5000 BCE
- **I16011:** 54, grave 54, 5300-5000 BCE
- **I16012**:1489, grave 29, 5300-5000 BCE
- **I16013**: 1508, grave 44 , 5300-5000 BCE

- **I16014**: 1503, grave 39, 5300-5000 BCE
- **I16015**: 1529, grave 66, 5300-5000 BCE
- **I16016**: 1522, grave 59, 5300-5000 BCE
- **I16239**: 1463, grave 1, 5300-5000 BCE
- **I16240**: 1511, grave 47, 5300-5000 BCE
- **I16242**: 1507, grave 42, 5300-5000 BCE
- **I16245**: 1479, grave 19, 5300-5000 BCE
- **I17339**: 1484, grave 24, 5300-5000 BCE
- **I17340**: 1493, grave 33, 5300-5000 BCE
- **I17341**: 1531, grave 68, 5300-5000 BCE
- **I17343**: 1539, grave 76, 5300-5000 BCE
- **I17344**: 1464, grave 3, 5300-5000 BCE
- **I17345**: 1468, grave 5, 5300-5000 BCE
- **I17545**: 1476, grave 16, 5300-5000 BCE
- **I18093**: 1469, grave 6, 5300-5000 BCE
- **I18094**: 1478, grave 17, 5300-5000 BCE

- **I18097**: 1485, grave 25, 5300-5000 BCE
- **I18111**: 8541, grave 21, 5300-5000 BCE
- **I25176**: 1516, grave 53, 5300-5000 BCE
- **I18143**: 5628, grave 8, 5300-5000 BCE
- **I18144**: 8539, grave 36, 5300-5000 BCE
- **I25201**: 1487,  grave 27,  5300-5000 BCE
- **I18091**: 1465, grave 2, 5300-5000 BCE
- **I16009**: 1481, grave 23, 5300-5000 BCE
- **I16246**: 156078, grave 9, 5300-5000 BCE
- **I16007**: 1513, grave 49, 5300-5000 BCE
- **I16241**: 1475, grave 15, 5300-5000 BCE
- **I17538**: 1501,  grave 38, 5300-5000 BCE
- **I11873**: 1512, grave 48, 5300-5000 BCE
- **I25175**: 1515, grave 52, 5300-5000 BCE
- **I18105**:1533, grave 70, 5300-5000 BCE

##### 4.30 Bajč, site Medzi Kanálmi, district Nové Zámky (Slovakia)

**Authors:** Ivan Cheben, Matej Ruttkay

The site of Bajč - Mezi kanmi is located in south-western Slovakia in the district of Nové Zámky, on the left bank of the Žitava river. Maps from the 18th and 19th centuries show that it was originally an island in the flood plain of the Žitava river.

Archaeological research was carried out here in 1987-1992 (research leaders M. Ruttkay, I. Cheben). On an area of approximately 35,000 $m^2$, a large settlement area from several periods was investigated - Neolithic, Chalcolithic, Hallstatt, Late Iron Age (La Tene), and settlements and burial sites from the 7th - 18th centuries.

The site was most intensively settled in the Neolithic - LBK, at the end of the period of the cultural development of the Želiezovce group. Archaeological structures from this period were found only on the northern, elevated part of the hill. It is possible that the southern part was waterlogged and unsuitable for settlement at that time, as the river level was higher at that time.

Hundreds of settlement features (568 settlements pits and 205 post-holes) were excavated from this period and four graves with skeletons were uncovered. It has not been possible to clearly identify the remains of the characteristic houses of this period. Their presence could be partly indicated by groups of post-holes. Based on the evaluation of the pottery, it was determined that the objects and graves chronologically belong to Stage III of the Želiezovce Group. The individual graves with skeletons were situated in the southern area of the investigated area but did not form any coherent group in terms of their layout. These were separately deposited remains in grave pits, and the grave furnishings showed great variation. In addition to the four graves with skeletals, finds of human bones were found in several settlement features. In terms of anthropological determination, long bones (femur) and parts of the skull appeared to be more prominent. Similar to the graves, with skeletons, in the settlement pits the human skeletal parts do not show any grouping or form any structure. In all probability, both the graves and the objects with human bones were disparately distributed over the settlement area. This fact may indicate the different relationship of the population of the time to the human remains.

The analysed grave 1 had an oval pit with dimensions 150x180cm. A skeleton was uncovered at a depth of 55cm. The dead was placed on its left side with its lower limbs curled up. Orientation east-west, head to the east, and face south. The grave had an exceptional assemblage for that time: 8 clay vessels, a radiolarite blade, a stone mace head, a spondylus bracelet, a stone shoe-adze, a necklace of stones and shells, a stone mat, an antler pendant, a bone socket, the remains of a goat or sheep, and other finds. It was probably a woman with an estimated height of 158.3 cm [67–70]

- **I24296:** Objekt 479, 5500-4500 BCE
- **I24869:** Objekt 550, 5500-4500 BCE
- **I24874:** Objekt 528, 5500-4500 BCE
- **I24871**: Objekt 571 , 5500-4500 BCE

##### 4.31 Patince (Slovakia)

**Author:** Matej Ruttkay

- **I24872**: Objekt 74, 5500-4500 BCE
- **I24865**: Objekt 163, 5500-4500 BCE

##### 4.33: Jelšovce (Slovakia)

**Authors:** Matej Ruttkay, Jozef Bátora

During extensive rescue research at the polycultural site in Jelšovce, in the district of Nitra in southwestern Slovakia, it was possible to examine a part of the Neolithic settlement, which was represented by the Želiezovce group. Among the uncovered houses, house 615 stood out from the others, in the foundation gutter of which were found the skeletons of two adult women aged 40-50 years old. These skeletons were emarked as grave 615A and 615 B.

The burial of individuals directly in the foundation gutter has not yet been well documented in the Central European area from the early Neolithic period ([71], 18-20). The find from Jelšovce is therefore of particular importance.

Because traces of injuries were found on the skulls of both women buried in the foundation gutter of the house [72], a possibility is that both women died violent deaths as part of ritual ceremonies held in connection with the start or end of the construction of a new house [73].

- **I24295**: Jelšovce 615b, 5500-4500 BCE

  Skeleton A was located in the lower narrowed part of the western half of the foundation gutter at a depth of 43 cm, oriented in the NE - SW direction. Due to the narrow space (the width of the trough was 23-30 cm), the skeleton was forced into the gutter. There were 3 vessels at a depth of 15 cm above the skull of the skeleton.

- **I24868**: Jelšovce 615a/87, 5500-4500 BCE

  Skeleton B was located in the eastern part of the gutter at a depth of 30-36 cm. It was placed on the right side, oriented in the WSW - ENE direction. The view of the facial part of the skull was directed to the SSE. No grave goods were found by the skeleton.

**4.34: Nitra Mlynárce**

**Authors:** Matej Ruttkay, Jaroslava Ruttkayová

The burial site was found in 1951 during construction activities, when many graves had already been destroyed. The graves were located in several places over time. During 1951-53 at least 16 graves were examined on the right bank terrace of the Nitra River, five of them concentrated in an area of 8.5x3.5m (graves 1-5/51), another group consisted of 10 graves (graves 1-10/52), and an additional grave (1/53) was found in 1953. The orientation of the graves was inconsistent. The dead were most often placed in a crouched position on the left flank, with only grave 1 on the right flank. In one of the best-preserved graves, the dead lay on their backs with their arms strongly bent at the elbows and their hands at their shoulders, their face tilted towards the south and their lower limbs bent towards the south. Graves 1/52 and 2/52 were probably in superposition. Traces of red dye were found on the skeletons. The inventory of the graves consisted of pottery, an amphibolite axe and hoof wedges, shell and marble beads, bone needles, and chipped stone industry. Evidence of LBK settlement was found in the immediate vicinity [74].

- **I7893**:  NTMY_3/52, 5300-5000 BCE

- **I7894**: NTMY_4/52, 5300-5000 BCE

- **I7895**: NTMY_5/52, 5300-5000 BCE

- **I7896**: NTMY_7/52, 5300-5000 BCE

- **I7892**: NTMY_2/52, 5300-5000 BCE

#### Section 5: Classification of individuals with genetic methods

**Authors**: Pere Gelabert

From the PCA two outliers were obvious, both labeled as ALPC. Individual (I1877) is almost entirely of Early European Farmer (EEF) ancestry (95%), and one Starčevo individual has substantial WHG ancestry (I1876) (18%) [75]. Nevertheless, we used qpWave to systematically determine the outliers (Supplementary Figure 1). Those that show consistent deviation from the general cluster are labeled EXC in Supplementary Table 1.

Previous ancient DNA studies of the Körös and Starčevo archaeological cultures [75] documented that members of these communities varied in their degree of admixture with local hunter-gatherers and thus were far from genetically homogeneous. Given small sample sizes, however, it was impossible to identify cultural features associated with those people more likely to have experienced this admixture. Pooling the individuals from each culture, and excluding those with elevated WHG ancestry (excluding the two full WHG individuals I1507 and I4971) we found that the statistic $f_4$(Turkey N, WHG; Körös, Starčevo) has a value of 0.0002 (Z=1.89) suggesting no differential presence of WHG between Körös and Starčevo. To further explore these relationships, we applied qpAdm to model the Körös genomes using Balkan_N and WHGA as possible sources after the removal of the Körös and Starčevo individuals from the Balkan_N set. Both populations can be modeled with Balkan_N genomes ($p$>0.01). Four Körös individuals have higher than average WHG values: I18642 has 11%, I17931 has 11%, I2373 has 9%, and I4971 is an unadmixed WHG. Additionally, we report two Starčevo individuals with higher-than-average WHG values: I4918 with 7% and I6699 with 19% (Supplementary Table 3).

We tested for differences in the WHG source populations for the different European farmer groups: from a far western European context (Loschbour from Luxembourg) or a central European context (KO1: an individual with WHG ancestry found at a Koros site), using the statistic $f_4$(Mbuti, X; Loschbour, KO1). This statistic is sensitive to differences in rates of allele sharing of a test population with the two important sources for WHG ancestry in

Europe. We performed these analyses at a populational level excluding the individuals with elevated WHG ancestry, labeled HGEXC in Supplementary Table 1. Neither result was statistically significantly asymmetric at the |Z|>3 significance level. We then used the statistic $f_4$(Mbuti, X; Körös, Starčevo) to test the previously suggested hypothesis [76] that the ALPC derived ancestry from Körös while the LBK derived ancestry Starčevo (Supplementary Table 4). The results do not reveal any significant asymmetries, in the sense that none of the comparisons yield a Z-score > |3|.

To obtain further insight into the WHG ancestry, we generated diploid genotype calls for the admixed ALPC individuals using the software GLIMPSE 2 [77], and performed a local ancestry-segmentation analysis using RFMix 2.03 [78], aiming to detect segments that are likely to derive from WHG. We also explored if $f_4$ tests on the WHG segments would give more resolution regarding the origin of the WHG in the ALPC, but this was not so (Supplementary Table 4).

### Section 6: Population size trajectory inference

**Authors**: Romain Fournier, Pier Francesco Palamara

**Methods**

HapNe is a method that can estimated demographic size changes in aDNA data over the few thousand years before an individual lived [79]. HapNe assumes that the individual is drawn independently from a single panmictic population. This hypothesis is likely to be inaccurate when individuals from an archaeological site are studied, as relatives are likely to be located in the same burial site. To mitigate this, we only retained a single individual with the highest coverage within each identified family. However, the presence of undetected families or other structures in the remaining samples might introduce linkage disequilibrium (LD) that is not accounted for in the HapNe panmictic population model. To mitigate this potential issue, we performed a filtering step based on the genetic relationship matrix in each group. We ran HapNe's cross-chromosome LD test to compute an approximate p-value for the hypothesis that there is no underlying structure in the input samples after filtering using these two steps. This hypothesis was not rejected for the Schletz massacre samples (approximate p-value = 6.7%) and Nitra (approximate p-value = 40.6%). However, the hypothesis was rejected for the individuals of Polgar (approximate p-value < 0.0001), and thus we removed this site from our analyses of inference of population size changes over time.

We ran HapNe using default parameters and in "fixed prior" mode, which is recommended when comparing groups of different sample sizes or coverage. We ran HapNe without correcting for time heterogeneity for the Schletz samples, as they originate from the same time point, and assumed a uniform distribution between 5300-5000 BCE for the samples from Nitra. Finally, generations were converted to years using a factor of 29.1 years per generation [79]. After selecting the individual with the highest coverage for each identified family, we used plink 2.0 [80] to compute the genetic relationship matrix (GRM). We then projected the individuals onto the first two principal components of the matrix, shown in

Supplementary Figure 21. We removed all visual outliers (in red in Supplementary Figure 21) before testing for the presence of population structure.

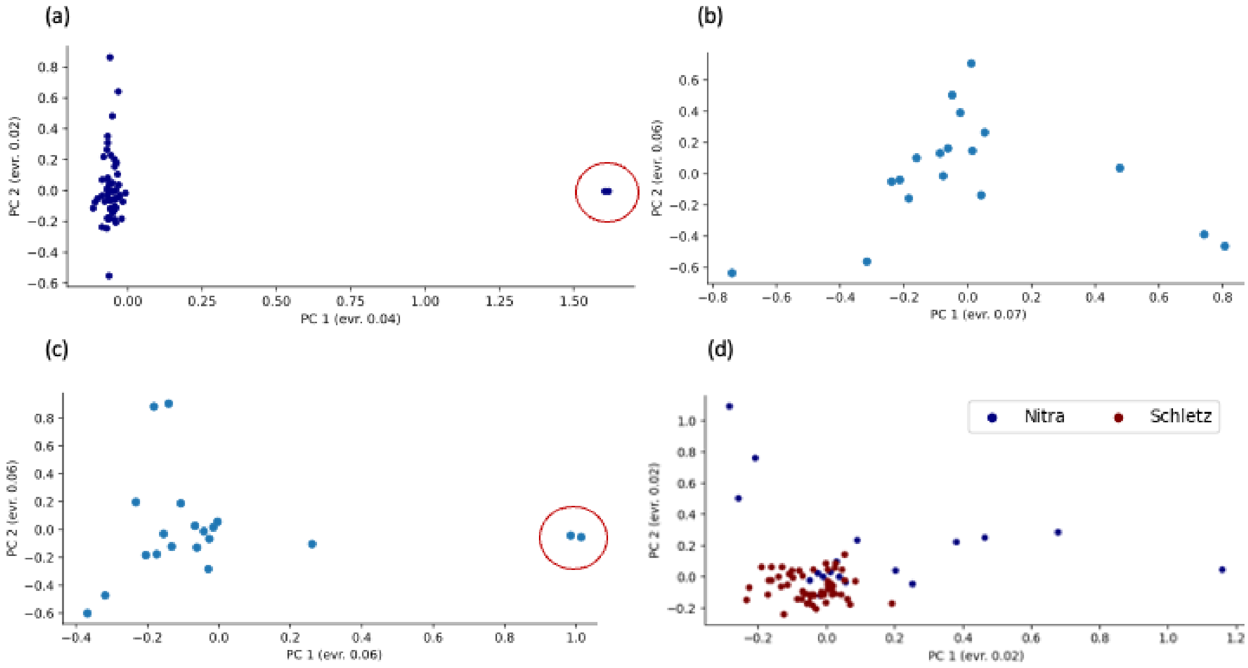

**Supplementary Figure 21: Projection of unrelated individuals of different groups onto the first two principal components of their GRM.** (a) Schletz massacre. (b) Nitra. (c) Polgar. (d) Schletz (s=56) and Nitra (s=18). Individuals filtered out in (a-c) are circled

#### Section 7: Selection scans in the diploid data

**Authors:** Xin Huang, Martin Kuhlwilm

##### Filtering data

After obtaining the imputed dataset, individuals with imputation quality score less than 0.8 were removed. We then extracted biallelic single nucleotide polymorphisms (SNPs) with moderate quality or high-quality imputation quality. Only SNPs with ancestral alleles in the 1000 Genomes Project [81] were used for further analysis. The ancestral allele of each SNP was required to match either the reference allele or the alternative allele. The final SNP dataset was annotated with three databases—RefSeq, dbNFSP version 4.2c and dbSNP version 150 with left-normalization—using the human reference genome hg19 coordinates by ANNOVAR version 2022Oct05 [82–85]. In total, 62 diploid imputed genomes from the ALPc population and 137 diploid imputed genomes from the LBK population were used for the studies of natural selection.

##### Testing for evidence of selective sweeps in the thousands of years before the early farmers we analyzed lived

To detect signatures of positive selection in the data, the iHS and nSL scores were estimated with selscan 2.0 [86]. Only SNPs with a minor allele frequency (MAF) larger than 0.05 were used as cores when calculating these scores. Other parameters were set to the default values in selscan 2.0. Unnormalized iHS and nSL scores were normalized using the norm program from selscan 2.0 with the default settings. To detect signals of long-term balancing selection in the data, the B1 scores were estimated with unfolded allele frequencies by BetaScan [87]. Only SNPs with MAFs larger than 0.05 were used during the whole genome scan. SNPs were further removed if they were in the regions defined by the RepeatMasker table, simple repeats table, and segmental duplication table from the UCSC Table Browser with hg19 coordinates (last accessed December 2022). SNPs were also removed if they had p-values less than $10^{-3}$ from exact tests for Hardy-Weinberg equilibrium in each population by PLINK 1.09 [88,89]. Following Siewert and Voight (2017), only SNPs with a minor allele frequency larger than 0.15 were used as cores when calculating the B1 scores. Other parameters were set to the default values in BetaScan.

**Results and Discussion**

We tested for signatures of positive selection with both the iHS and nSL scores (Figure 6A–6D), because the iHS score may only perform well in detecting hard selective sweeps, while the nSL score is expected to be sensitive to soft as well as hard selective sweeps [90]. We chose those SNPs with either absolute normalized iHS or nSL scores in the top 0.05% as candidates (Supplementary Table 11). Based on both the iHS and nSL scores, fifty genes harbor signatures of positive selection in both the ALPc and LBK populations (Supplementary Table 11). Several candidate genes are plausibly associated with human pigmentation. For example, the *MLPH* gene encodes melanophilin, which may affect melanosome transport [91]. A previous study also suggested this gene had strong XP-EHH signals in non-African populations [92] who lived approximately 7000 years after the individuals we analyze here. The *PRKCH* gene encodes the PKCη protein in melanocytes and may participate in the protein kinase C-dependent pathway to regulate melanogenesis [93]. The *PTPRN2* gene had a higher level of expression in lightly pigmented melanocytes than darkly pigmented melanocytes, a phenotype that is similar to that of human pigmentation gene *SLC45A2* [94]. The expression levels of *CDH12*, *ERBB4*, and *MACROD2* was significantly changed in hyperpigmented skin using meta-analysis, indicating they may affect human pigmentation [95].

We also tested for evidence of long-term balancing selection with the B1 score from BetaScan [87]. We chose those SNPs with B1 scores in the top 0.05% as candidates (Supplementary Table 11). Similar to previous studies using different approaches to detect long-term balancing selection with modern human populations [96,97] , the strongest signals are from the HLA region on human chromosome 6 (Figure 6, Supplementary Figure 7). Twenty-nine genes harbor signatures of long-term balancing selection in both the ALPc and LBK populations (Supplementary Table 12). Most of them were reported as significant outliers in a previous study using genomes from modern European populations: GBR and TSI [97], indicating long-term balancing selection may persist at these genes.

We explored the possibility that some of the signals of selection would be associated with differential proportions of WHG ancestry and thus we computed the average WHG ancestry in the sections of the genome where the SNPs were located (Supplementary Table 11) and plotted the average selection scores from candidate SNPs within each gene against the WHG ancestry (Supplementary figure 8). The correlation coefficients provide suggestive but not compelling evidence that non-WHG ancestry contributed more to the top iHS/nSL scores (the best P-value of 0.027 is not significant after Bonferonni correction for four hypotheses tested).

### Section 8: Supplementary Methods

#### Laboratory procedures

We generated powder from the skeletal remains of all individuals that are listed in the Supplementary material. Supplementary Table 1 shows the list and details of the individuals. The powder was produced from the cochlea [98], ossicles [99], or teeth in clean rooms at the University College Dublin or the University of Vienna.

We extracted DNA in dedicated ancient DNA laboratories at Harvard Medical School or the University of Vienna, following published protocols [100,101]. Duble-stranded libraries were prepared from the extracts, using either dual-barcoded double-stranded libraries [102] or dual-indexed single-stranded libraries [103], both treated with uracil-DNA glycosylase (UDG) to reduce the rate of ancient DNA damage [104]. Double-stranded libraries were treated in a modified partial UDG preparation ('half'), leaving a reduced damage signal at both ends (5′ C-to-T, 3′ G-to-A). For some individuals with little success, we made more than one library per extract. The list of the libraries generated in this study is presented in Supplementary Table 2.

The newly produced libraries were captured with un-solution target hybridization to enrich sequences that overlap the mitochondrial genome and about 1.24 million genome-wide SNPs [105–108] ('1240K'). Then, captured libraries were indexed with two seven-base-pair indexing barcodes to the adapters of each double-stranded library. The indexed and pooled libraries were sequenced in an Illumina NextSeq500 instrument with 2 × 76 cycles or an Illumina HiSeqX10 instrument with 2 × 101 cycles and reading the indices with 2 × 7 cycles (double-stranded libraries).

After sequencing, paired-end libraries were merged, Before alignment, we merged paired-end sequences, retaining reads that exhibited no more than one mismatch between the forward and reverse base if the base quality was ≥20, or 3 mismatches if the base quality was <20. A custom toolkit (available at https://github.com/DReichLab/ADNA-Tools) was used for merging and trimming adapters and barcodes. The merged reads were aligned with BWA samse v.0.7.15-r114053 using typical aDNA parameters (-n 0.01, -o 2, and -l 16500) to the reconstructed human mtDNA consensus sequence (RSRS)(Behar et al. 2012) and the human reference genome version hg19. We removed duplicates with Picard MarkDuplicates tool [109]. After this, We trimmed two terminal bases from UDG-half libraries to reduce damage-induced errors.

To discard contaminated samples, we discarded libraries with less than 3% of cytosine-to-thymine substitutions at the end of the sequenced fragments. and point estimates of mitochondrial DNA (mtDNA) contamination below 5% using contamMix v.1.0-1248, and point estimates of X chromosome contamination (in males) below 3%; We also used contamLD to confirm low contamination rates (less than about 6%). The results are presented in Supplementary Table 1. For SNP calling, we randomly sampled an overlapping read with minimum mapping quality of ≥10 and a base quality of ≥20. Individuals with <30,000 covered SNPs were excluded from quantitative analyses. For first-degree relatives, we always excluded the individual with less coverage from the pair in all the population genetics analyses.

**Bioinformatics**

Genetic data were merged with published datasets of Early Neolithic and Mesolithic individuals [75,107,110–114]. We excluded from the analyses all individuals with a 1st degree relative, showing clear signs of human contamination, or with low coverage (less than 30,000 SNPs in the autosomes).

We used qpAdm to classify the individuals of the dataset using the strategy described in [115] running it per single individuals with (Balkan_N and WHGA, Old Steppe) (Sup table) as a source and (Turkey_N, OldAfrica, WHGB, Russia_Afanasievo) as right populations, accepting models fitting with p-values > 0.05. We removed individuals showing Steppe ancestry, and confirmed that these individuals were misassigned chronologically by direct dating. qpAdm was run again, now with Balkan_N and WHGA as sources and Turkey_N, OldAfrica, and WHGB as the right populations. This was used to estimate the proportions of EEF and WHG components on the genomes. After this initial step, we grouped the individuals with similar ancestry profiles.

We used Principal Components Analysis (PCA) with the smartpca package of EIGENSOFT to graphically represent the individuals and the relationship between them and to position the individuals within the main axis of ancestry (WHG and Balkan_N). We used f-statistics to compare the relationship between groups of individuals using admixtools 7.0.2 [116].

DATES 3600 [117] was used to date the admixture events detected in the qpAdm analyzes using Balkan_N and WHGA as source populations.

HapROH from [118] was used for inferring runs of homozygosity on individuals with more than 400,000 SNPs covered in the autosomes. The methodology described in [119] was used for IBD reconstruction from the diploid imputed data.

We ran qpWave grouping all the individuals from the same culture and location. This strategy was used to describe genetic outliers using the strategy described in [115] . The Körös and Starčevo included in Patterson et al 2022 as WHGA (I1507, I4971) or BalkanN (I0174, I1876, I1508, I2794) were used among the source populations and therefore we do not present qpWave or qpAdm results of these. Individuals with a p-value > 0.05 were labelled as outliers. The results are presented in Supplementary Figure 1.

For the local ancestry analyses, we selected a set of 6,237,504 autosomal SNPs present in the imputation panel, having more than a 95% of genotype probability quality. We also selected all the ALPC individuals with an imputation quality score (IQS) > 0.9. A total of 55 out of 77 individuals passed this filter (Table S5). The focus on ALPC aims to understand specific patterns of admixture of this admixed population.

A total of 55 phased genomes were analyzed with RFMix 2.03 [78], setting 14 generations of admixture time and 2 EM interactions, -c 0,2 cM, -G 14, -t 500 and the rest with default parameters. Chromosomes were collapsed and plotted accepting sites with >0.9 probability.

We used the output bed files from RFMix to isolate the WHG ancestry with Plink 1.9 and subtract the WHG fragments from the pseudo-haploid calls. We used these data to perform f-statistics analyses with ADMIXTOOLS 7.0.2.

We calculated the average WHG ancestry of ALPC genomes in chunks of 0.2 cM and added this information to the selection scan results (Supplementary Table 11). We calculated the average ihs/nsl score for the same fragments and plotted this relationship with R in Supplementary Figure 8. To test for associations between selection coefficients and WHG ancestry, we computed the Spearman correlation coefficients.

**Grave Distances Calculation**

The distances between graves in the Nitra and Polgar cemeteries were calculated using ImageJ and R. Pixel coordinates of each grave were recorded in ImageJ from Figure 3. These data were then used to compute the relative distances between each individual. These calculations were performed in R.
